# Topological incongruence between Median-Joining Networks and Bayesian inference phylogenies

**DOI:** 10.1101/2022.10.01.510440

**Authors:** Sungsik Kong, Santiago J. Sánchez-Pacheco, Robert W. Murphy

## Abstract

Inferring phylogenies among intraspecific individuals often yields unresolved relationships (i.e., polytomies). Consequently, methods that compute distance-based abstract networks, like Median-Joining Networks (MJNs), are thought to be more appropriate tools for reconstructing such relationships than traditional trees. MJNs visualize all routes of relationships in the form of cycles, if needed, when traditional approaches cannot resolve them. However, the MJN method is a distance-based phenetic approach that does not involve character transformations and makes no reference to ancestor–descendant relationships. Although philosophical and theoretical arguments challenging the implication that MJNs reflect phylogenetic signal in the traditional sense have been presented elsewhere, an empirical comparison with a character-based approach is needed given the increasing popularity of MJN analysis in evolutionary biology. Here, we use the conservative Approximately Unbiased (AU) test to compare 85 cases of branching patterns of cycle-free MJNs and Bayesian Inference (BI) phylogenies using datasets from 55 empirical studies. By rooting the MJN analyses to provide directionality, analyses find substantial disagreement between computed MJNs and posterior distributions on BI phylogenies. The branching patterns in MJNs and BI phylogenies show significantly different relationships in 37.6% of cases. Among the relationships that were not significantly different, 96.2% show different sets of relationships. Our results indicate that the two methods provide different measures of relatedness in a phylogenetic sense. Finally, our analyses also support previous observations of the statistical hypothesis testing by reconfirming the over-conservativeness of the Shimodaira–Hasegawa test versus the AU test.

## 2 Introduction

Phylogenetic inference involves statistical analysis of the differences between taxa and then represents their evolutionary relationships in a branch-like structure (Felsenstein, 2004). The tree model strictly assumes vertical evolution where a single ancestral taxon splits into two daughter taxa that denotes a speciation event, resulting in a diagram consisting of a set of bifurcations. Such bifurcations often sufficiently describe speciation events but overlook common and important evolutionary histories that also generate novel lineages, like hybridization, due to structural constraints. The ability to infer and illustrate such complex evolutionary phenomena is a critical component for the expansion of evolutionary research (Blair and Ané, 2020, Kong, 2022).

The concept of a phylogenetic network has been proposed as an alternative to the tree model. In brief, a phylogenetic network considers reticulation events in the form of a directed acyclic graph with at least one vertex with in-degree two and out-degree one, representing non-treelike evolutionary scenarios. In practice, no authoritative definition of ‘phylogenetic network’ appears to be available and thus it is often described in a way that the authors happen to be interested in (Linder and Rieseberg, 2004, Gusfield et al., 2007), although Kong et al. (2022) present formal definitions of classes of phylogenetic networks in more mathematical terms.

Networks are largely classified into explicit and abstract networks. Explicit networks describe evolutionary relationships and many efforts to efficiently infer such networks have been made (e.g., PhyloNet (Than et al., 2008, Wen and Nakhleh, 2018); SNaQ (Solís-Lemus and Ané, 2016)). However, current inference methods require high computational capability and they become very expensive even to infer the relationships among two dozen taxa (Hejase and Liu, 2016). In contrast, abstract networks visualize possible routes of relationships of the nucleotide arrangements in the input sequence alignment based on overall similarity. Intriguingly, unrooted abstract networks enjoy increasing popularity in evolutionary studies even though they do not possess evolutionary significance.

The Median-Joining network (MJN) is one of the most popular methods to compute abstract networks. The method clusters sequences on the basis of overall similarity and the lack of a root precludes identification of the direction of evolution (Kong et al., 2016, SánchezPacheco et al., 2020). A rooting option is available when computing MJN, but this procedure is often bypassed in practice. The cyclic structure in MJNs must be distinguished from the reticulation in explicit networks as they do not represent complex evolutionary scenarios but merely illustrate the algorithm’s inability to discern optimal connections (Salzburger et al., 2011). Despite these flaws, the application of MJNs is not waning but rather flourishing with nearly 12,000 citations as of writing, probably because of the fast computation and freely available, simple to use software NETWORK (Bandelt et al., 1999; Fluxus Technology, 2015; available on http://www.fluxus-engineering.com).

Bayesian inference (BI) reconstructs historical ancestor–descendent relationships (Yang and Rannala, 1997, Huelsenbeck and Ronquist, 2001). Sampling the posterior probability in BI methods is not trivial, and computational efficiency is achieved through application of Markov chain Monte Carlo (MCMC) that approximates the posterior distribution of trees (Mau and Newton, 1997, Yang and Rannala, 1997, Mau et al., 1999, Li et al., 2000). However, BI phylogenies may be unsuitable to infer relationships among intraspecific individuals because the produced tree will have poor resolution as BI samples multiple trees that require merging using consensus criteria (Posada and Crandall, 2001). Moreover, setting appropriate prior parameters to conduct BI approaches is not always straightforward and the estimated posterior probability can be excessively liberal when compared to the bootstrap support from a maximum likelihood phylogeny (Suzuki et al., 2002). Nevertheless, BI is one of the most reliable methods available today for phylogenetic tree reconstruction.

Branching patterns among a set of phylogenetic trees with the same ingroup taxa often differ due to random chance, sampling error, and many others factors. Techniques for measuring and testing the significance of topological incongruence are used widely. Incongruence tests that consider character information are more powerful and useful than those that only consider tree shape or topology because they take into account both the topology and its underlying support. For example, the Shimodaira-Hasegawa (SH) test (Shimodaira and Hasegawa, 1999) is a non-parametric bootstrapping method that compares topologies in a likelihood framework. This test can compare multiple topologies considering the null hypothesis that all the trees tested are equally good explanations of the data whereas the alternative hypothesis is that one or several trees are better approximations of the data. Because the SH test can be very conservative in its rejection of the null hypothesis when the number of candidate trees is large, the Approximately Unbiased (AU) test (Shimodaira and Hasegawa, 2001, Shimodaira, 2002) attempts to ameliorate this bias.

Herein, we use non-recombining published empirical genetic datasets to observe the topological discrepancies between MJNs and BI phylogenies estimated from the same sequence alignments. Because the true relationships are unknown, we employ AU test to statistically test for the significance of the discrepancies. Assuming the BI phylogeny always yields a philosophically defensible, character-state based topology, we evaluate the performance of MJN analyses by scoring the percentages where two topologies differ significantly.

## 3 Methods

### 3.1 Data collection

We collected a set of published studies that computed MJNs in their analyses. We particularly looked for MJNs with no or a few cycles. While star-like MJNs, in which all nodes are individually connected to a central connection node, were most frequent in the literature, we avoided them as they are likely to yield a completely unresolved BI tree (pers. obs.). Starlike MJNs have no phylogenetic information. However, Bandelt et al. (1995) argued that such topologies characterize demographic expansions around a founder population, which is represented by the central node, in the past. The selected studies were further filtered by the availability of GenBank accession numbers for the DNA sequences used to compute the MJNs.

The outgroup sequences were also retrieved when available in the original article, although they were not available in most cases because the MJN method does not conduct outgroup rooting for its calculation. When the outgroup was absent, we selected outgroup sequences using two strategies: (1) we identified outgroup sequences from other studies on the same organisms in interest or (2) we manually selected them via Basic Local Alignment Search Tool (BLAST) implemented in GENEIOUS R6 (version 6.1.8; Kearse et al., 2012). Because the ingroup sequences were often from a single (or closely related) species that are expected to be very similar with each other, outgroup taxa that were excessively closely or distantly related were not expected to be very useful in reconstructing relationships. We selected multiple taxa within the sister group when possible, which produced more consistent results than any other outgroup selection strategies (Luo et al., 2010), preferably those with low pairwise identical sites (i.e., *<* 80%) and/or with high identical site (i.e., *>* 95%) compared to the ingroup sequences.

Multiple sequence alignment was performed with MUSCLE (Edgar, 2004) implemented in GENEIOUS R6. All alignments were performed using default parameters, and they were visually inspected and manually corrected whenever appropriate. Two sets of alignments, one without outgroup (denoted by Alignment 1 hereafter) and another with outgroup (Alignment 2), were saved.

### 3.2 Topology inference

For each Alignment 1, we computed a MJN using NETWORK (ver. 4.6.1.2; Bandelt et al., 1999) with the default setting, where a weight applied to each character state (i.e., nucleotide) and an explicit parameter *ε* were set to 10 and 0, respectively. An equal weight was applied to all characters differences in the alignment to the construction of MJNs, although Bandelt et al. (1999) suggest to increase the value in case some character states were more informative in identifying the clusterings than the others, assuming mutations in conservative sites were more valuable than the mutations in hypervariable sites. The explicit parameter *ε* specified a weighted genetic distance to the sampled sequences in the dataset, within which potential median vectors (unsampled hypothetical vectors that linked either sampled vectors, median vectors, or both) may have been constructed. For example, a greater value of *ε* would have resulted in the construction of more median vectors between haplotypes, thus yielding very complex networks (i.e., full median networks) that were often impossible to interpret. Because we aimed to remove cycles in the network, we kept *ε* at its lowest setting. The Optional Post-processing/Maximum Parsimony (MP) calculation implemented in NETWORK (Polzin and Daneshmand, 2003), which would have taken all produced median vectors and linkages into account, was not conducted (see Kong et al. (2016) for arguments against the so-called MP in MJN analysis).

After constructing the network, we manually eliminated any cycle constructed by the MJ method to allow tree topologies to be compared. When one or more haplotypes created a cycle, which we referred to as ‘causative haplotype(s)’, these were arbitrarily chosen and deleted from the alignment. For example, if the MJN analysis yielded a network with a cycle shown in Figure 1, the elimination of any one of the haplotypes A, B, or C resulted in a network free of cycles; haplotypes D and E were not causative haplotypes because the cycle would persist even when these were omitted. The deletion of causative haplotypes and the construction of MJNs were repeated until the final network was free of cycles. Identical deletion of the causative haplotypes was also done for Alignment 2, and this was termed Alignment 2a.

**Figure 1:**
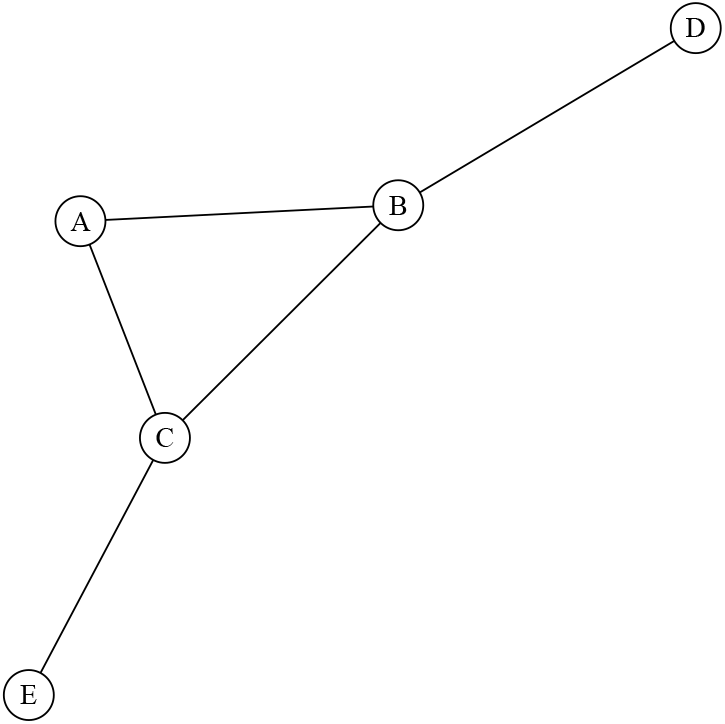
A Median Joining Network (MJN) with five haplotypes (labeled A–E) and a cycle. Eliminating any of the causative haplotypes A, B, or C will result in a network free of cycles. Haplotypes D and E are not causative haplotypes as their deletion will not remove the cycle.

Bayesian inference phylogenies were reconstructed for Alignment 2a and rooted with an outgroup in the alignment. First, we employed MrModelTest (version 2.3; Posada and Crandall, 1998) to select the best DNA substitution mode by the Akaike information criterion. Next, we conducted a single MCMC analysis of 10^7^ iterations in MrBayes (version 3.2.2; Huelsenbeck and Ronquist, 2001), each of which had four chains (three hot and one cold) with the default priors. One tree was saved every 10^3^ generations. The analysis was stopped if the standard deviation of split frequencies fell below 0.01 after the 10^7^ generations. If not, 10^4^ generations were added until the value fell below 0.01. The final 50% majority rule consensus tree was saved.

Because the BI phylogenies incorporated information about the direction of evolution among the ingroup haplotypes, we used this directionality to manually convert each cycle-free unrooted MJN into a rooted branching topology following Salzburger et al. (2011) with minor modifications. We saved the BI tree and the transformed and rooted MJN topology in Newick format. The largest challenge in transforming the cycle-free MJNs into trees was locating a root that would provide the same directionality as in BI phylogenies. When such root position was not present in MJNs, we selected the largest haplotype cluster or the longest edge in MJN as a starting point of the evolutionary direction. When all haplotype clusters were in equal size, we arbitrarily selected a place that produced the most similar topology as the BI tree. In rare cases, where multiple root positions were equally plausible, we produced multiple MJN topologies using different root positions. Finally, the outgroup taxa in the BI phylogenies were removed before statistical comparison to ensure two (or more in case of multiple plausible roots for the MJN) topologies contained identical sets of tips.

### 3.3 Analysis

The topologies obtained from MJN and the BI analyses were statistically compared to evaluate their topological incongruence using an AU test (Shimodaira, 2002) implemented in the software CONSEL (Shimodaira and Hasegawa, 2001) as well as SH test implemented in PAUP* 4.0b10 (Swofford, 2003). Site-wise log-likelihood matrix for each tree using Alignment 1 was obtained using PAUP* followed by execution of SH test with 10^3^ bootstrap replicates using the Resampling of Estimated Log Likelihoods (RELL) method (Kishino et al., 1990). Because CONSEL did not read the matrix from PAUP*, the output matrix was transformed into CONSEL-readable format using a python script written by Mark Holder (available on https://phylo.bio.ku.edu/slides/lab7-Testing/paup-site-like-forconsel.py). The transformed matrix file was treated using `makermt` function in CONSEL to read the matrix and generate 10^5^ bootstrap replicates using the RELL method. The resulting file with extension .`rmt` was processed with CONSEL, which read the bootstrap-replicates of log-likelihood from `makermt`, and calculated *p*-values for the AU test, followed by `catpv` function to visualize the final *p*-values with the significance level *α* set as 0.05.

The AU test, which was originally intended to assess the confidence for a tree among a set of optimal trees, computes the *p*-value to the difference in likelihood between the input topologies based on the same dataset that represents the possibility that the tree is the true tree. This property of *p*-value may be informative if the tree was inferred under a single methodology or in different methods that are equally reliable. We have prior knowledge that the MJN method is a philosophically unsuitable for phylogenetic inference (see Kong et al., 2016) due to its phenetic-based algorithm. Therefore, we assumed that the *p*-value only reflected significant differences in topologies, and not the probability of being correct.

## 4 Results

Figure 2 summarizes the number of sequences, the scale (i.e., intra- or interspecific) and type (i.e., mitochondrial or nuclear DNA) of data of the 85 empirical datasets extracted from 55 studies analyzed in this study. Additional information on the sequence length, nucleotide model selected, *p*-values from AU and SH tests for both MJN and BI topologies, and the source where the sequences were extracted from for each case is summarized in the Supplementary Material 1. Focusing on a diversity of organisms, a total of 4149 ingroup DNA sequences were analyzed, and each dataset contained from seven to 212 sequences. A total of 265 outgroup sequences were included, ranging from one to six per dataset. Seventyfive out of 85 datasets contained less than 100 sequences and 57 datasets contained less than 50 sequences. The shortest length of sequence alignment was 235 base pairs (bp) and the longest was 6490 bp. Further, 68 out of 85 datasets had the sequence length of less than 1000 bp after alignment and only one dataset had sequence length of more than 2000 bp. Fifty out of 85 datasets (58.8%) were mitochondrial loci and 35 (41.2%) involved nuclear genes. Forty-nine (57.6%) out of 85 datasets centered on one species and 36 (42.4%) contained two or more species. The computed MJNs for each dataset are available in the Supplementary Material 2. Mesquite executable NEXUS files, each of which contains sequence alignment, 50% majority rule consensus BI phylogeny, and the transformed MJN topology for each case, are provided in the Supplementary Material 3.

**Figure 2:**
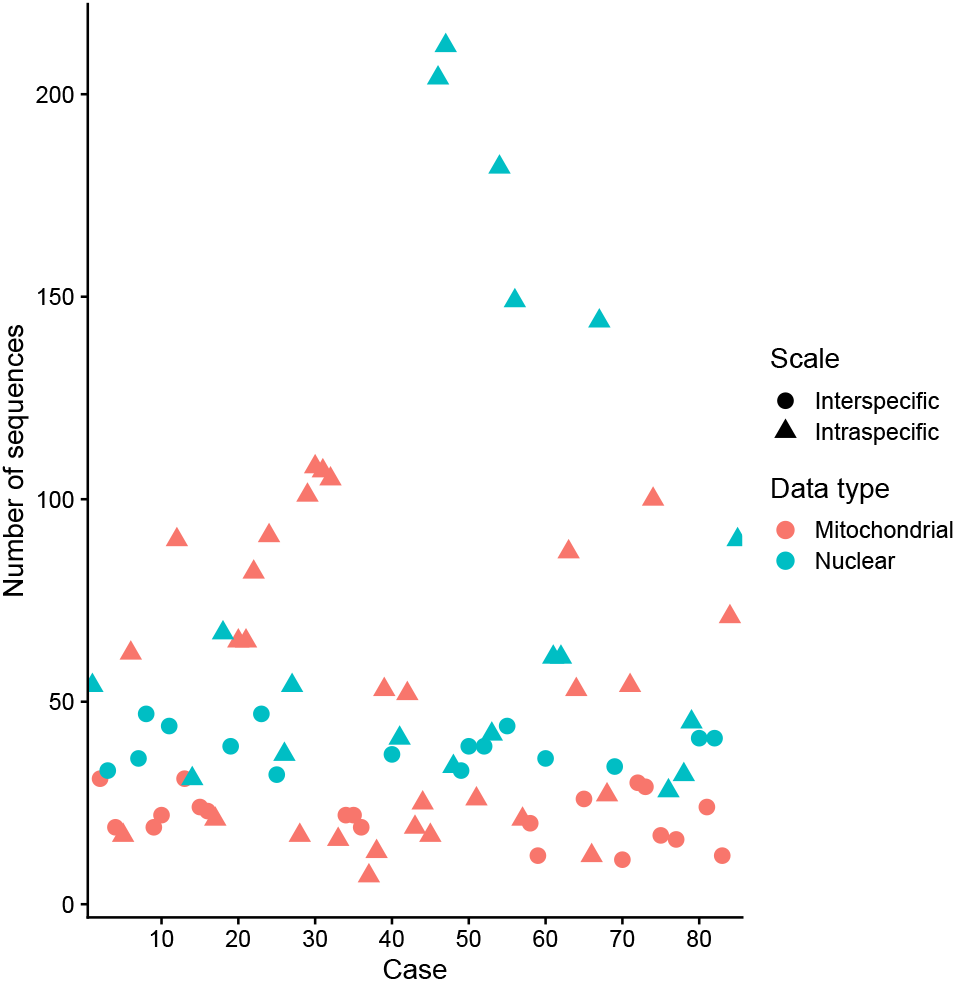
Summary of the 85 analyses conducted in this study. Y-axis represents the number of sequences. Each point represents a dataset, where circles and triangles represent whether the dataset contains inter- or intraspecific terminals, respectively. Red and green color represents whether the loci in the datasets are mitochondrial or nuclear DNA.

Topologies differed significantly (AU *p ≤* 0.05) in 32 out of 85 cases (37.6%). Among the 53 cases where the significance was not observed, 51 MJN and BI topologies (96.3%) showed some discrepancies in branching patterns, and the topologies were identical in two cases only. The discrepancies included positions of clades (or clusters in MJN), constituents of a clade (or cluster), or the branching pattern of the topology (see Case Studies below).

Significant topological discrepancies were not dependent on whether the dataset was intra- or interspecific, mitochondrial or nuclear DNA, or sequence length of the dataset. While the difference of likelihood between the two topologies was generally small (i.e., *<* 100) in most of the cases, four cases showed relatively large difference (191.7–802.2), and significance was found for all four.

While SH and AU tests generally exhibited similar results, an overly conservative pattern in the SH test was observed as stressed by Strimmer and Rambaut (2002) and Shi et al. (2005). When two competing topologies were not significantly different, both tests agreed in all cases. The two tests agreed in 16 out of 32 cases when the significance was observed between the topologies. However, the two tests did not agree in 16 out of 32 cases when discrepancies between the two topologies were observed, and in all of these cases, significance found using the AU test was veiled in the SH test.

### 4.1 Case study 1: Significantly different topologies

Yu et al. (2012) analyzed mitochondrial DNA sequences from *Ochotona curzoniae* (the plateau pika) of the Qinghai-Tibetan Plateau, China. We selected 27 D-Loop sequences (GenBank accession numbers JN165313–17; JN165320; JN165322; JN165324–29; JN165332, JN1653323; JN165335–45; JN165350) that were included in the MJN presented in Figure 3a of Yu et al. (2012). Note that some haplotypes used to compute the original MJN were excluded in our analysis to eliminate unwanted cycles. We selected three outgroup sequences of *O. collaris* (AF348080), *O. princeps* (AJ537415) and *Lepus sinensis* (KM362831) where the former two were suggested in the original study and the latter was selected using BLAST.

**Figure 3:**
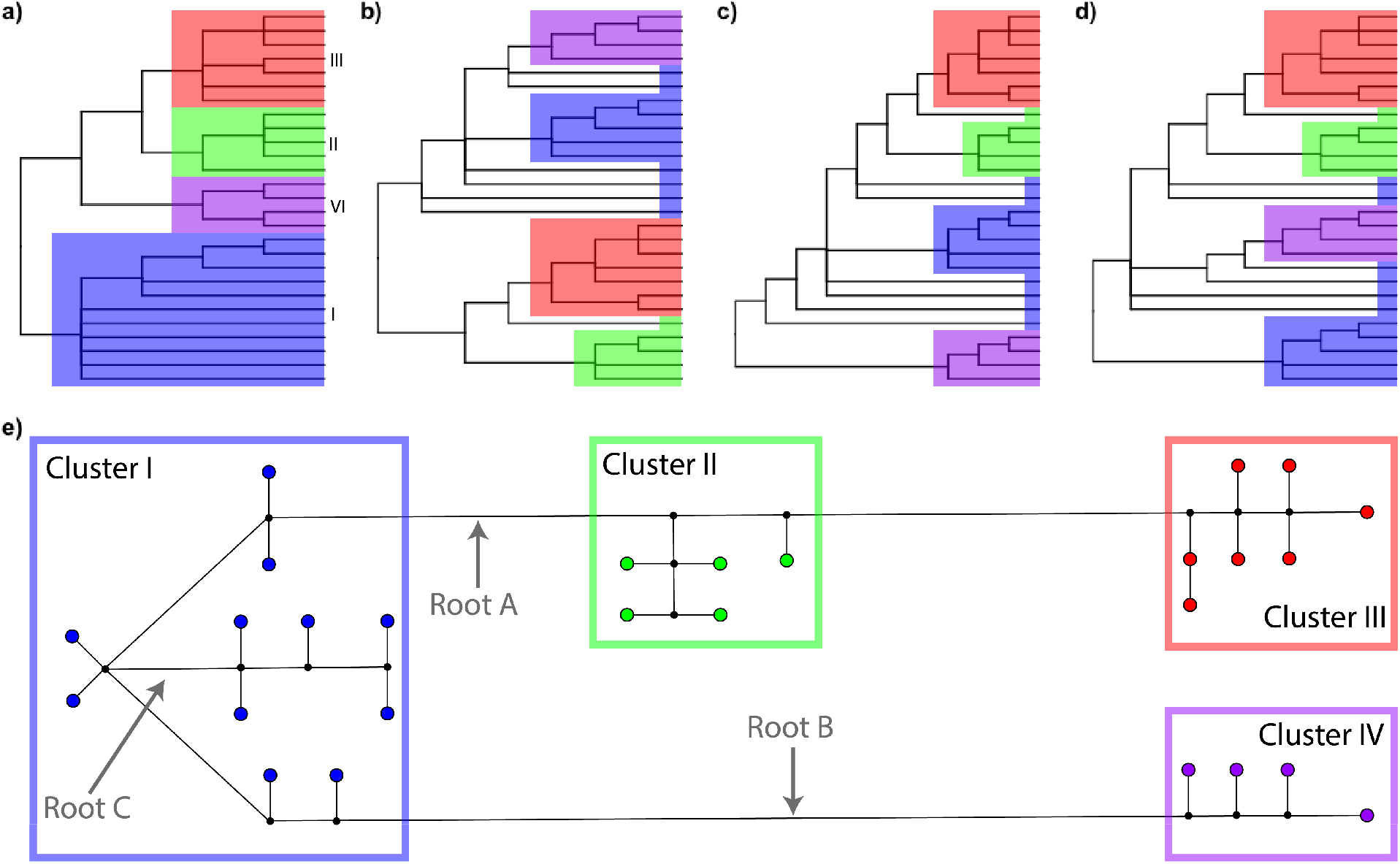
(a) Reconstructed 50% majority-rule consensus Bayesian inference (BI) phylogeny using 27 *Ochotona curzoniae* D-Loop sequences from Yu et al. (2012), rooted with outgroup sequences (not shown here). (b) MJN topology using the Root A, (c) Root B, and (d) Root C, that are selected in the computed MJN in (e). Four clades were identified and labeled in the BI phylogeny, and we color coded the members of each clade that is applied for all topologies and MJN. Small black dots in the MJN represent computed median vectors.

The final alignment was 672 bp long. We set *ε*=0 and weight=10 for computing the MJN. No preor post-processing option was taken. Each sequence represented an active, unique haplotype (i.e., 27 terminal vertices exist in the MJN) and a total of 142 mutations were counted for the shortest network. For BI analysis, the GTR+I+Γ substitution model was selected for the sequence evolution and a single MCMC analysis of 10^7^ iterations was performed.

Figure 3 shows the BI 50% majority-rule consensus phylogeny (Fig. 3a), MJN (Fig. 3e), and three transformed MJNs using the root positions A, B, and C (Fig. 3b–d, respectively) that are indicated in the MJN. The BI phylogeny had four primary clades (Clades I, II, III, and IV are highlighted in blue, green, red, and purple, respectively, in Figure 3a), and their nodes, which represented the most recent common ancestor (MRCA), had posterior probabilities ranging from 0.86 to 0.99, except for Clade I with 0.62. Four clusters in the MJN corresponded to the BI phylogeny (the sequences that belong to the Clades I–IV in the BI phylogeny are colored using the same scheme described above in Figure 3e), but no root position that would provide the same evolutionary direction as in the BI phylogeny was available. The three most probable root positions were A, B, and C in Figure 3e. Roots A and B were placed on the edges with the longest lengths. Although the edge between Clusters II and III was long, it was not selected as a possible root placement because the two clusters were sister taxa in the BI tree. Because a root may not necessarily position on long edges, we also identified root C as it retained Clades II and III despite the short length that separated some members of Cluster I. None of the topologies resolved monophyly of Clade I (Figs. 3b–d). PAUP* was used to calculate site-wise log-likelihood scores for all four rooted topologies using the GTR+I+Γ nucleotide substitution model.

The AU test (Table 1) showed that the BI topology differed significantly in branching pattern from all three MJN topologies (*p ≤* 0.05). In comparison, the SH test showed that the BI topology differed significantly from the MJN using root C only, although root positions A and B had small *p*-values.

**Table 1:** The Approximately Unbiansed (AU) and Shimodaira-Hasegawa (SH) test results for four topologies from BI and MJNs using three different roots of *Ochotona curzoniae* D-Loop sequences from Yu et al. (2012). For each tests for the topology listed under the column labeled as Item, Obs. represents the observed log-likelihood differences, AU and SH represent their computed *p*-values.

| Topology | Item | Obs. | AU | SH |
| --- | --- | --- | --- | --- |
| 1 | BI | -13.2 | 0.960 | 0.954 |
| 2 | MJN, root A | 13.2 | 0.046 | 0.055 |
| 3 | MJN, root B | 13.2 | 0.045 | 0.055 |
| 4 | MJN, root C | 13.2 | 0.031 | 0.042 |

**Table 2:** AU and SH test results for topologies from BI and MJN of *Coreoperca whiteheadi* cytochrome b sequences from Cao et al. (2013). Obs. represents the observed log-likelihood differences, AU and SH represent p-values for each tests for the topology listed under the column labeled as Item.

| Topology | Item | Obs. | AU | SH |
| --- | --- | --- | --- | --- |
| 1 | BI | -4.3 | 0.857 | 0.825 |
| 2 | MJN | 4.3 | 0.143 | 0.175 |

### 4.2 Case study 2: Insignificantly different topologies

We used the mitochondrial cytochrome b sequence dataset of *Coreoperca whiteheadi* (East Asian perch) from Cao et al. (2013). We selected 53 sequences of *C. whiteheadi* (JN315557– JN315609) along with outgroup sequences from *Lateolabrax japonicus* (DQ351531) and *Siniperca chuatsi* (KC888071), as suggested by the authors, plus *Trachinotus rhodopus* (AY050739) selected via BLAST. The length of final alignment was 1141 bp.

Analyses replicated the MJN in Figure 3a of Cao et al. (2013). The 11 haplotypes contained from one to 19 sequences, and the final network identified 203 mutations among the haplotypes. We used the GTR +I+Γ nucleotide substitution model for the BI phylogeny. The identified position of the root on the MJN (Fig. 4c) provided the same directionality as in the BI phylogeny (Fig. 4a). The root occurred on the longest edge of the MJN that separated Cluster IV from the rest (Fig. 4c), which represented 106 nucleotide differences. We converted the MJN into a tree using the selected root position (Fig. 4b).

**Figure 4:**
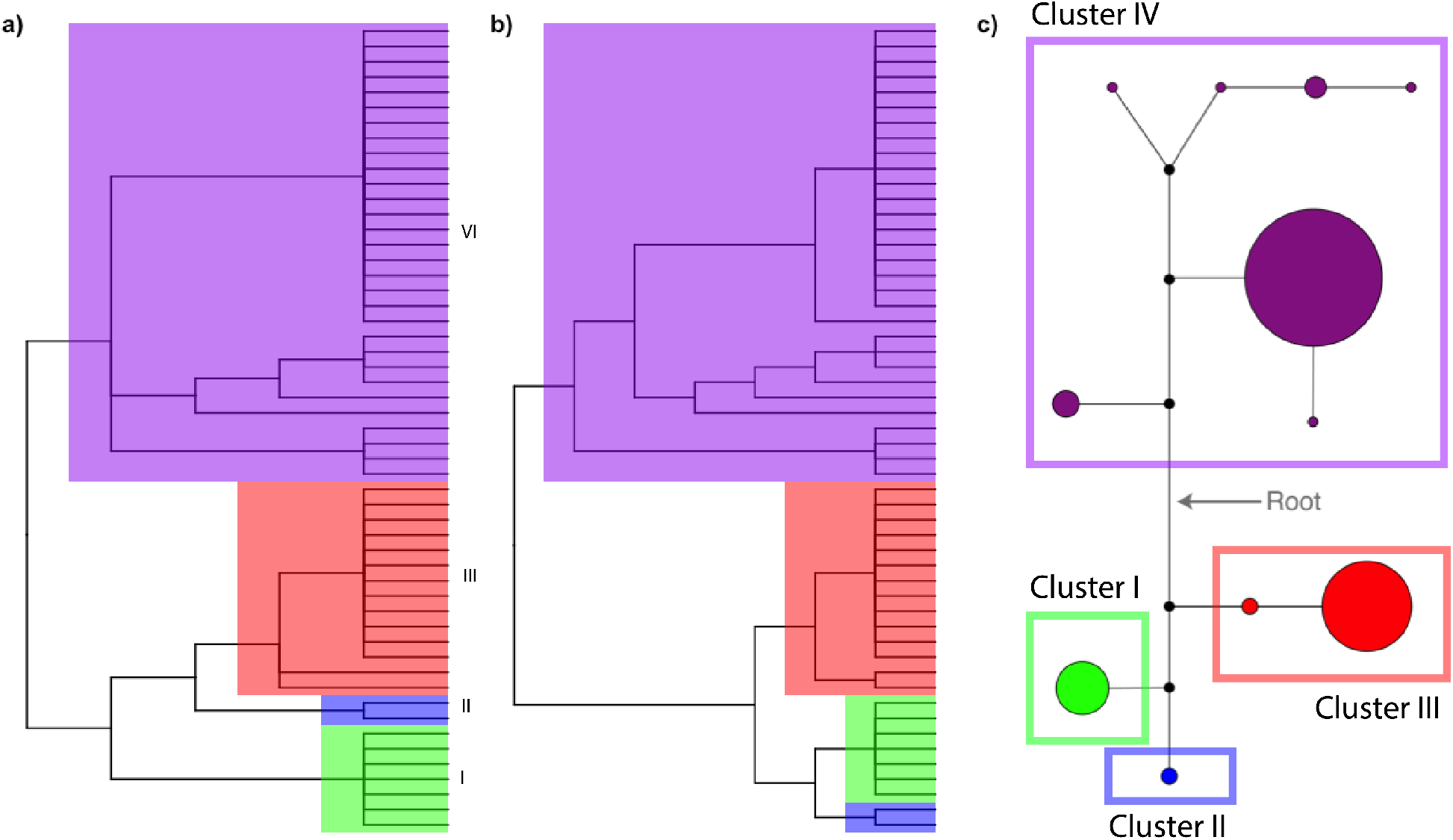
(a) Reconstructed 50% majority-rule consensus BI phylogenetic tree using 53 *Coreoperca whiteheadi* cytochrome b sequences from Cao et al. (2013), rooted with outgroup sequences (not shown here). (b) MJN topology using the identified root position in MJN shown in (c). Four clades were identified in BI phylogeny and we color coded each clade. Same color scheme was applied for all topologies and MJN. Four clades were identified and labeled in BI phylogeny, and we color coded the members of each clade that is applied for all topologies and MJN. Small black dots in the MJN represent computed median vectors.

The BI phylogeny had four primary clades (Clades I, II, III, and IV are highlighted in green, blue, red, and purple, respectively, in Figure 4a), although the relationships among some haplotypes were not clearly resolved in the BI tree, possibly due to the artifact of consensus tree production and the closeness of the sequences. However, the relationships between Clades I, II, and III were inconsistent between the BI phylogeny and MJN topology (Figs. 4a and 4b, respectively). In the BI phylogeny, Clades II and III shared a MRCA with posterior probability of 1.0. However, the MJN grouped Clusters I with II. Despite the differences in branching pattern, the relationships were not significantly different based on both the AU and SH test (Table 3). Although no significant difference occurred in the AU and SH tests, the biological interpretations based on the MJN would be problematic due to differences in the resolved branching order.

## 5 Discussion

### 5.1 Problems of Median-Joining Network analysis

Analyses find statistically significant discrepancies between topologies from BI and MJN in 37.6% of the cases evaluated. If BI produces a more reliable evolutionary history than MJN, then MJ method too often fails to produce a correct suite of relationships for confident evolutionary inference. Even in the 62.4% of cases where statistical significance does not occur, differences in branching patterns often represent different evolutionary histories among the sequences (see case study 2). Therefore, MJ largely produces misleading hypothesis of relationships and character evolution. Phenetic-based MJ has many unrealistic assumptions, such as ignoring homoplastic evolution. This has led many authors to criticize phenetic methods for failing to produce ‘true’ phylogenies (Lawrence et al., 2002, Cheema and Dicks, 2009). While MJ and BI produce marginally different sets of relationships in terms of the AU test output in more than half of the cases, sometimes the differences are critical, for example, when MJ fails to correctly identify monophyletic groups.

One of the biggest problems with MJNs, and probably all un-rooted networks, is the absence of evolutionary direction (Kong et al., 2016). Identification of monophyletic groups is only possible given direction, i.e., ancestor–descendent relationships. An outgroup serves to infer hypothetical ancestral states and in doing so, roots an un-rooted branching structure (Watrous and Wheeler, 1981, Maddison et al., 1984, Smith, 1994, Lyons-Weiler et al., 1998, Sanderson and Shaffer, 2002). Although the software NETWORK allows the user to root the network, this option actually links the outgroup sequence to the most similar haplotype of the already produced ingroup network (Kong et al., 2016, Sánchez-Pacheco et al., 2020). For example, Sakaguchi et al. (2012) employed the rooting option to root the MJN for 26 chloroplast haplotypes, yet the branching pattern differed from the inferred MP phylogeny using the same outgroup. Additionally, the option only allows one sequence to be entered, yet more than one outgroup taxon may be necessary to obtain reproducible results (Maddison et al., 1984, Smith, 1994).

Given its phenetic nature, overall similarity influences the MJN algorithm very strongly, especially in computations involving gaps and ambiguous states. When the amount of gaps is large due to excessive amounts of indels, MJ can create complicated cyclic structures that are almost impossible to interpret. Consequently, it has been recommended to minimize gap sites prior to analysis (Bandelt et al., 1999), but this can be a very subjective process (DeSalle et al., 1994). Moreover, gaps can be phylogenetically informative, improve branch support, and even change a topology (Simmons and Norton, 2014, Nagy et al., 2012). Trimming by the simple removal of the gaps can result in the loss of otherwise informative sites and, thus, it does not necessarily lead to better trees (Löytynoja and Goldman, 2008, Wong et al., 2008, Dessimoz and Gil, 2010, Wu et al., 2012).

The treatment of ambiguous states in the construction of MJNs is also problematic. Bandelt et al. (1999) suggested using ambiguous states in the alignment infrequently. However, MJ tolerates ambiguity codes such as *X* = R (purine) = *{*A, G*}* or *X* = N = {A, T, C, G} and others, where *X* represents the set of nucleotide states that specify the ambiguous state. Here, the MJ algorithm assigns the most common state of the other minimally distant sequences to the ambiguous position before starting. This procedure is arbitrary and can result in the loss of potentially informative variations in the data. To illustrate the problem, we use three hypothetical sequences with five bp each—SQ1=CAACG, SQ2=GCCAC, and SQ3=AGCGA (Fig. 5a)—used in the construction of Figure 3 of Bandelt et al. (1999). The sequences differ at three sites: sites 1 and 2, plus 3 for SQ1, 4 for SQ2, and 5 for SQ3. Figure 5h shows the MJN constructed using the alignment in Fig. 5a.

**Figure 5:**
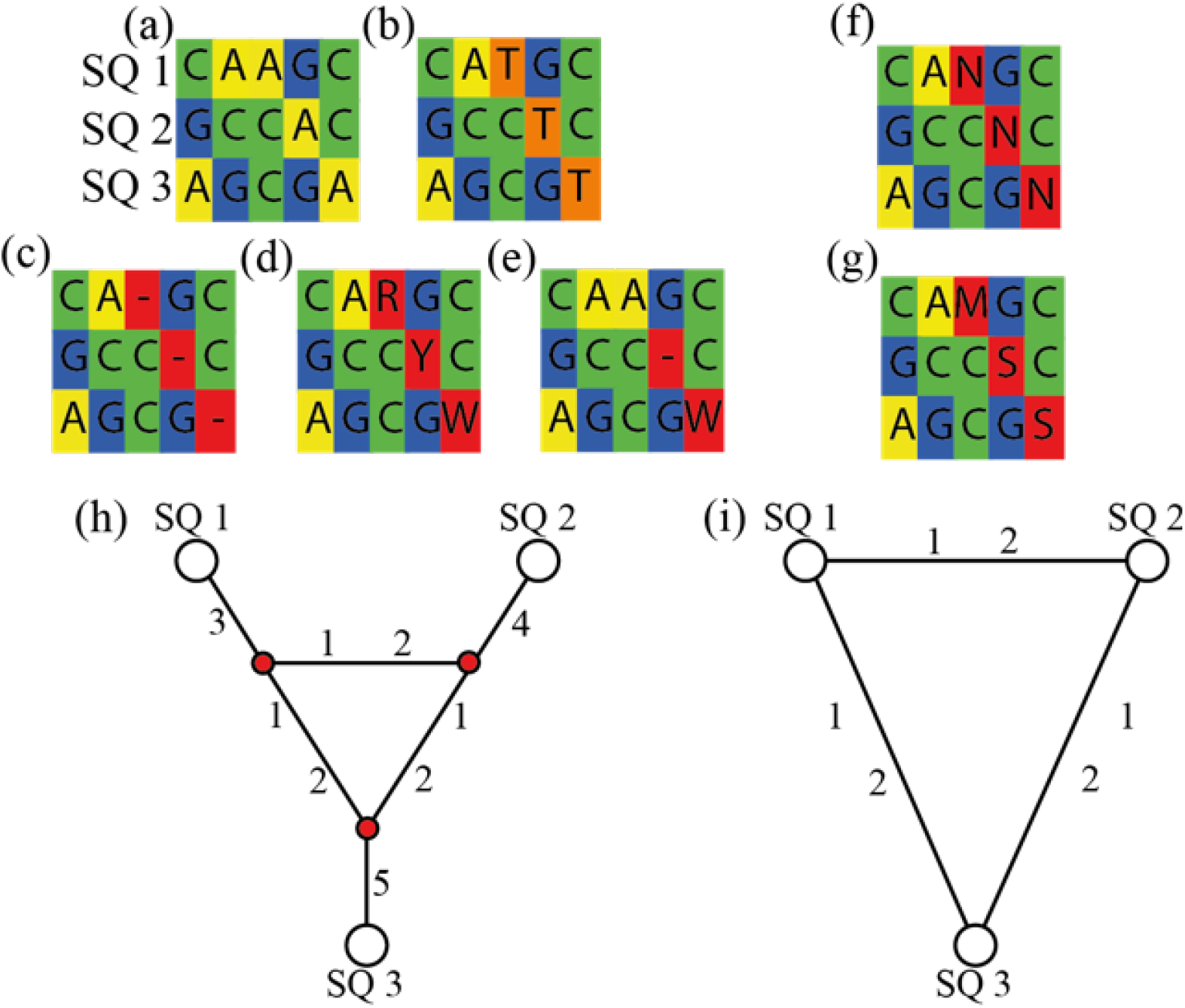
Example of the treatment of ambiguous states in the construction of MJNs. Sequence alignments (a–e) result in the MJN shown in (h) whereas the alignments (f) and (g) result in the MJN (i). Note the two MJNs have different biological interpretations as the algorithm arbitrarily removes variabilities in the presence of the ambiguous sites.

Because MJ simply identifies relationships based on overall similarity, the resulting network is the same when the character states that are unique for each haplotype (i.e., site 3 for SQ1, 4 for SQ2, and 5 for SQ3) differs from another character state, as long as it still distinguishes them. For example, when the sites 3, 4, and 5 in SQ1, SQ2, and SQ3, respectively, have nucleotide state A and then changed to T (Fig. 5b) or a gap (Fig. 5c), the network structure remains as in Figure 5h. Similarly, when the sites 3, 4, and 5 in SQ1, SQ2, and SQ3, respectively, are replaced by ambiguous sites (Fig. 5d) or a mixture of nucleotide, gap, and an ambiguous site (Fig. 5e), the network structure remains unaltered.

Figures 5d and 5e use ambiguous nucleotides that are not the same as in other sequences. For example, ambiguous site R exists in site 3 of SQ1, it can be either A or G, but not C (i.e., the state SQ2 and SQ3 possess at site 3). Similar ambiguous replacements for the other cases illustrate the problem. When we replace SQ1 with N (= A,T,C,G), or M (= A,C), the MJ algorithm arbitrarily switches the site to C by comparing it with the minimally distant sequence; this preference eliminates the median vector on the edges (Fig. 5f, 5g, and 5i) based solely on algorithm-assigned overall similarity. Networks in Figures 5h and 5i do not have identical biological interpretations, and more complex problems can arise when evaluating longer sequences and more taxa.

### 5.2 Excessive conservativeness of the SH test

Researchers often compare the significance of one or a set of optimal trees from different phylogenetic methods (e.g., Kazlauskas et al., 2019, Naser-Khdour et al., 2019, Spence et al., 2021) or how strongly competing topologies can be rejected relative to the preferred one (e.g., Espeland et al., 2018, Miller et al., 2020, Hime et al., 2021, Coleman et al., 2021, Yan et al., 2021) using SH and AU tests. Among many tests that can imply confidence in resolved nodes and trees, the SH and AU tests are most widely used. They employ similar rationale yet selection-bias in the SH test can lead to overconfidence in incorrect trees when more than two trees are compared (Shimodaira and Hasegawa, 1999, Goldman et al., 2000). Moreover, Strimmer and Rambaut (2002) pointed out that the SH test can also be very conservative and the confidence set can increase along with the number of trees being compared increases. Thus, Shimodaira (2002) proposed the AU test, which is less conservative than the SH test.

By comparing the outputs of AU and SH tests, we reconfirm the over-conservativeness of the SH test. For cases in which significant discrepancies exist between the two topologies in at least one of the tests, the two tests arrive at different confidence values in 16 out of 32 cases. Here, the SH test fails to detect significant discrepancies found by the AU test. In other words, the AU test confidently selects one topology out of the two topologies tested, but the SH test selects both trees as being equivalent (or the difference is insignificant) due to its conservative nature. This observation is congruent with the reanalysis of mammalian mitochondrial protein-coding sequences in Shimodaira (2002), in which the SH test resulted in eight confident trees out of 15 competing hypotheses, whereas AU resulted in only six.

## 6 Conclusions

In conclusion, MJN analysis is appealing because it is computationally simple and fast, and yields attractive figures. Nevertheless, MJ should not be used in evolutionary studies given its distance-based phenetics, the absence of direction (Kong et al., 2016, Sánchez-Pacheco et al., 2020), the inherent assumption that all lineages evolve at equal rates (i.e., overall similarity reflects phylogeny), and the extent of statistically significant mismatches between the MJN and BI topologies. The frequency that MJN shows problematic branching patterns, and thus incorrect inferences of evolutionary relationships among taxa, can be higher than 95% (as only two out of 85 cases show identical relationships between MJN and BI topologies). Indeed, the branching patterns produced by MJN analyses differ statistically significantly in major ways from BI trees in 37.6% of our case studies. This is not a trivial problem in an era when researchers are using phylogenetic trees to help solve a wide range of problems from identifying the source of human pathogens to making conservation decisions. Such enterprises require the use of the most robust trees possible, which, in turn, requires the use of defensible and rigorous methods for reconstructing those trees. Our study reinforces the decades-old recognition that phenetic-based algorithms are not a defensible way to hypothesize evolutionary relationships.

## Acknowledgements

We thank M. Holder for advice conducting the AU test.

## Supporting Information

Data available from the Dryad Digital Repository after the peer review.

