## Supplementary_Materials for "Topological incongruence between Median-Joining Networks and Bayesian inference phylogenies": README.txt

Each Mesquite executable (partially executable by PAUP) Nexus file stores sequence alignment, 50% majority rule consensus phylogeny constructed from MRBAYES, transformed MJN topology using the root identified from the BI topology, for the 85 cases evaluated. Note cases 28,33,34,70 are missing topologies and cases 22,25 are missing sequence alignment.
