## Supplementary figures and images for "Topological incongruence between Median-Joining Networks and Bayesian inference phylogenies"

### MJN_case1.pdf

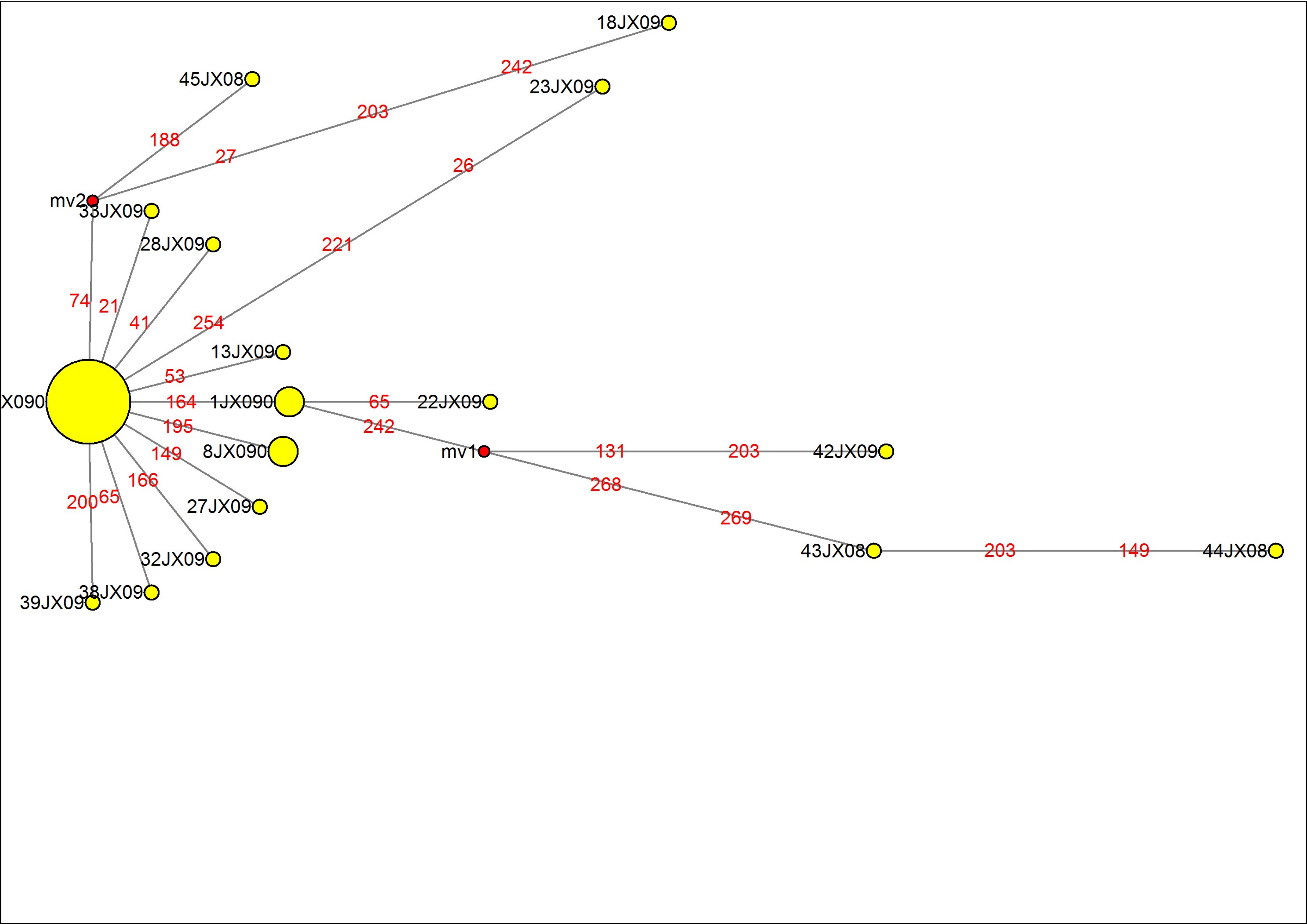

### MJN_case2.pdf

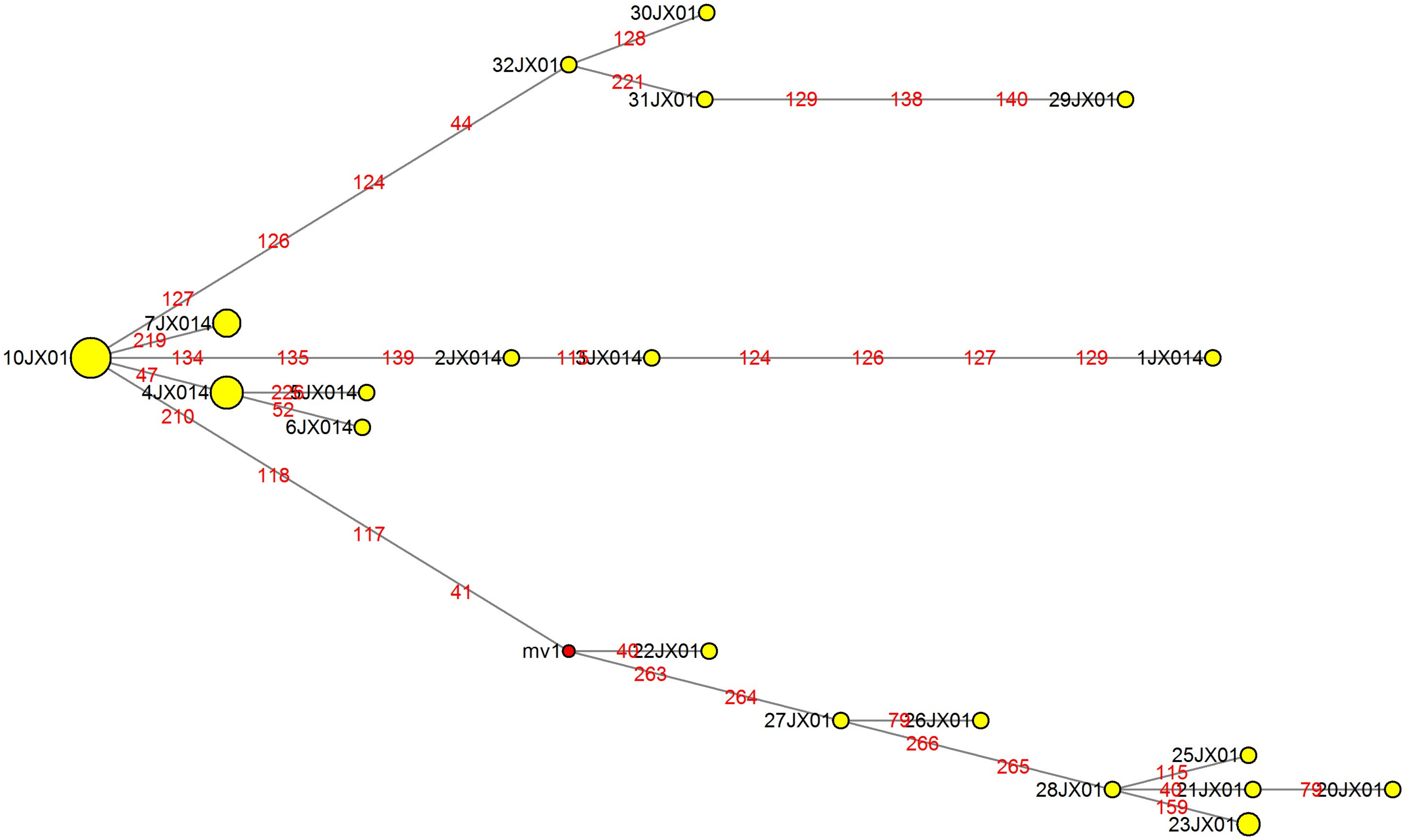

### MJN_case3.pdf

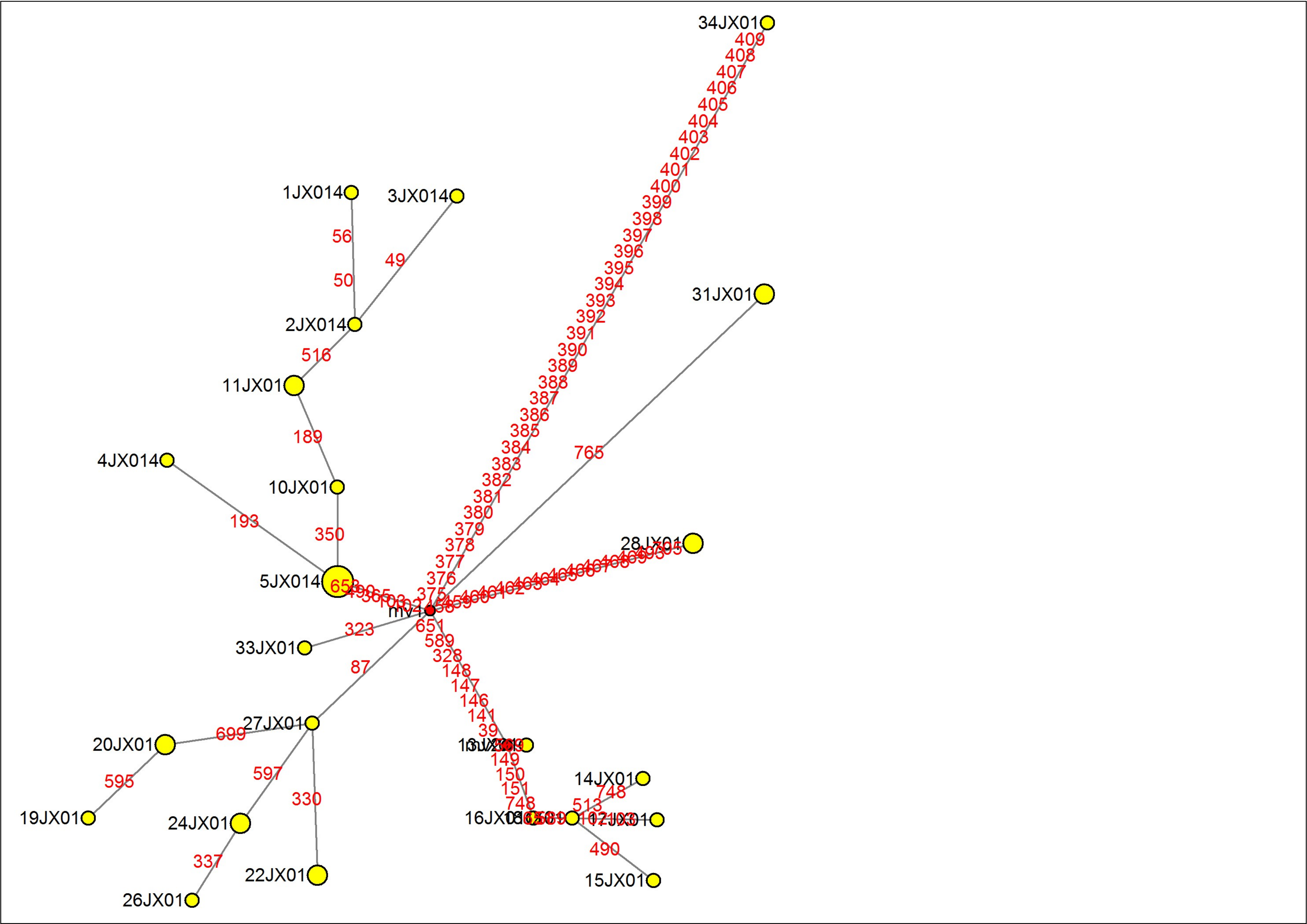

### MJN_case4.pdf

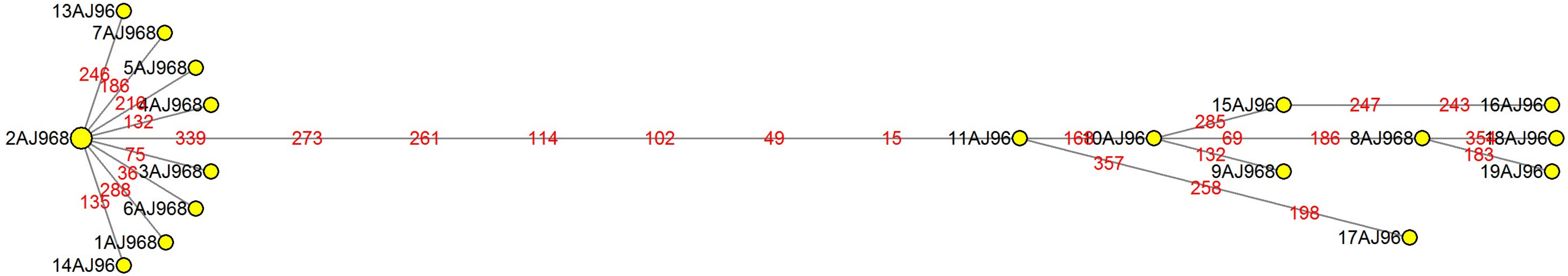

### MJN_case5.pdf

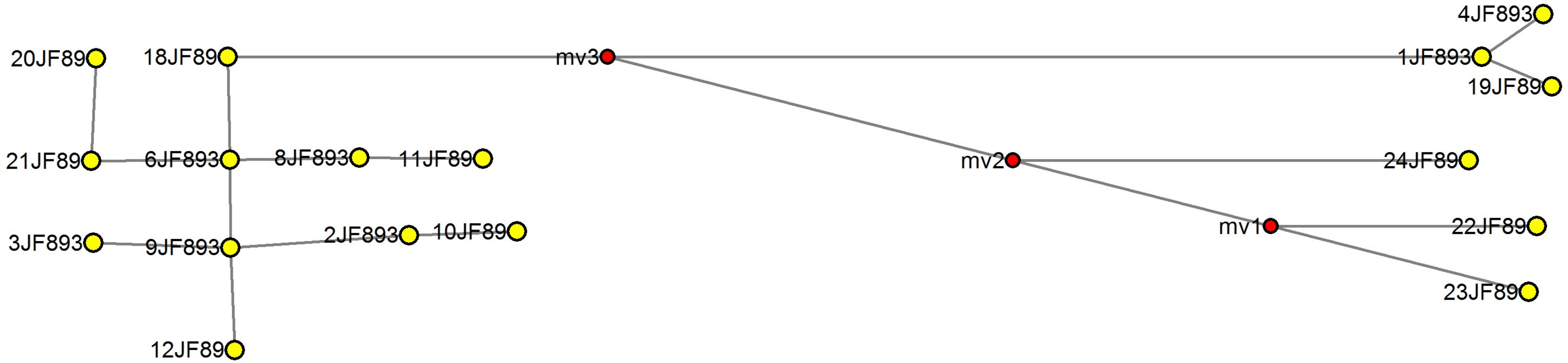

### MJN_case6.pdf

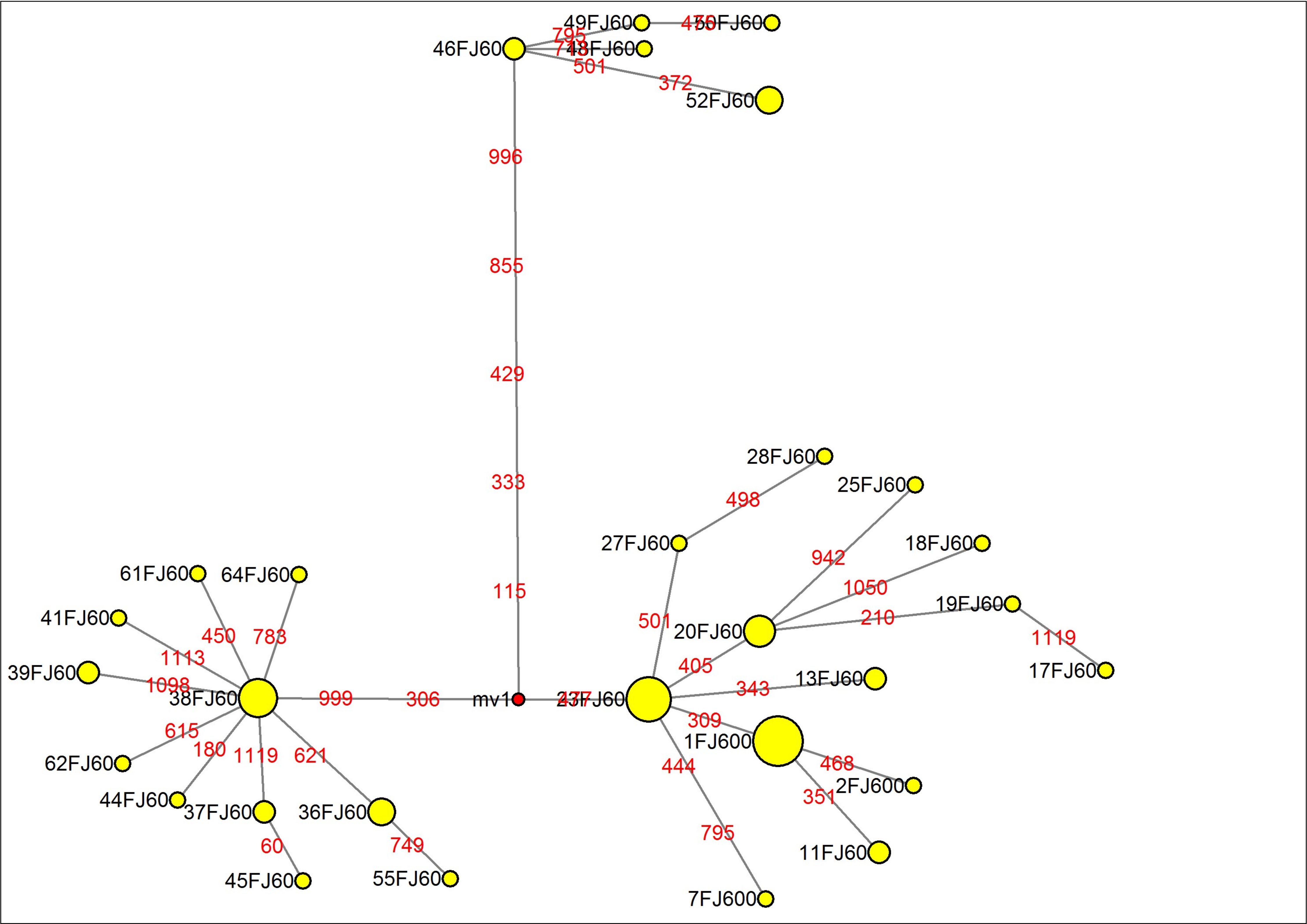

### MJN_case7.pdf

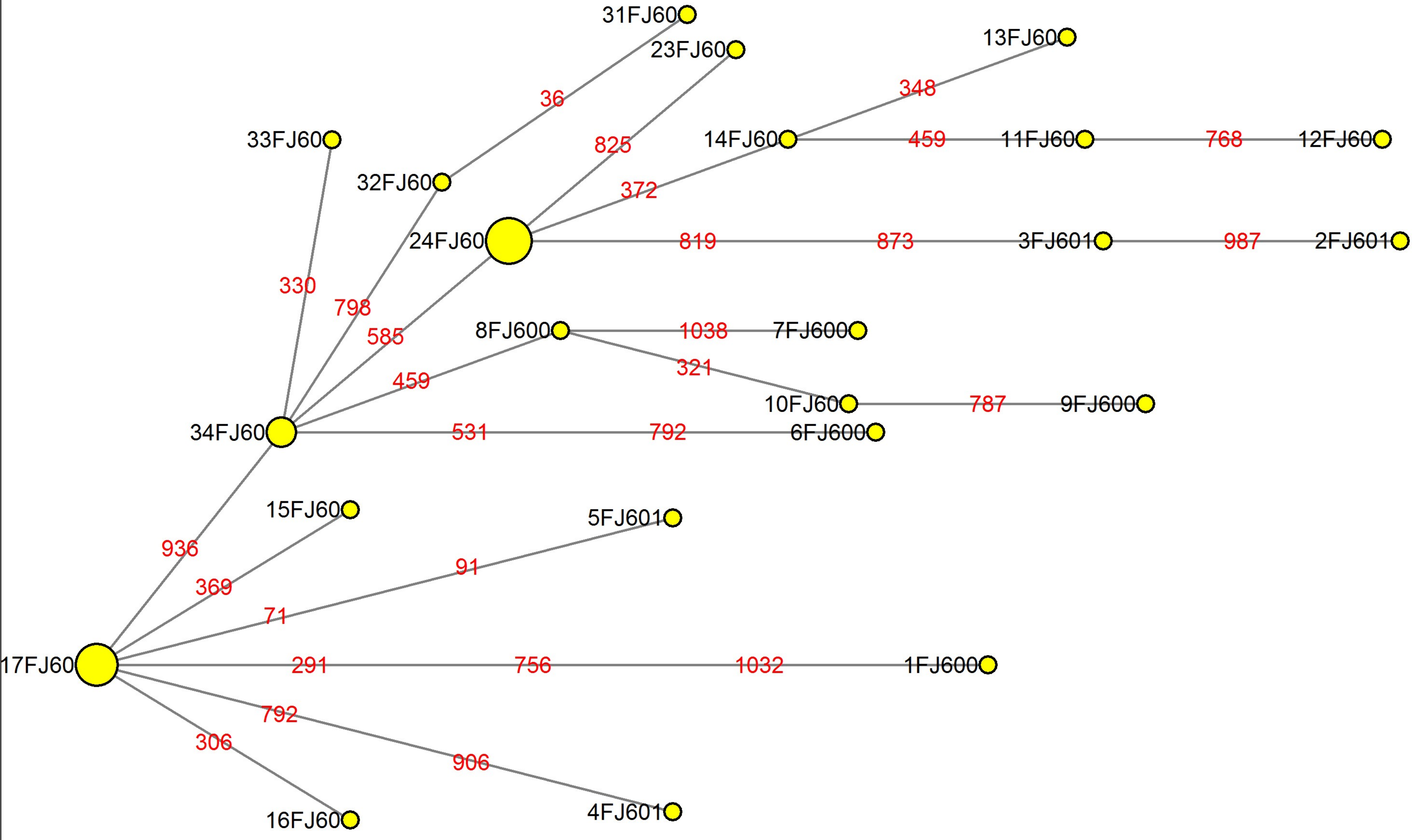

### MJN_case8.pdf

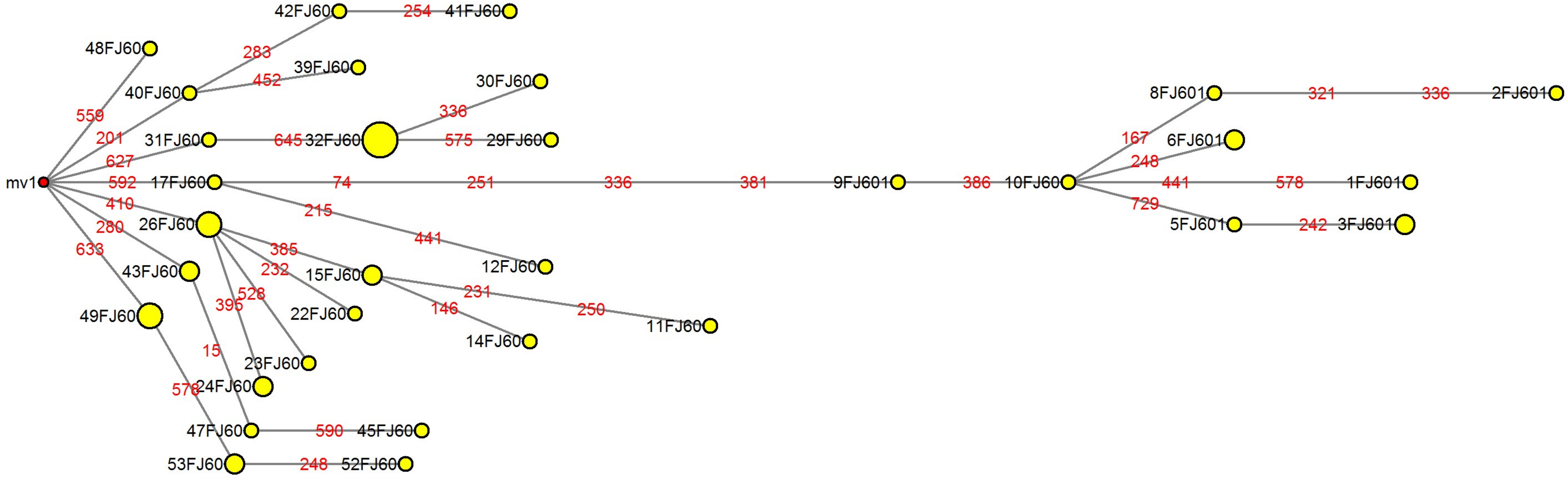

### MJN_case9.pdf

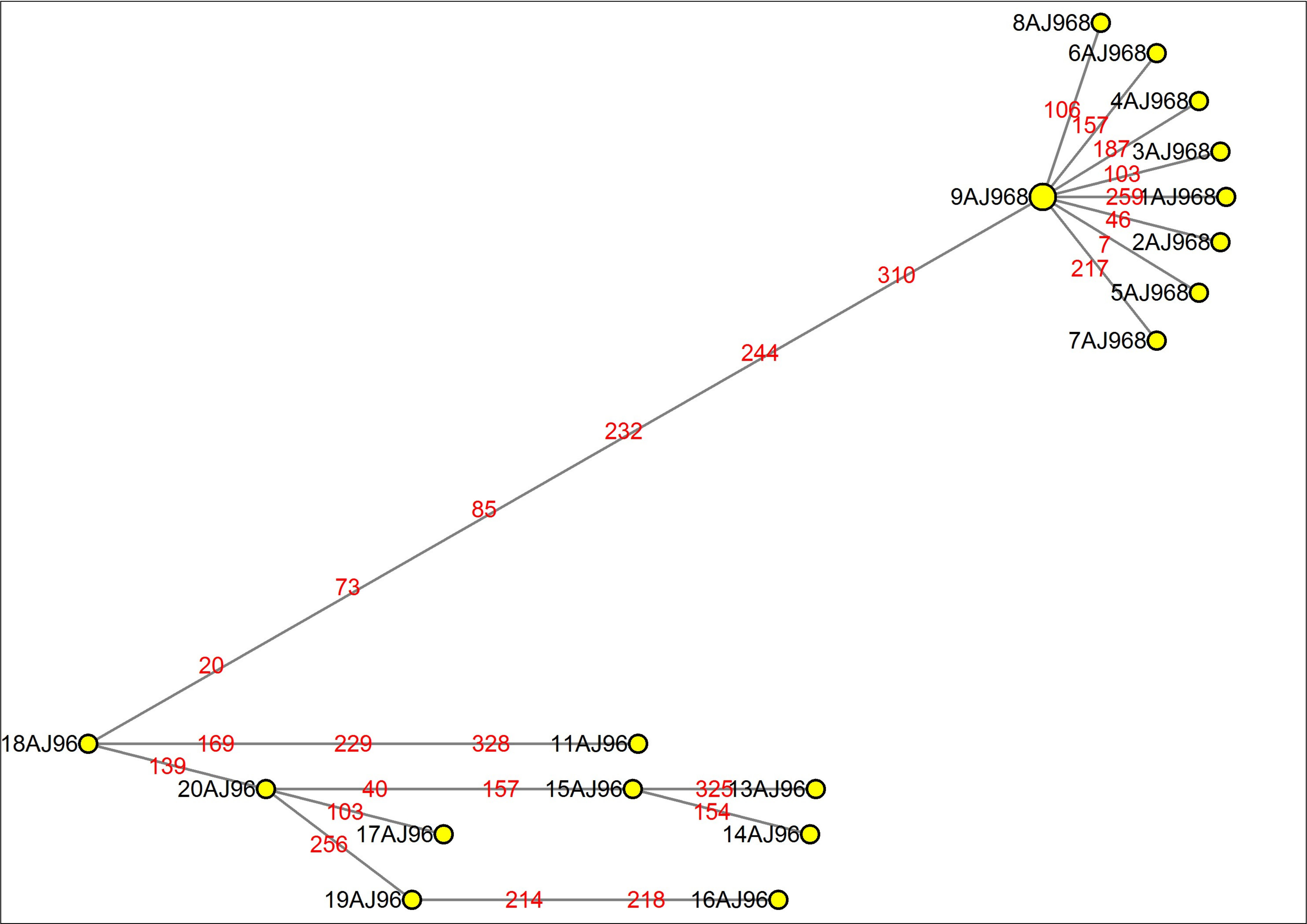

### MJN_case10.pdf

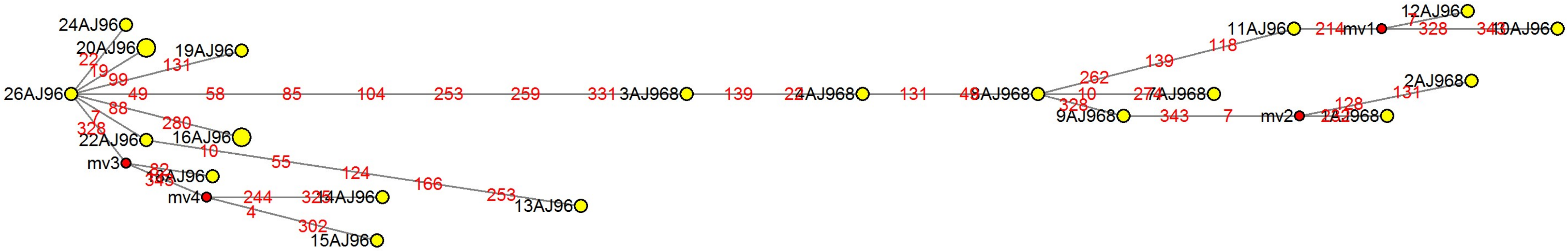

### MJN_case11.pdf

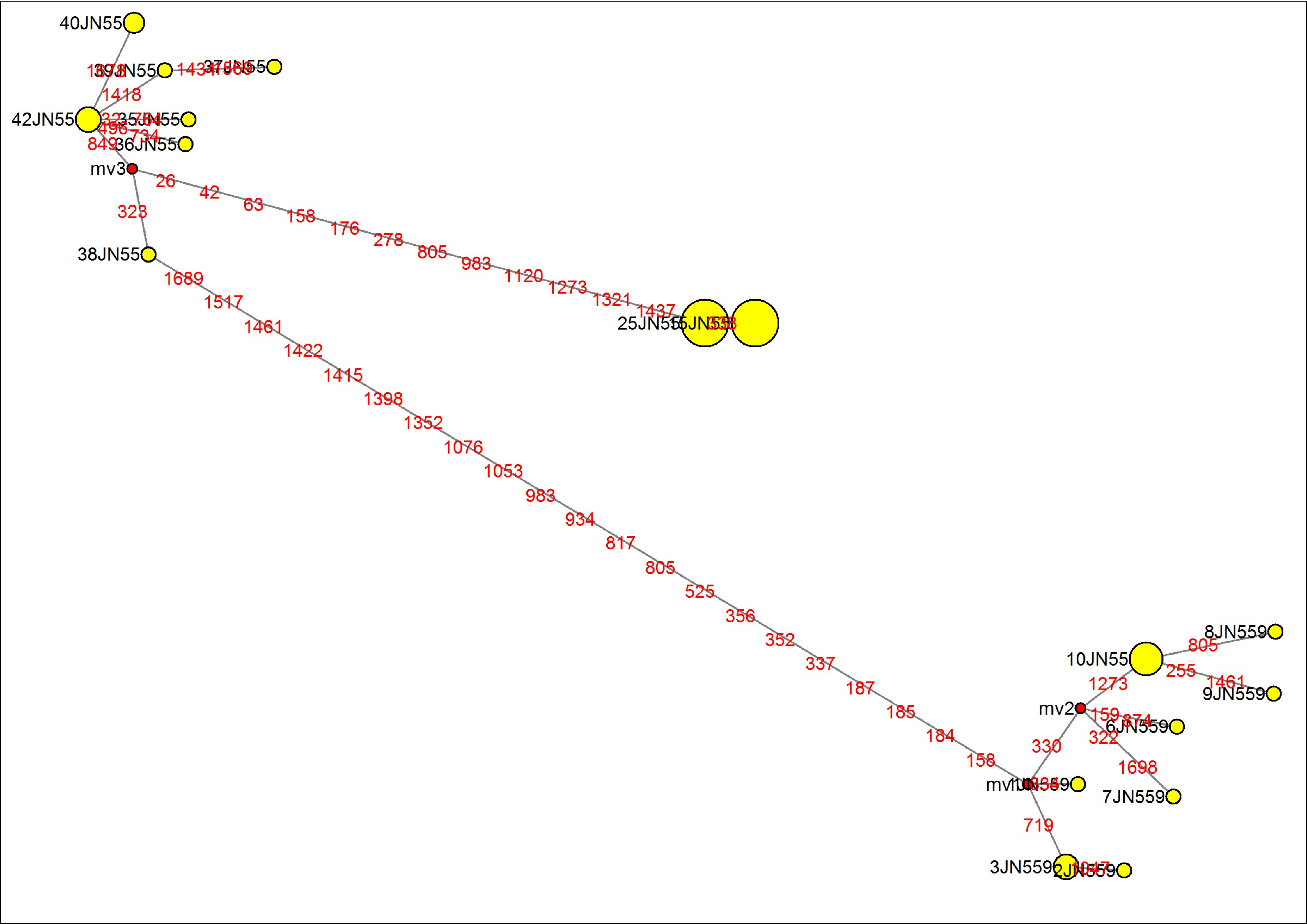

### MJN_case12.pdf

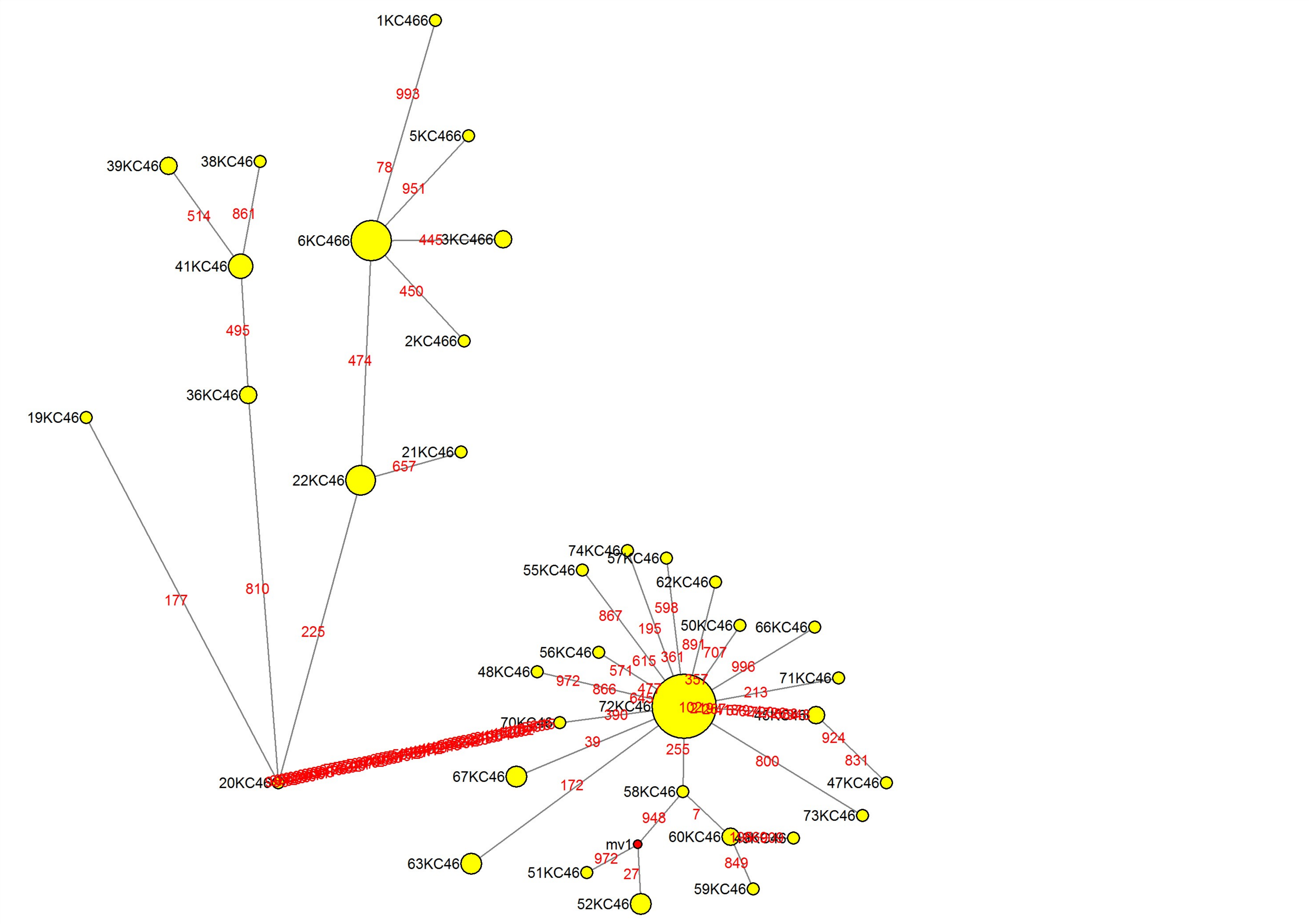

### MJN_case13.pdf

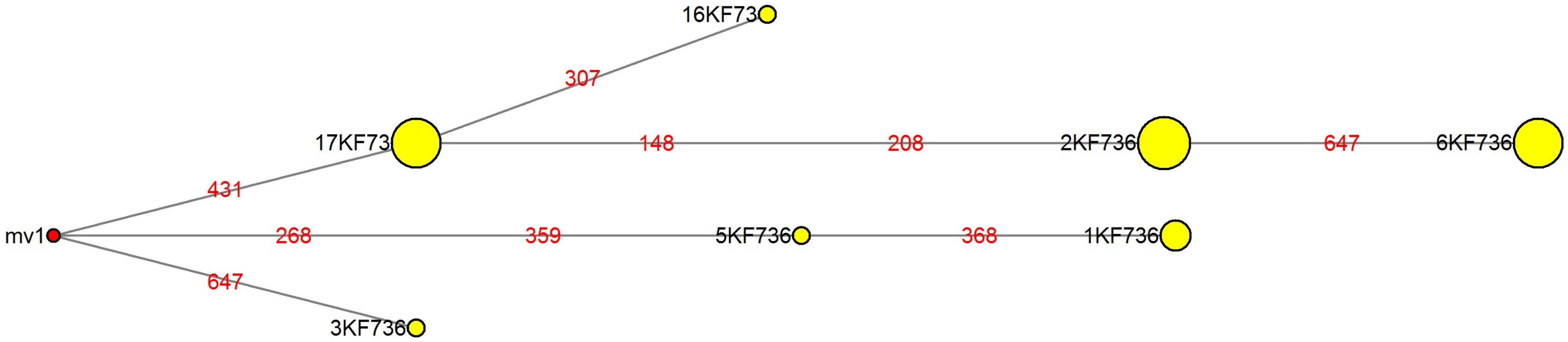

### MJN_case14.pdf

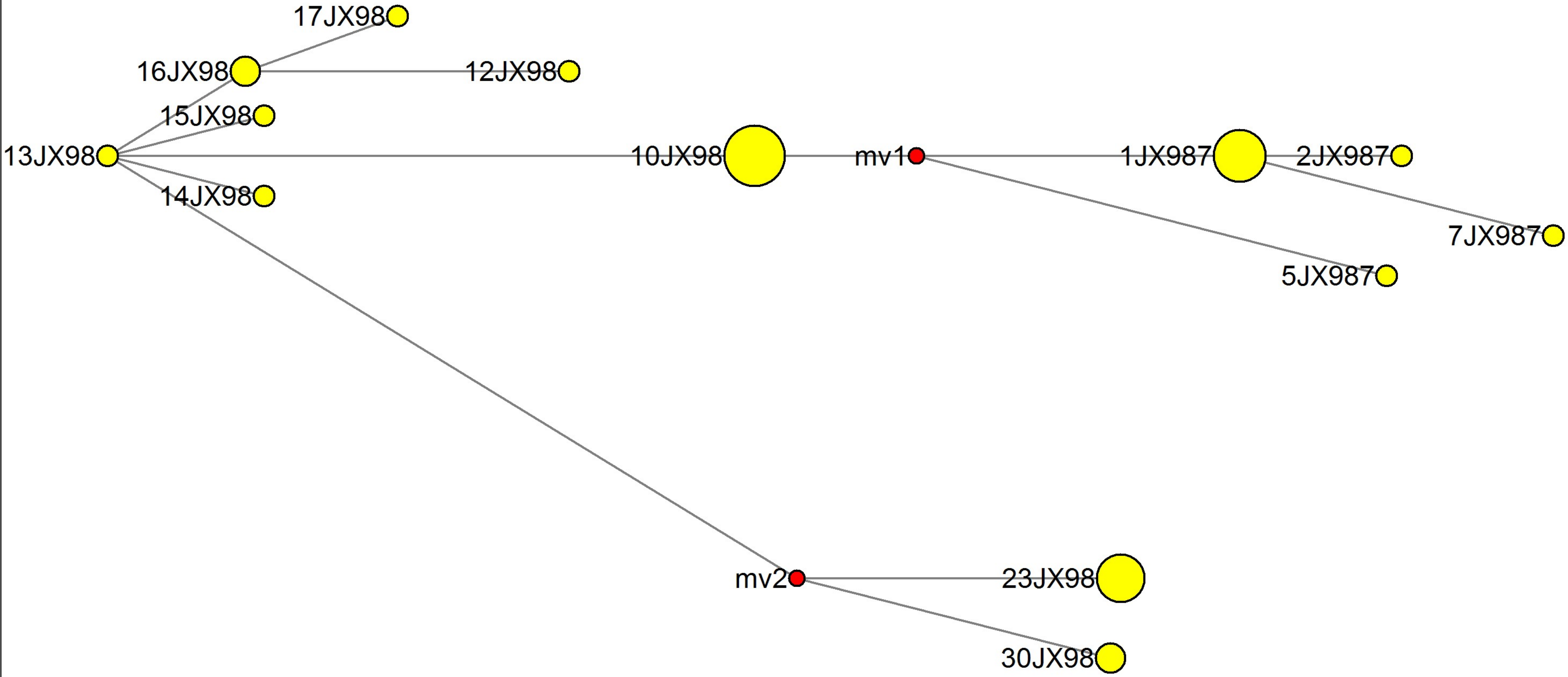

### MJN_case15.pdf

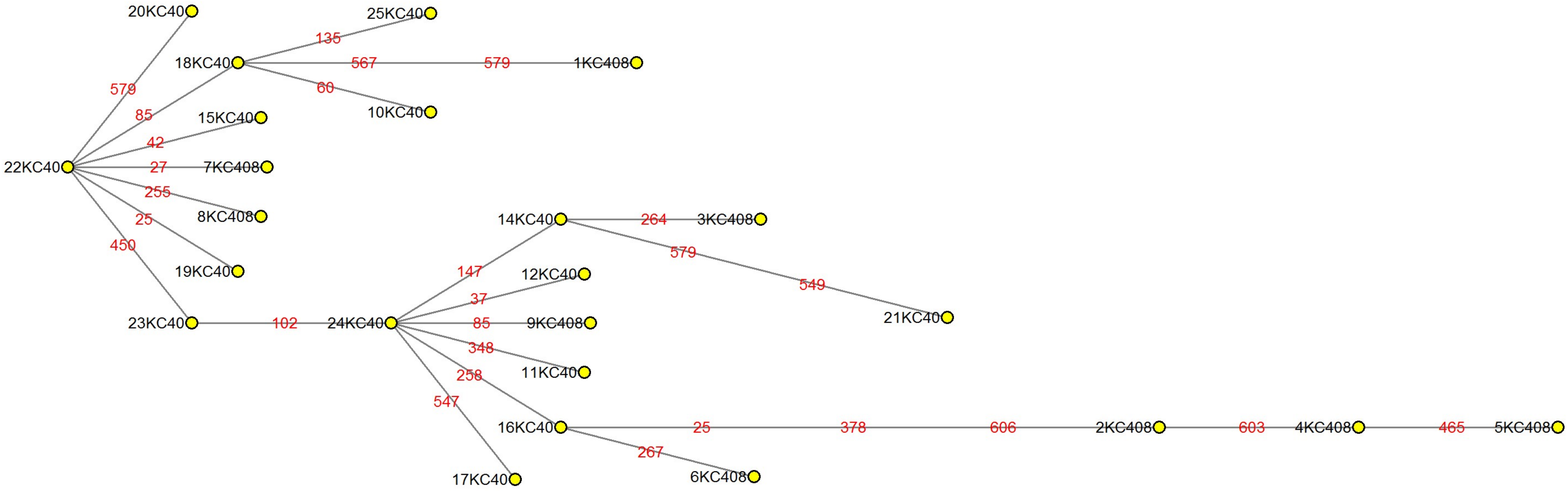

### MJN_case16.pdf

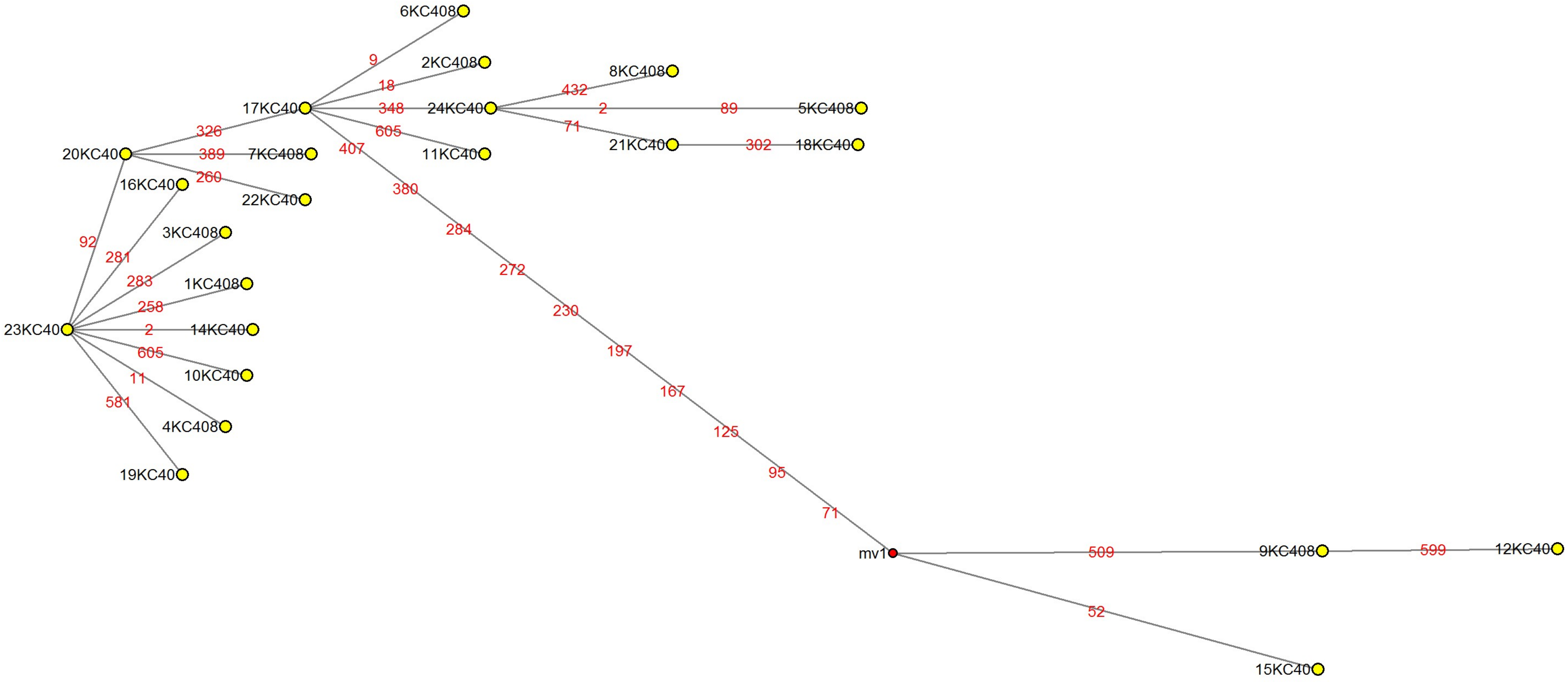

### MJN_case17.pdf

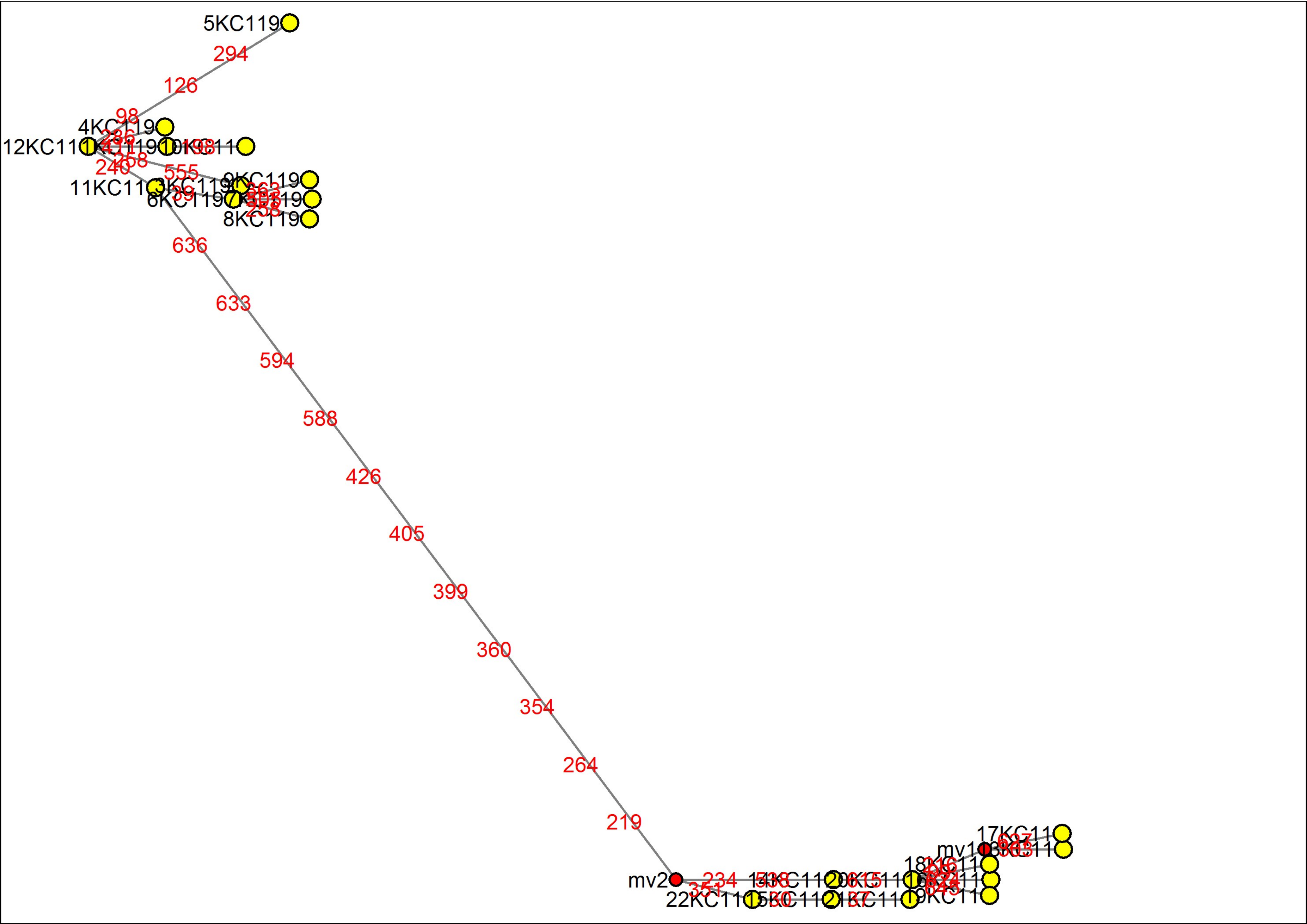

### MJN_case18.pdf

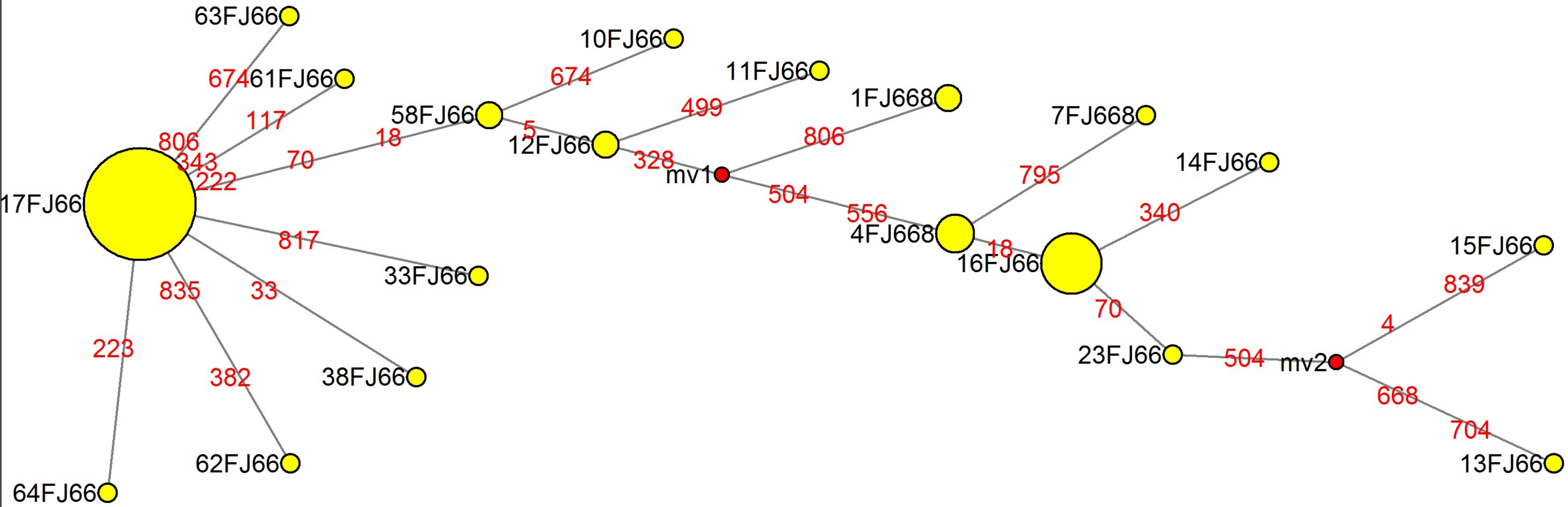

### MJN_case19.pdf

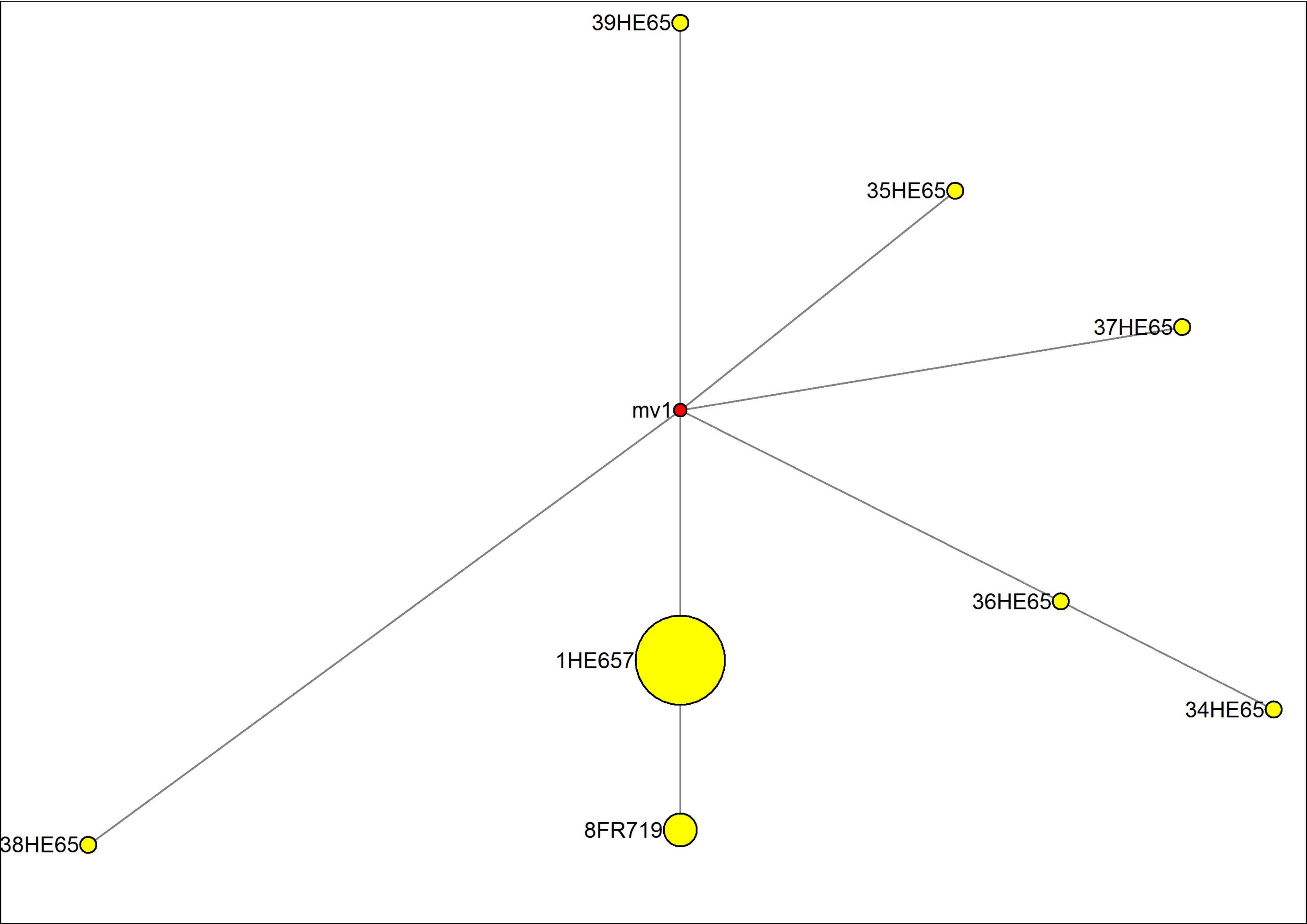

### MJN_case20.pdf

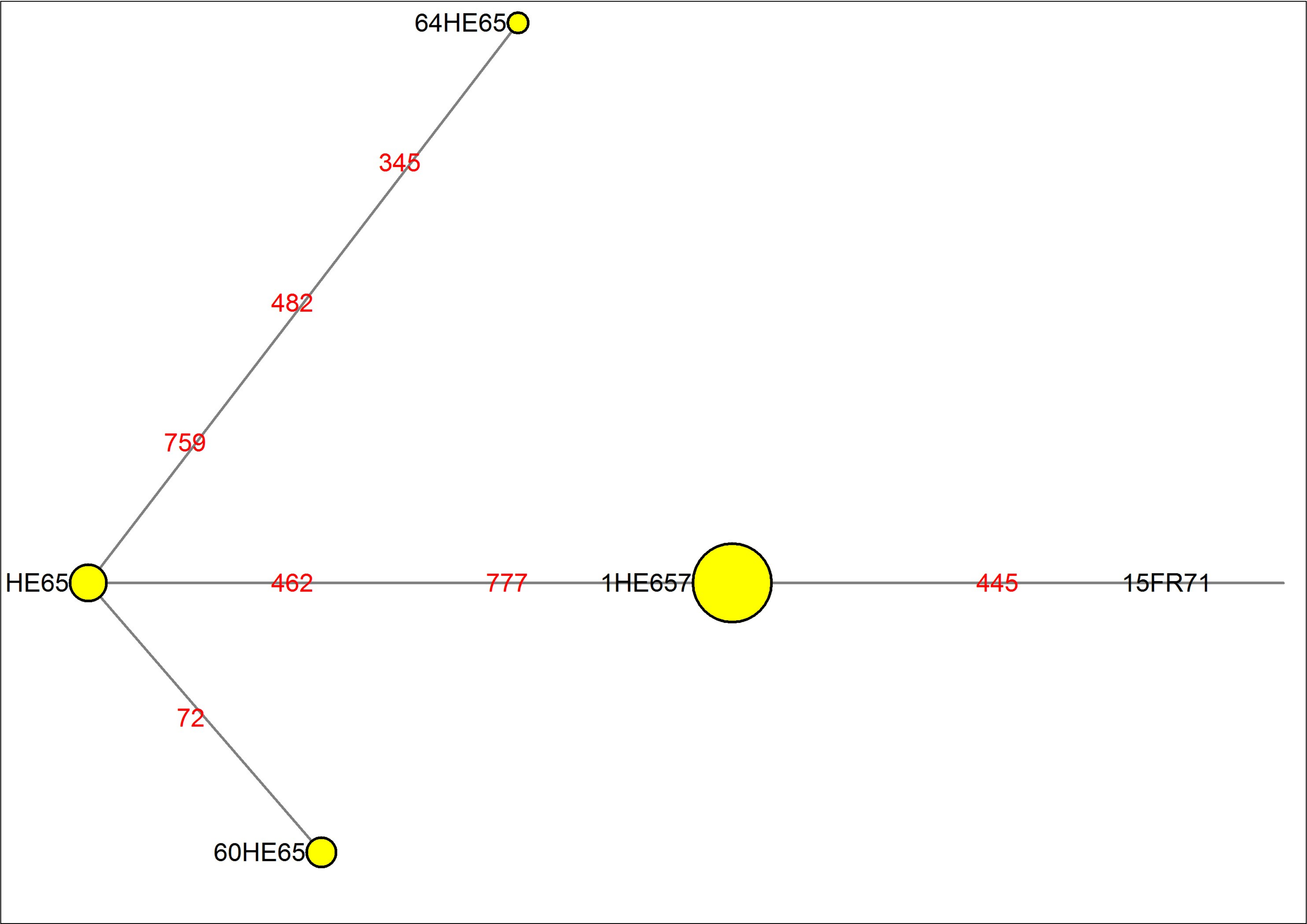

### MJN_case21.pdf

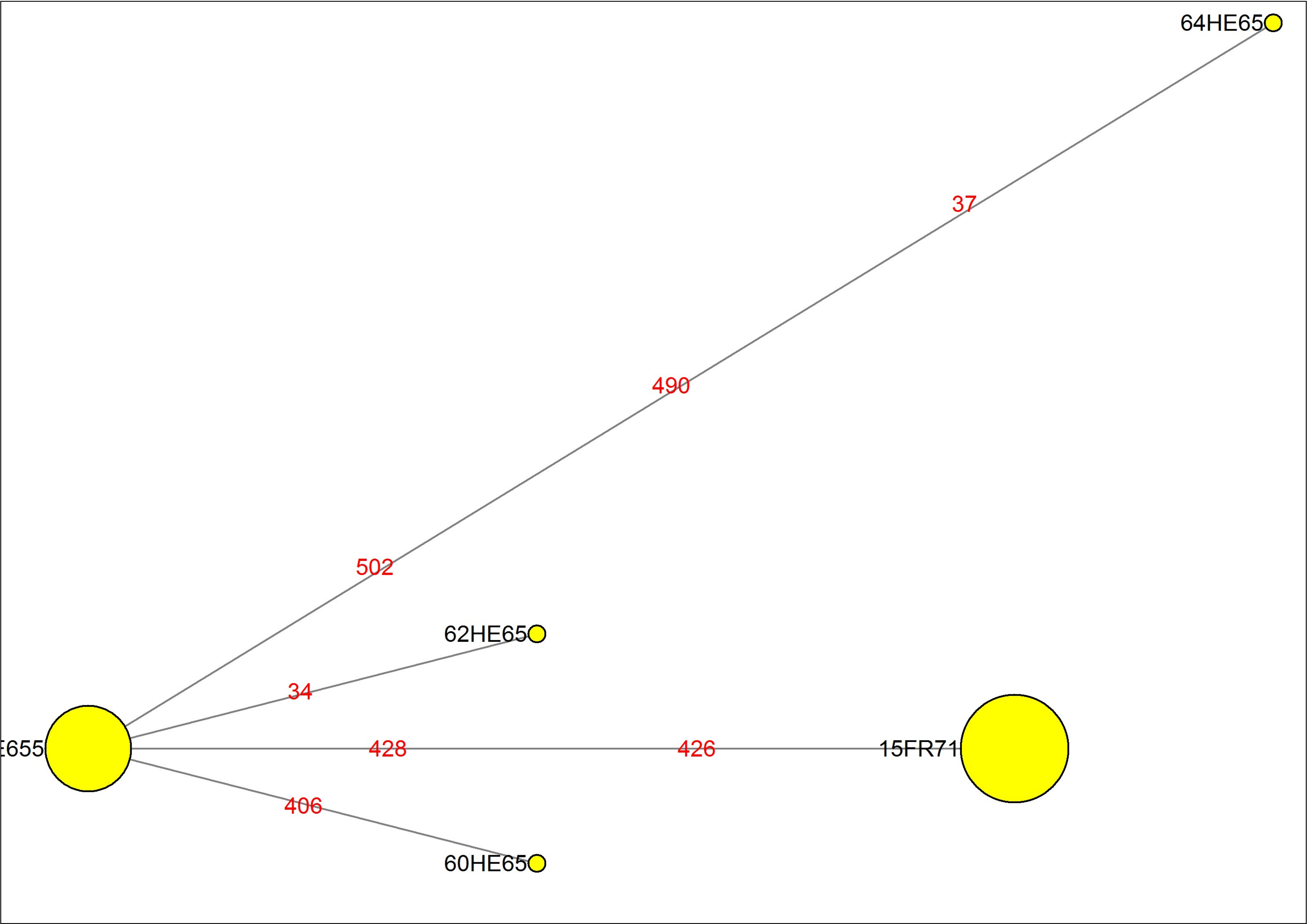

### MJN_case22.pdf

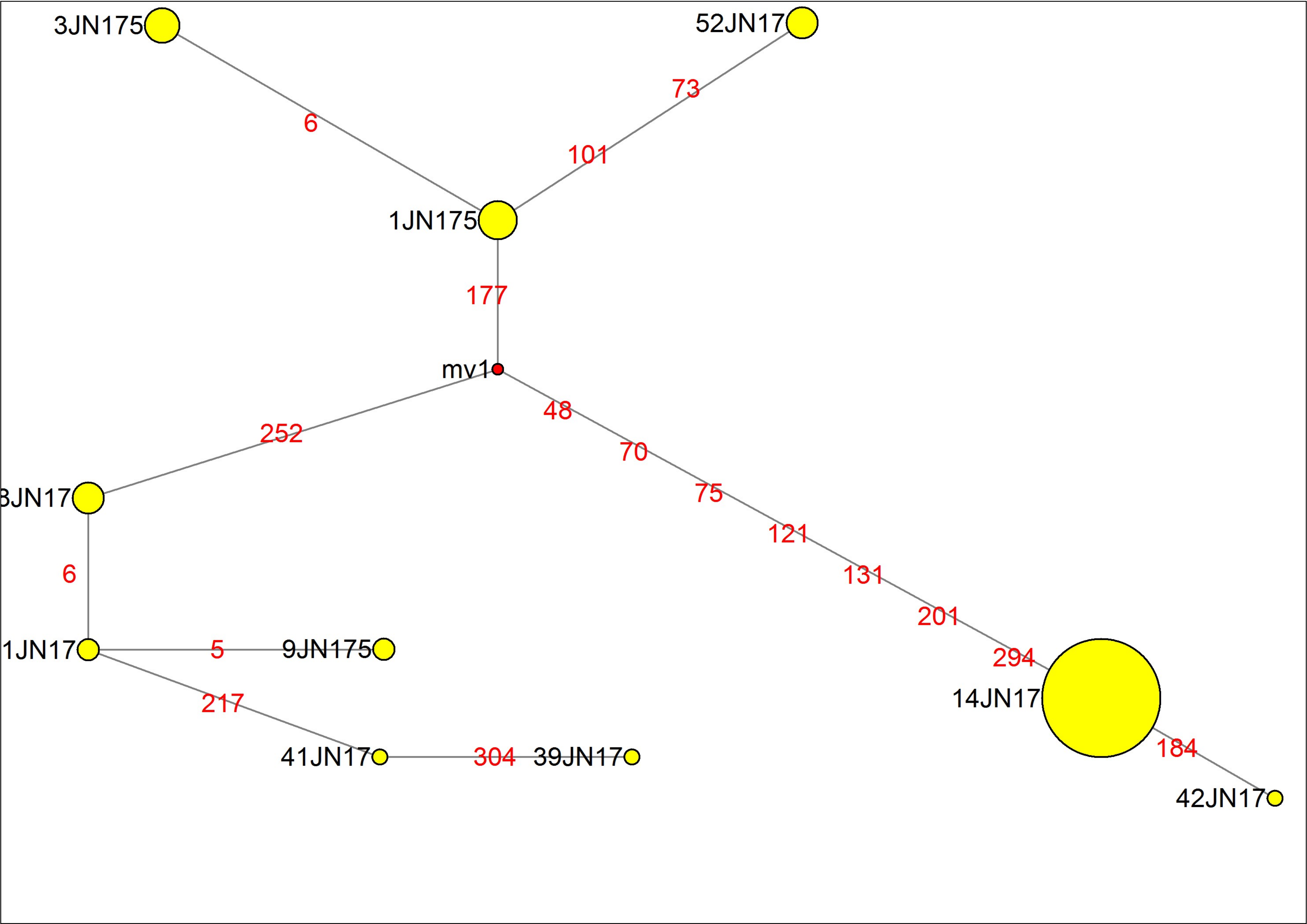

### MJN_case23.pdf

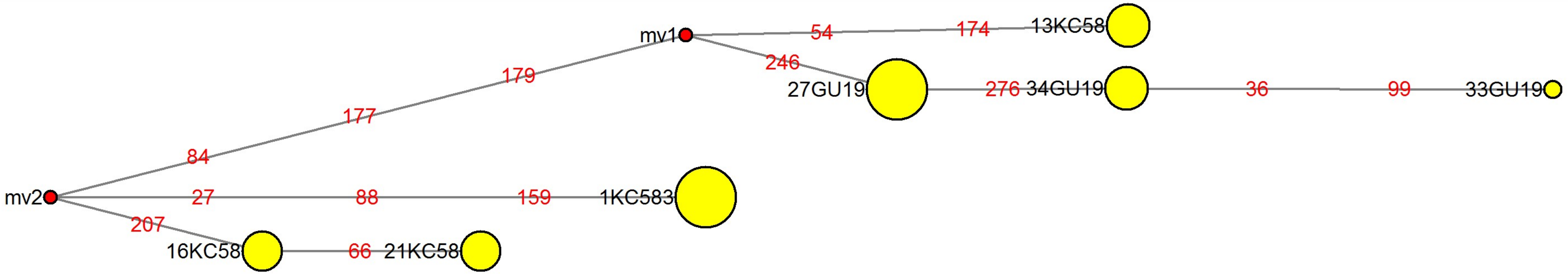

### MJN_case24.pdf

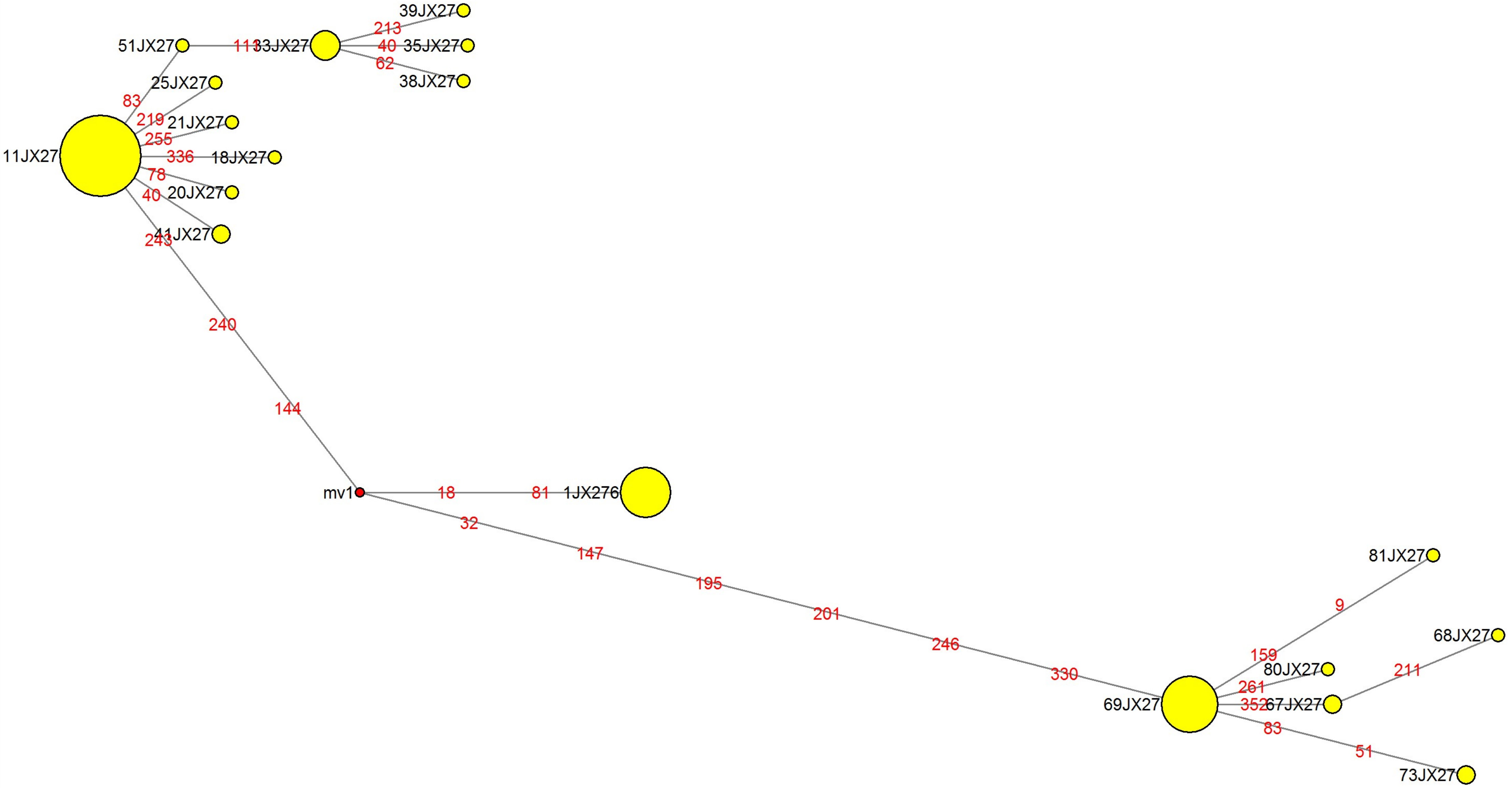

### MJN_case25.pdf

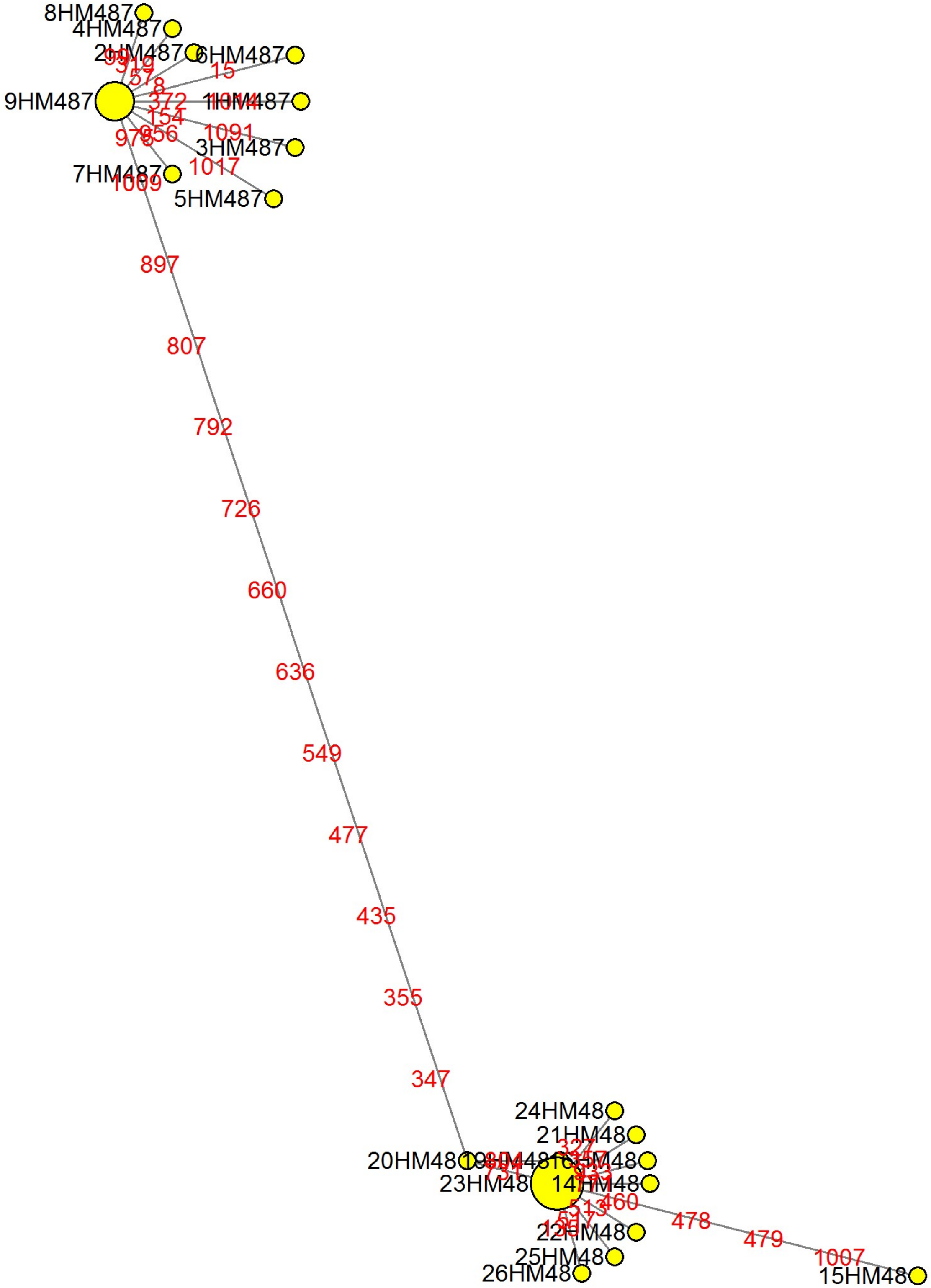

### MJN_case26.pdf

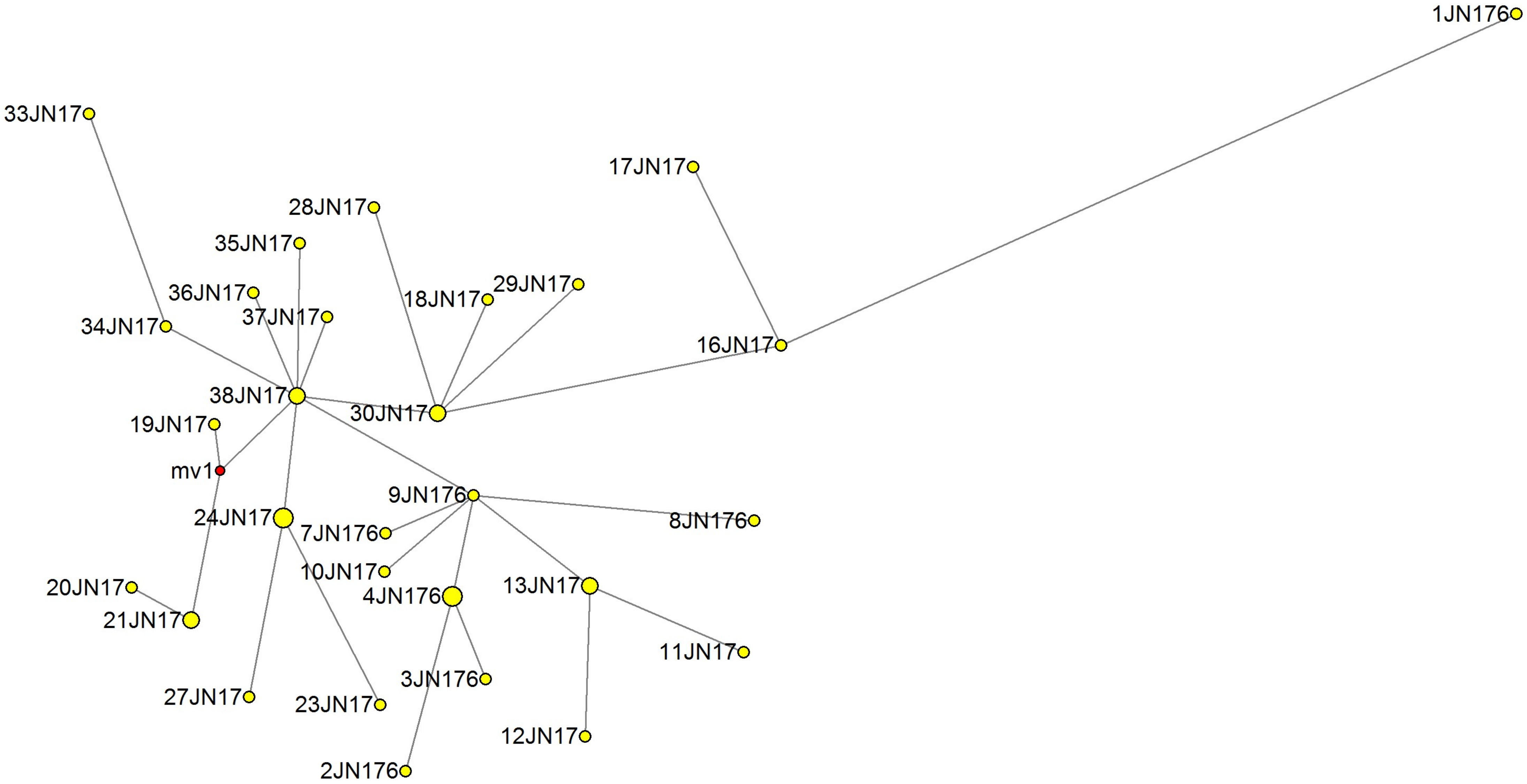

### MJN_case27.pdf

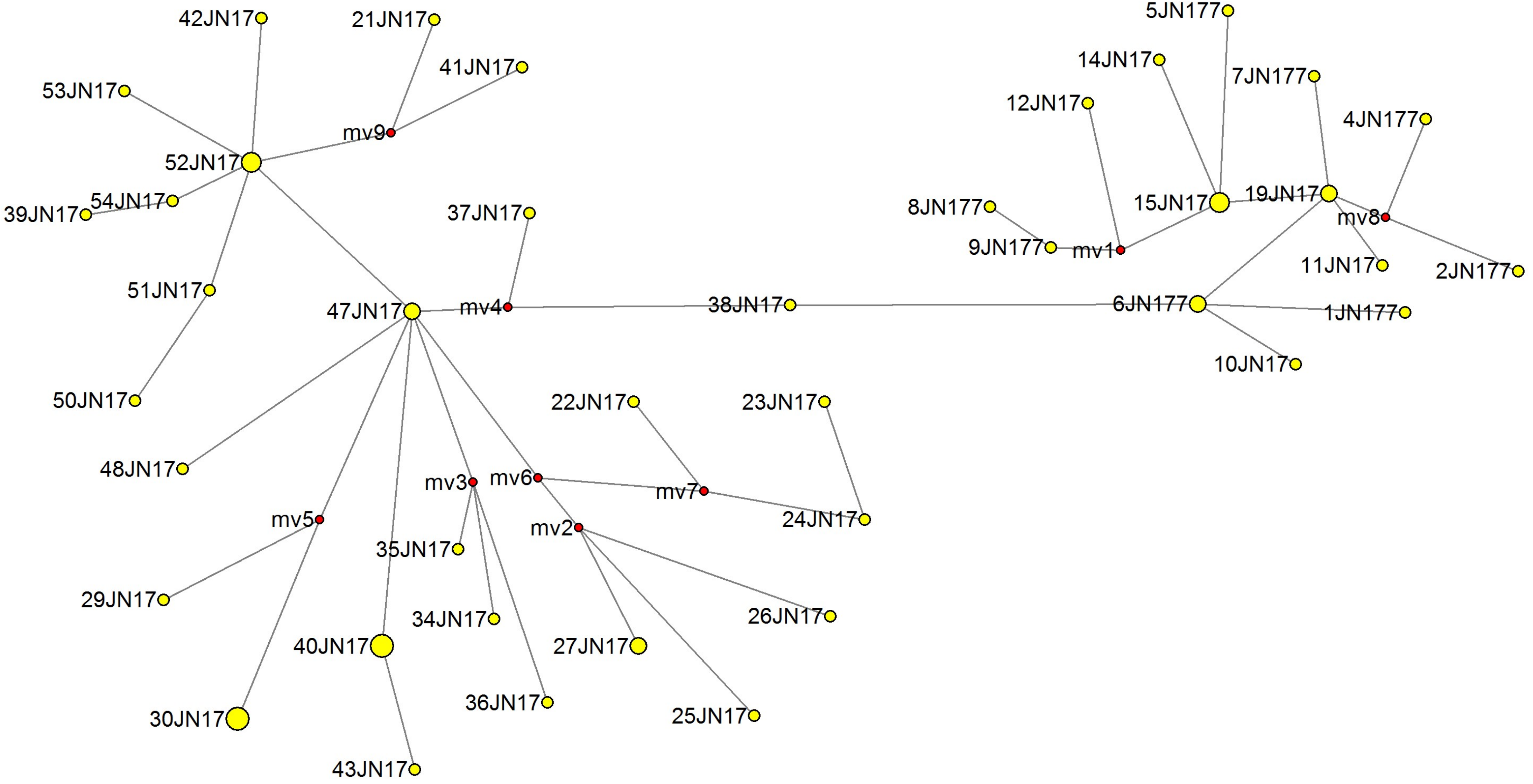

### MJN_case28.pdf

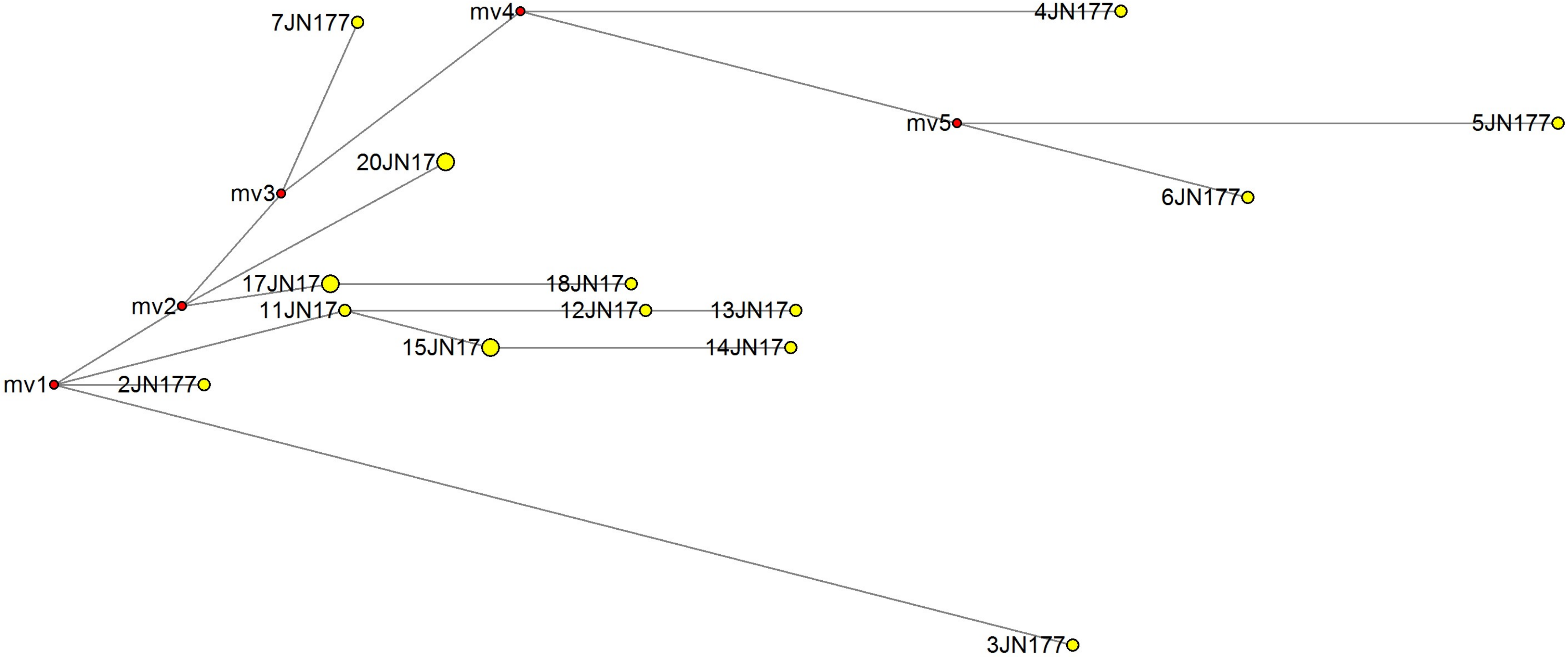

### MJN_case29.pdf

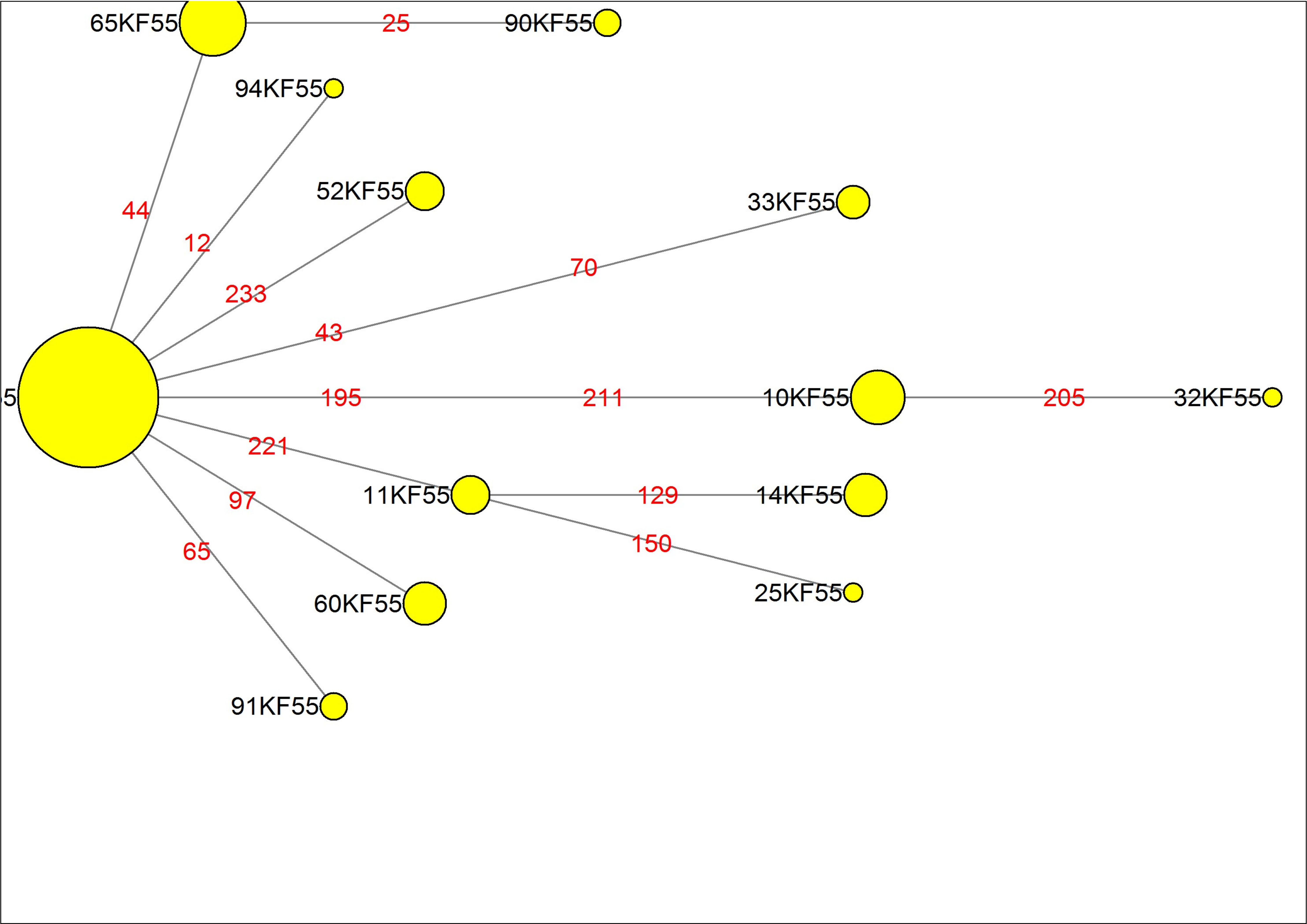

### MJN_case30.pdf

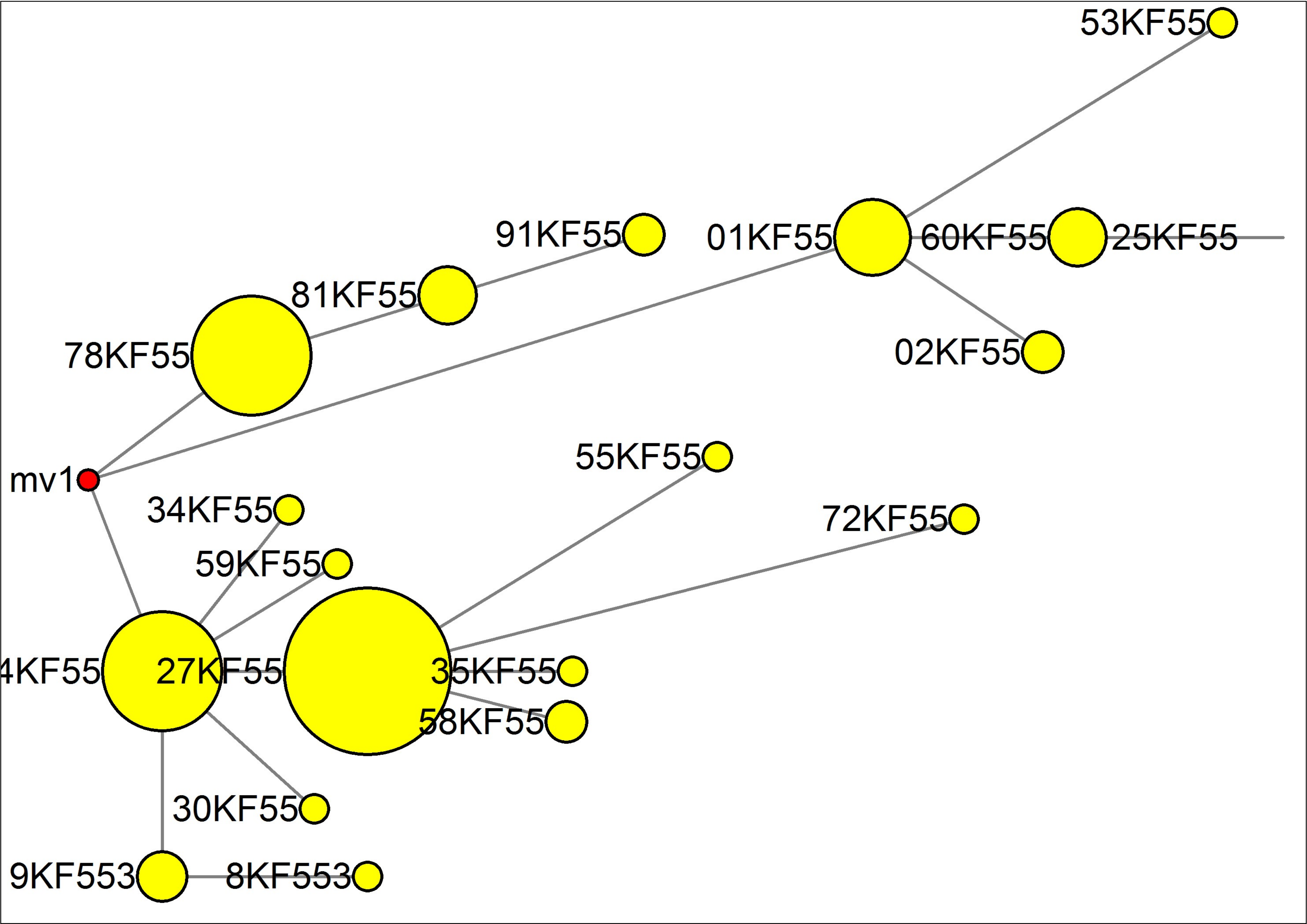

### MJN_case31.pdf

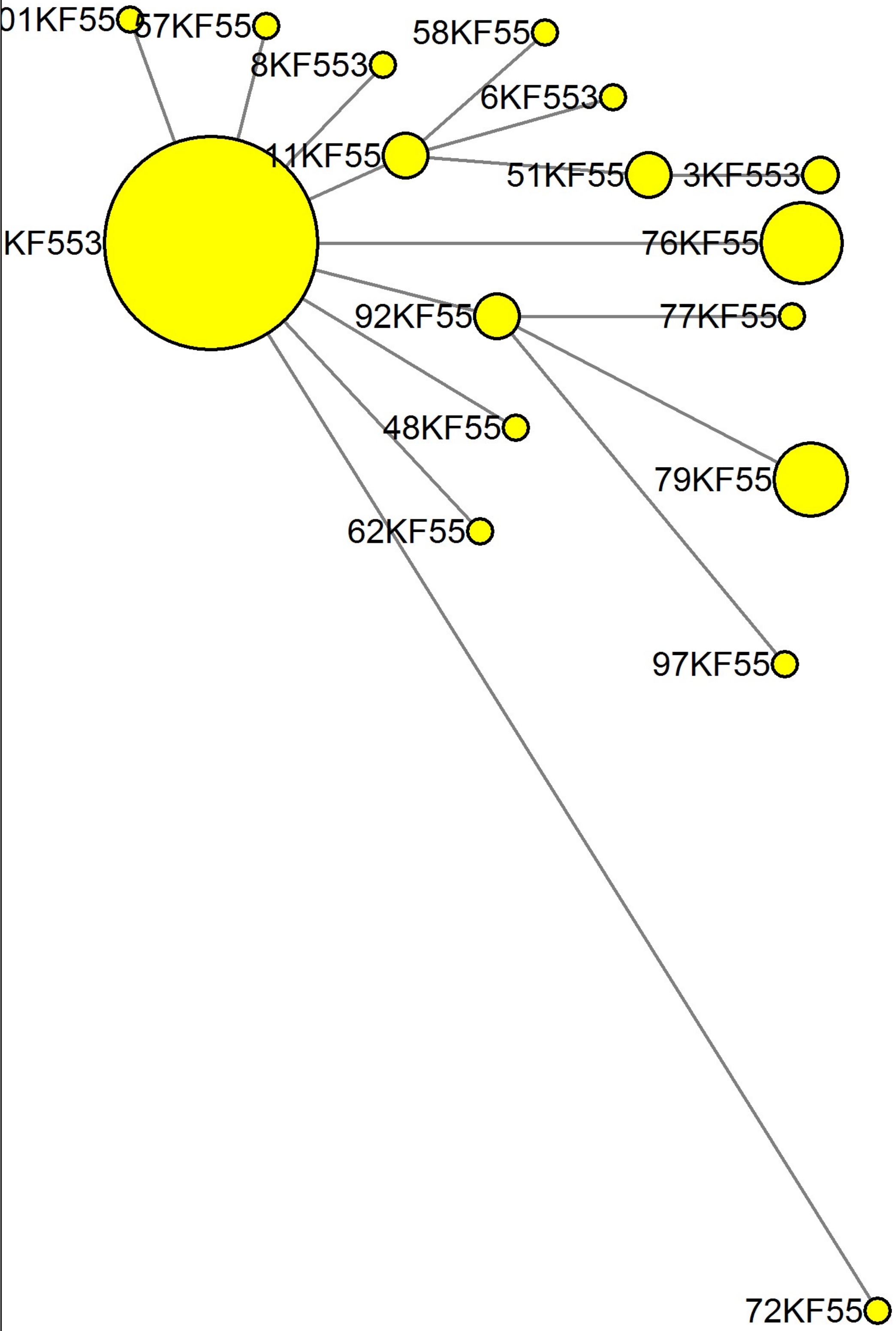

### MJN_case32.pdf

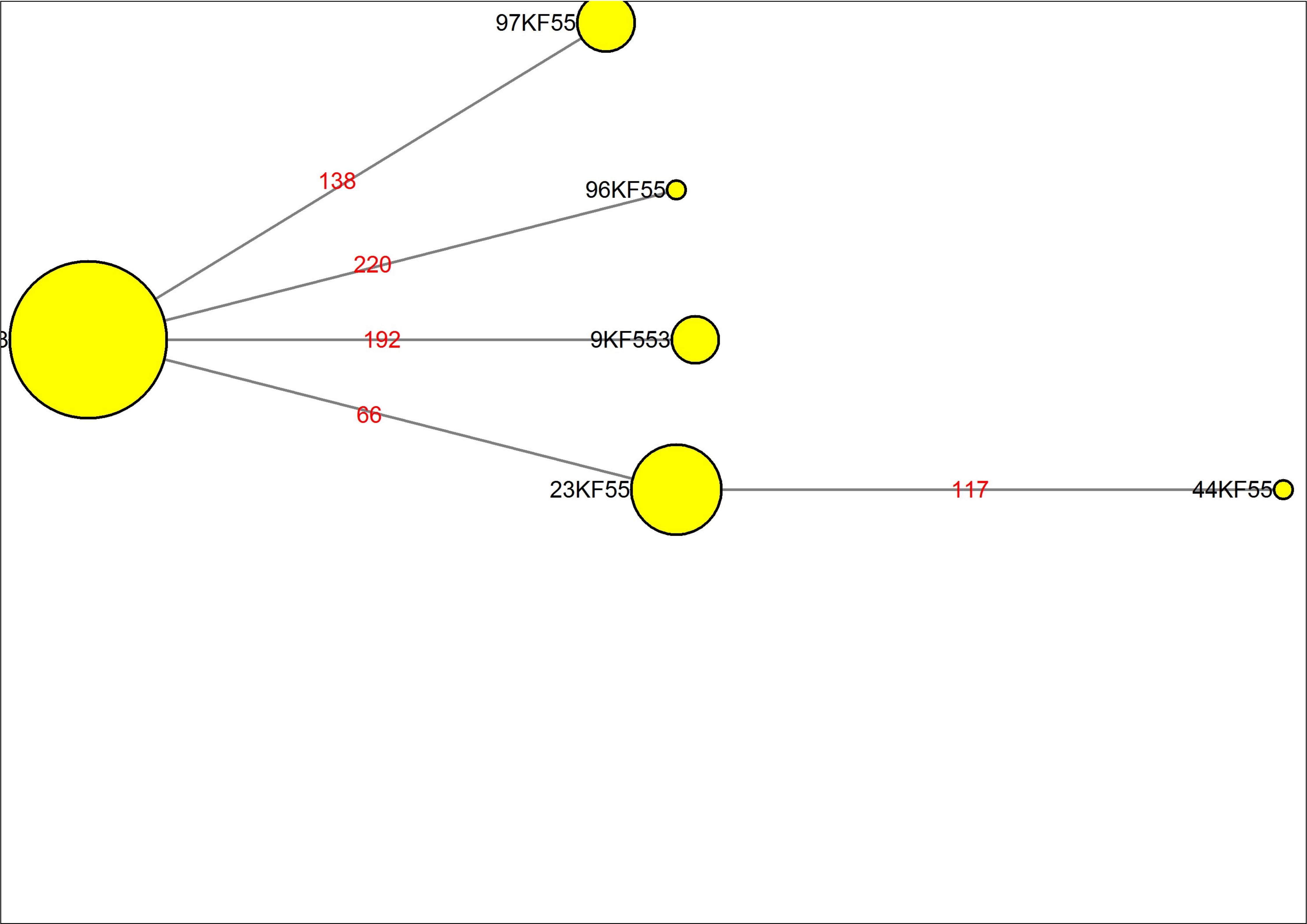

### MJN_case33.pdf

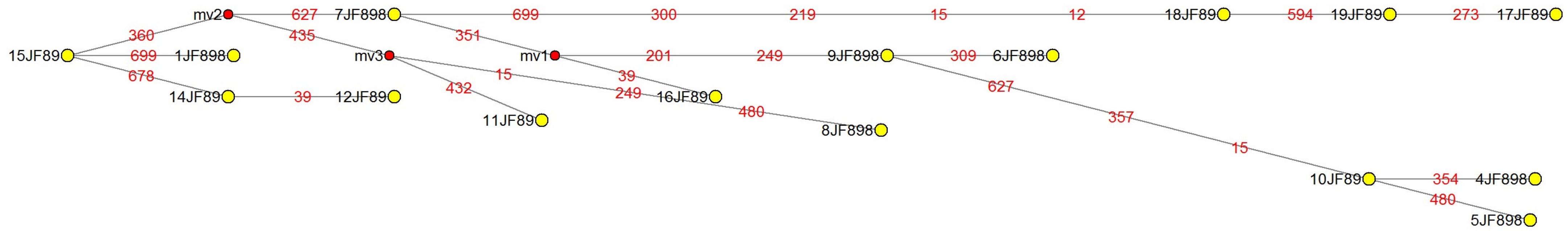

### MJN_case34.pdf

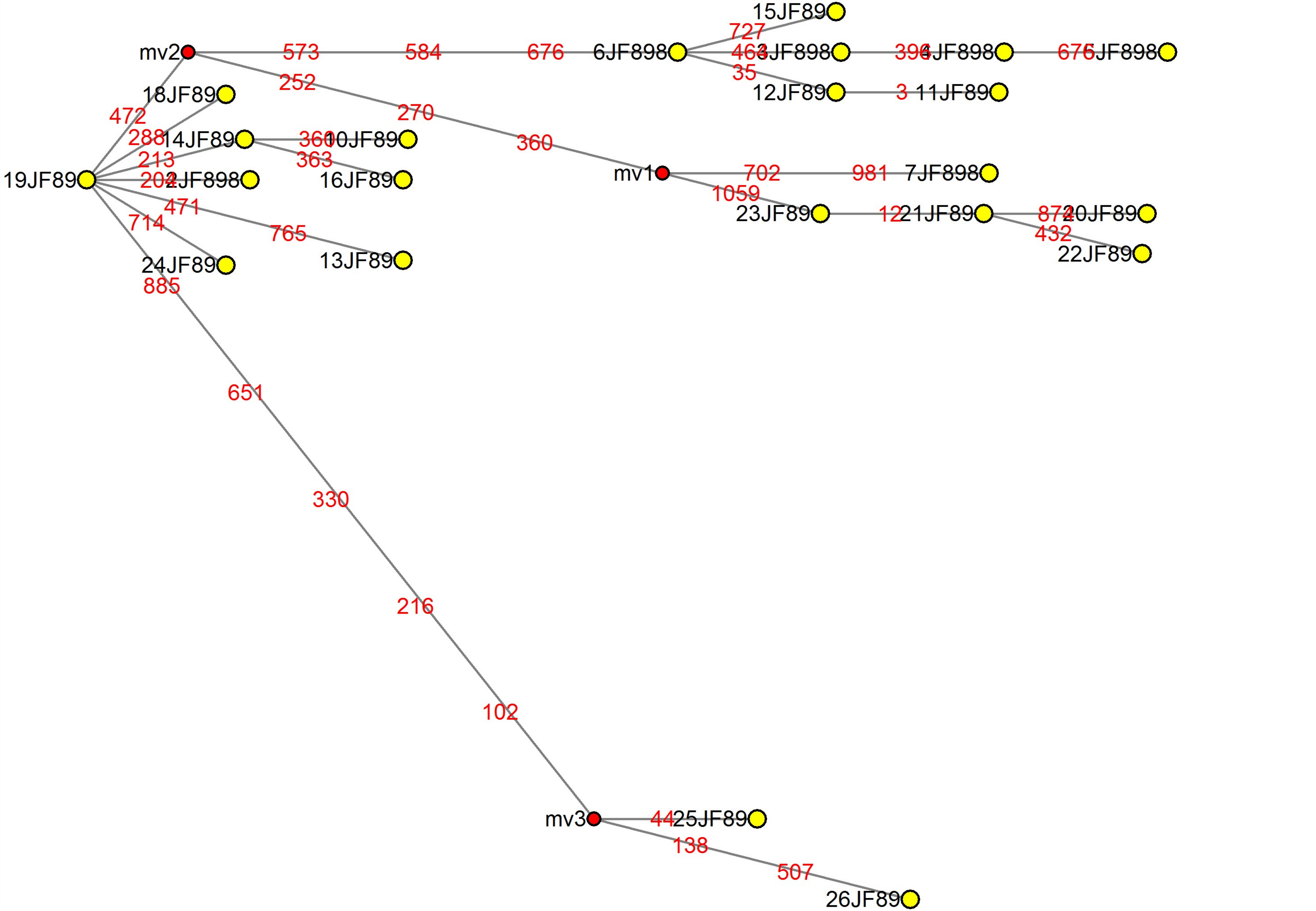

### MJN_case35.pdf

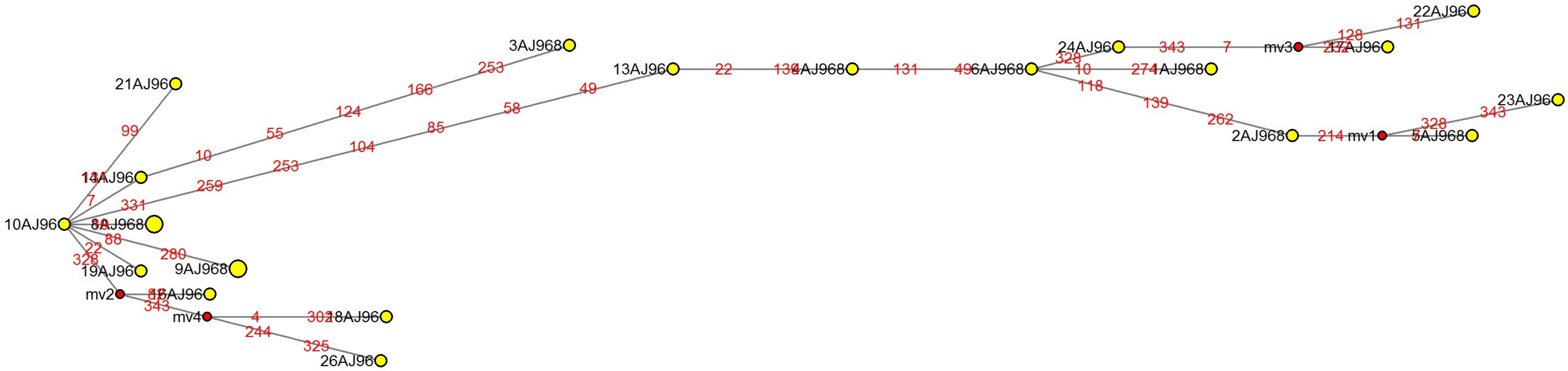

### MJN_case36.pdf

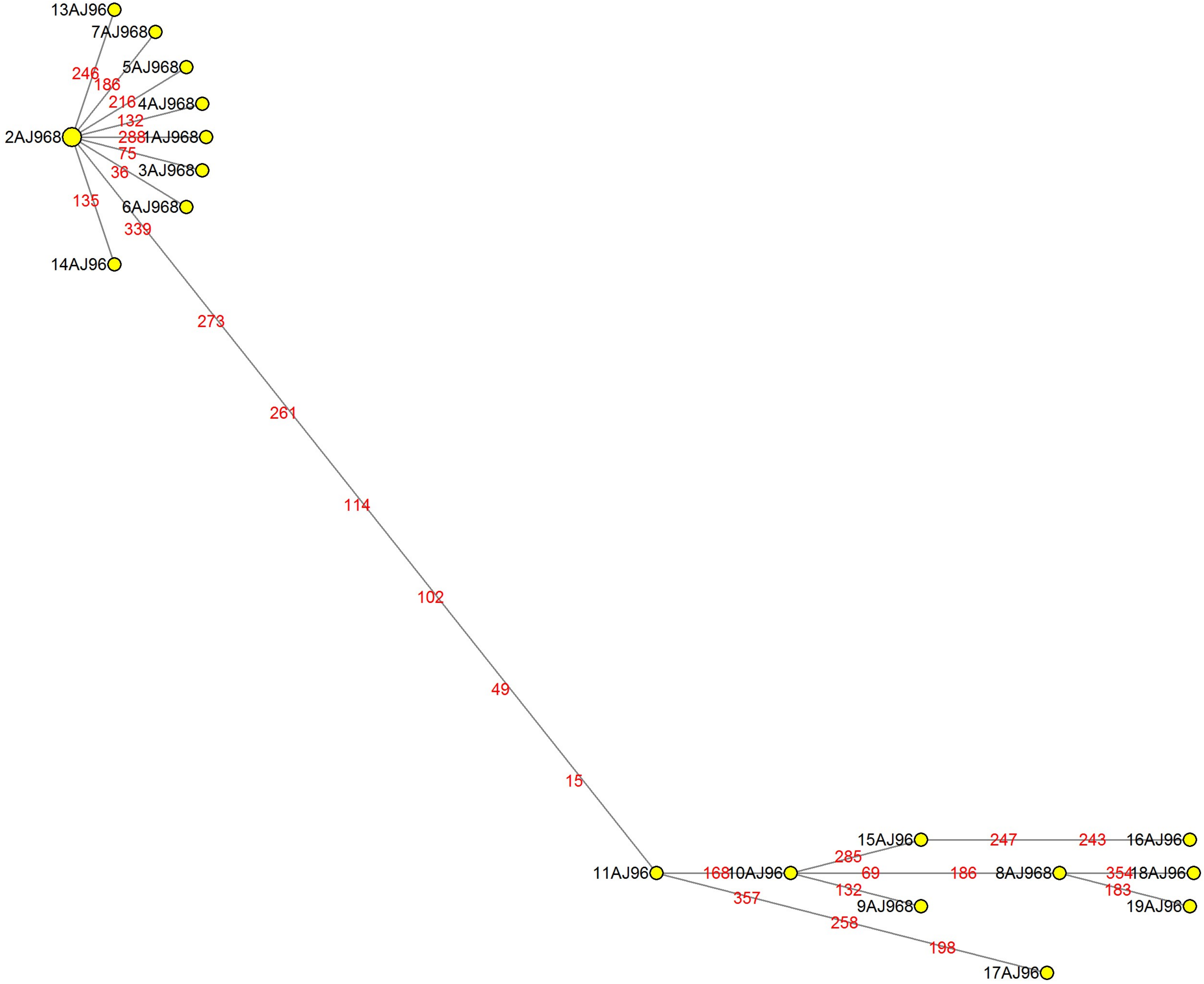

### MJN_case37.pdf

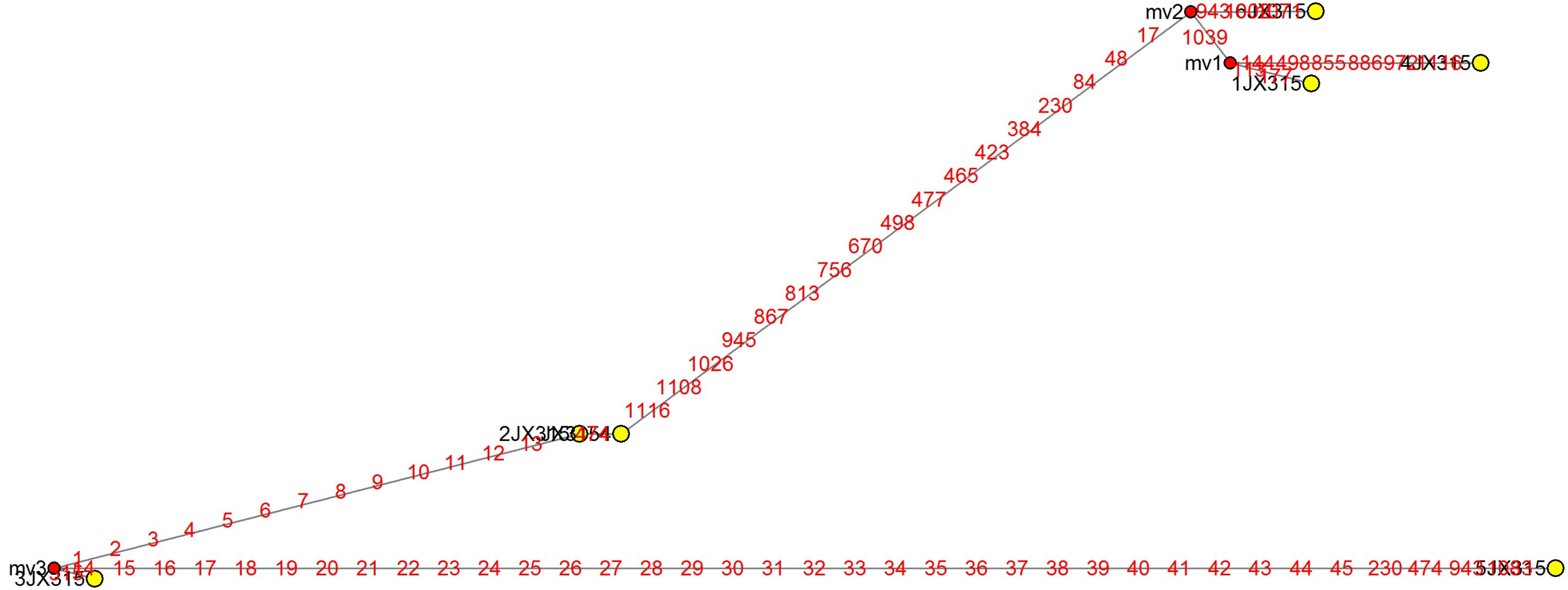

### MJN_case38.pdf

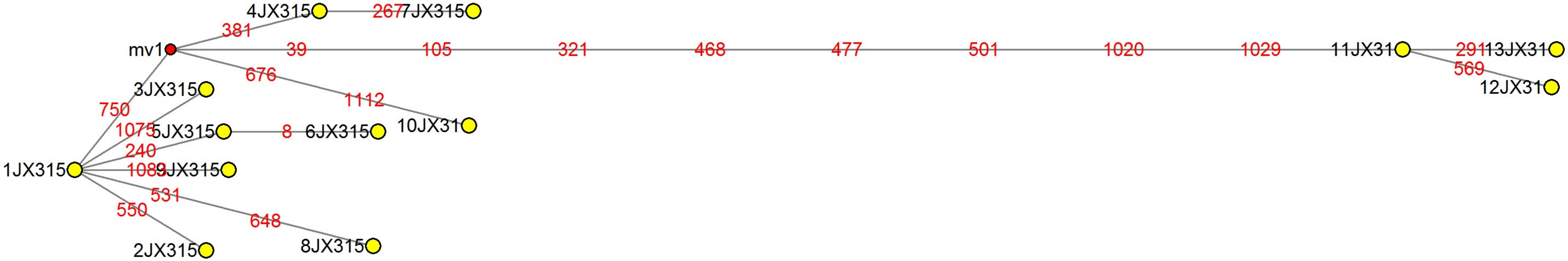

### MJN_case39.pdf

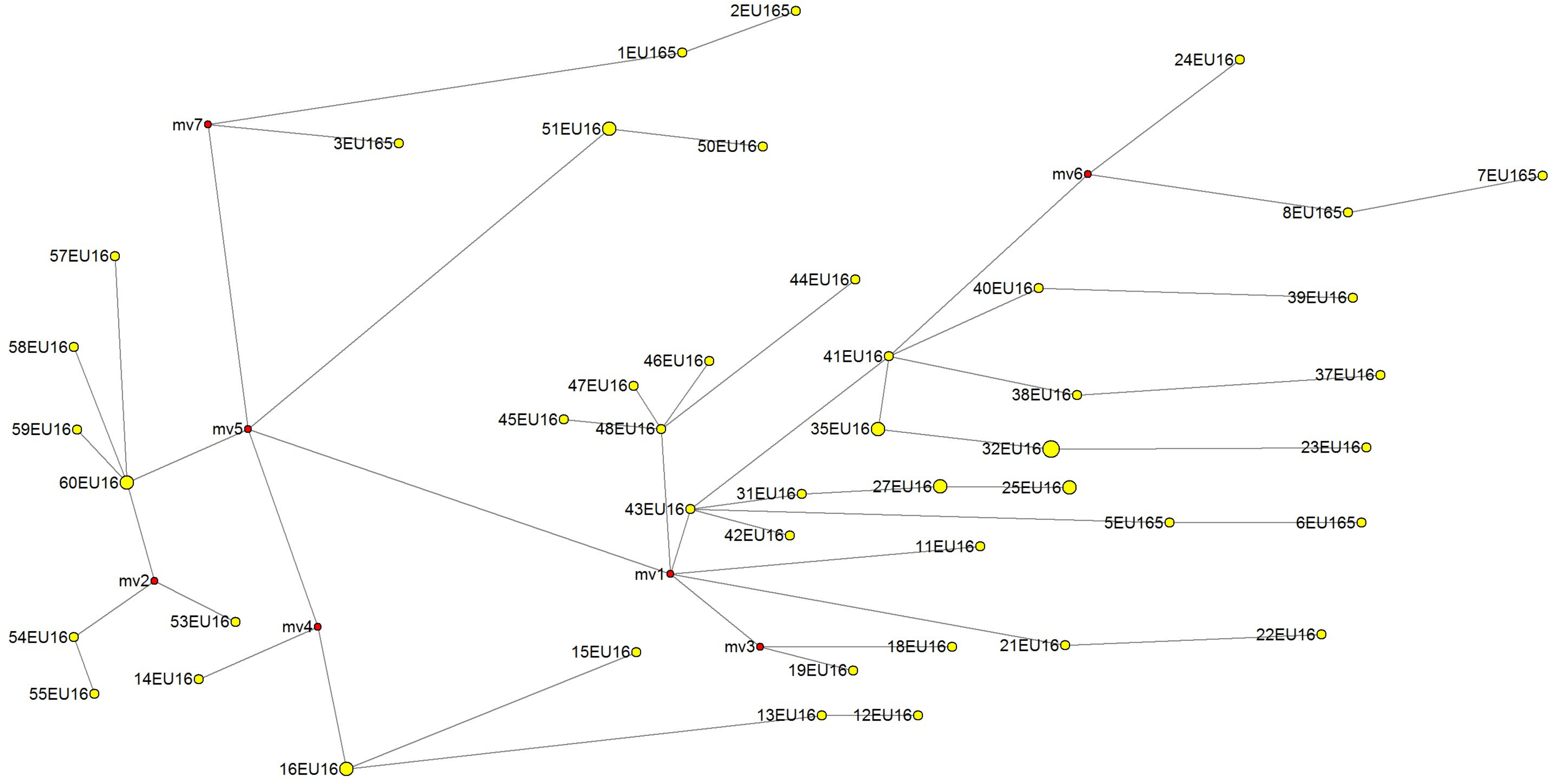

### MJN_case40.pdf

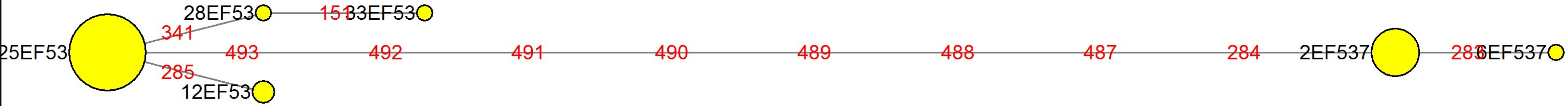

### MJN_case41.pdf

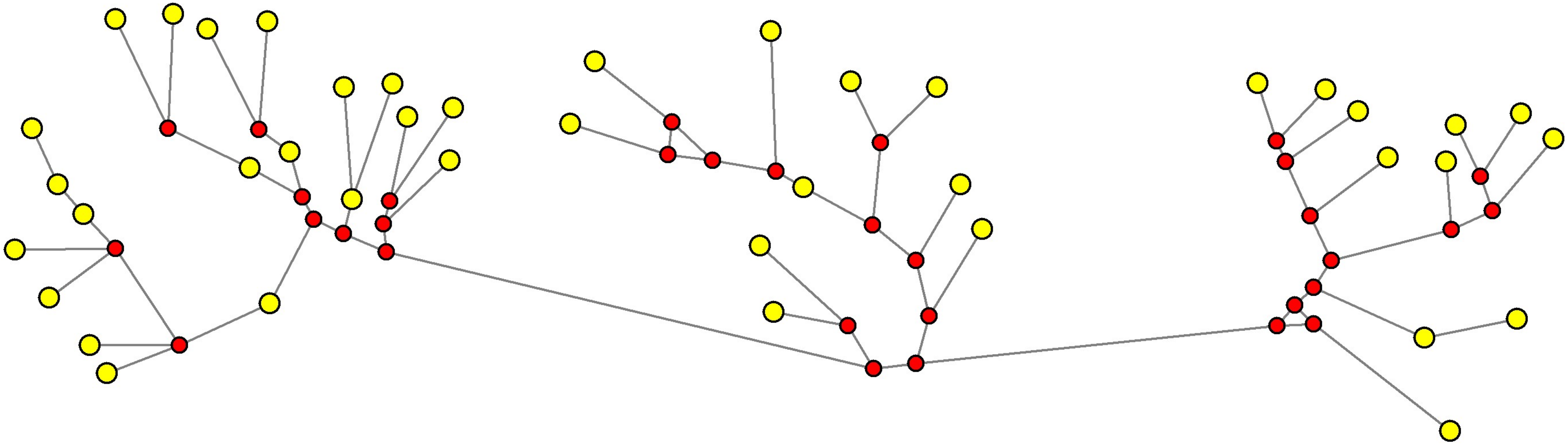

### MJN_case42.pdf

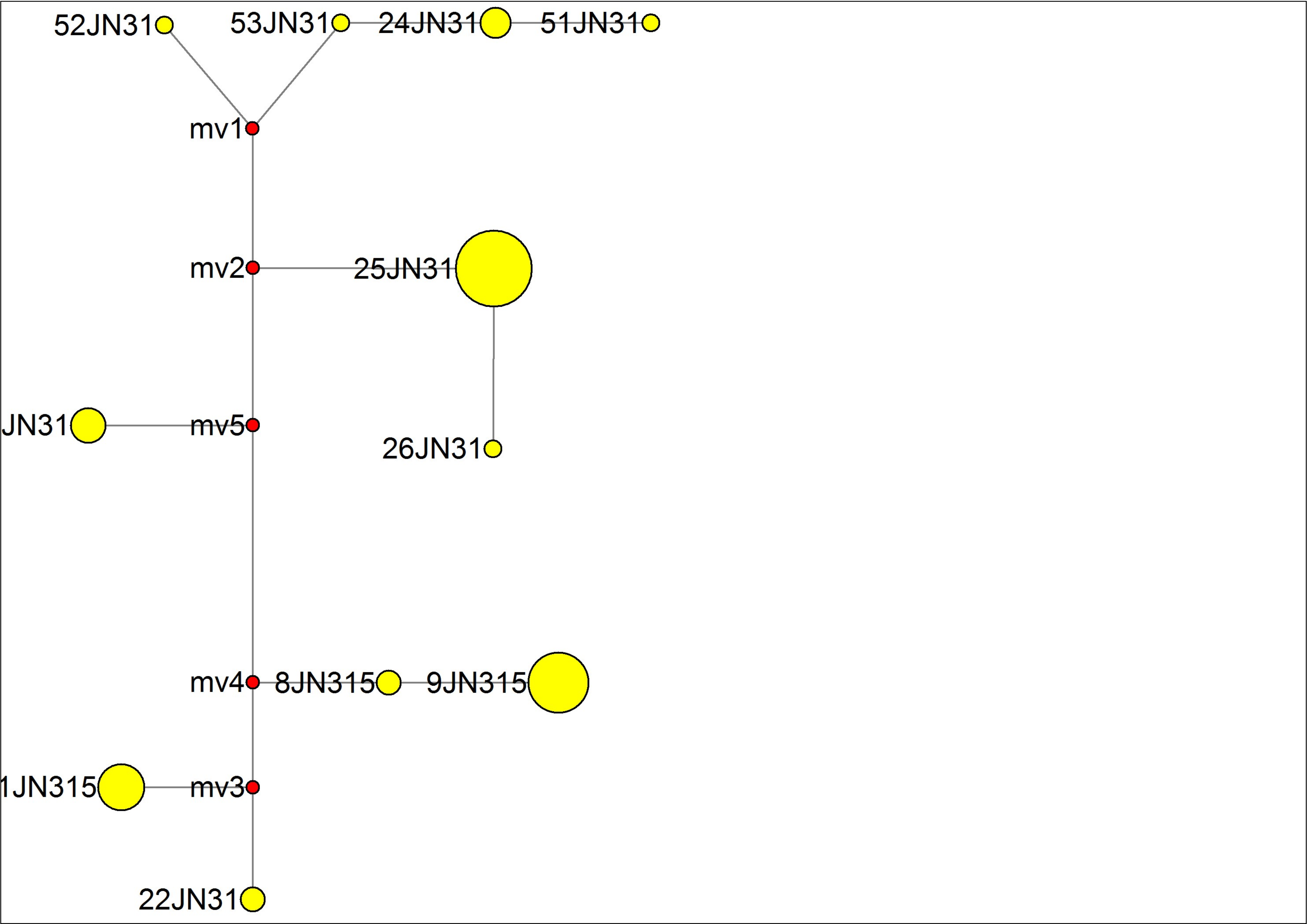

### MJN_case43.pdf

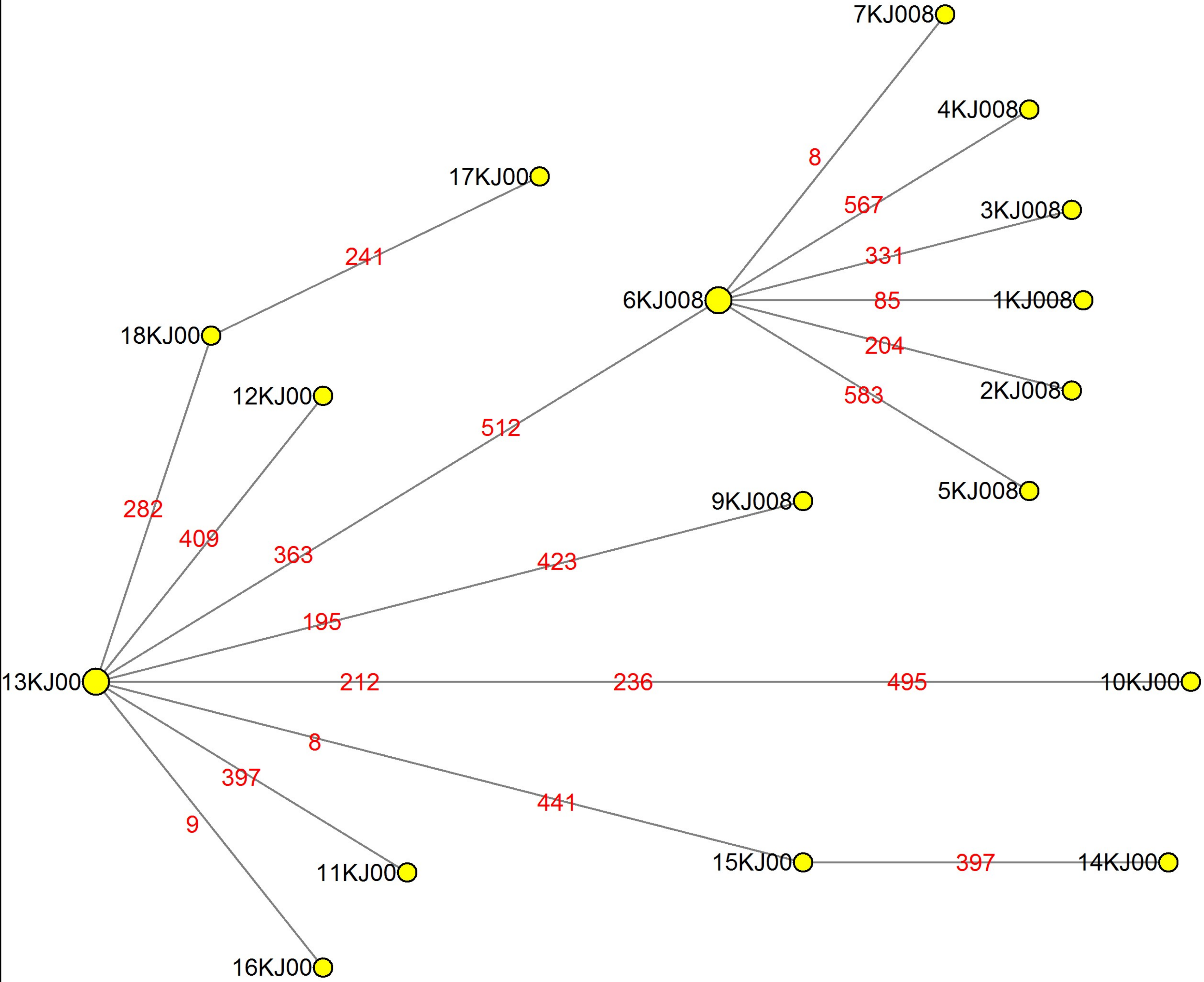

### MJN_case44.pdf

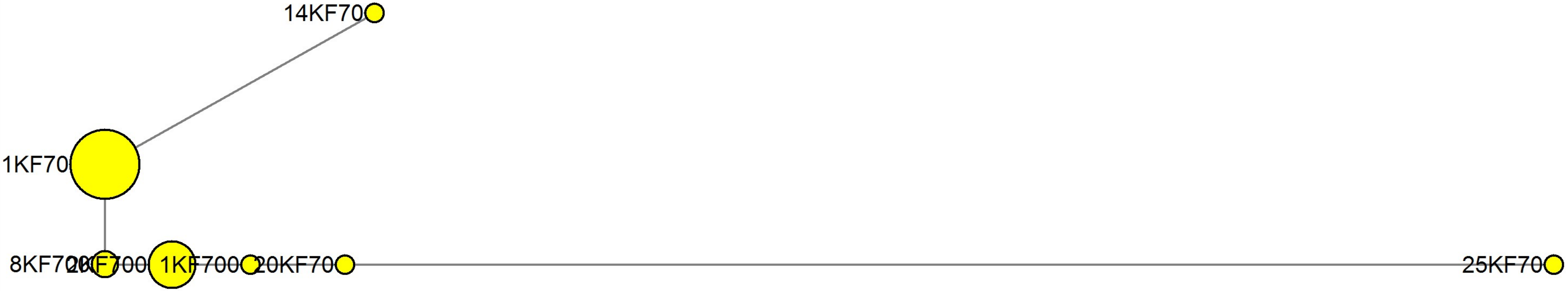

### MJN_case45.pdf

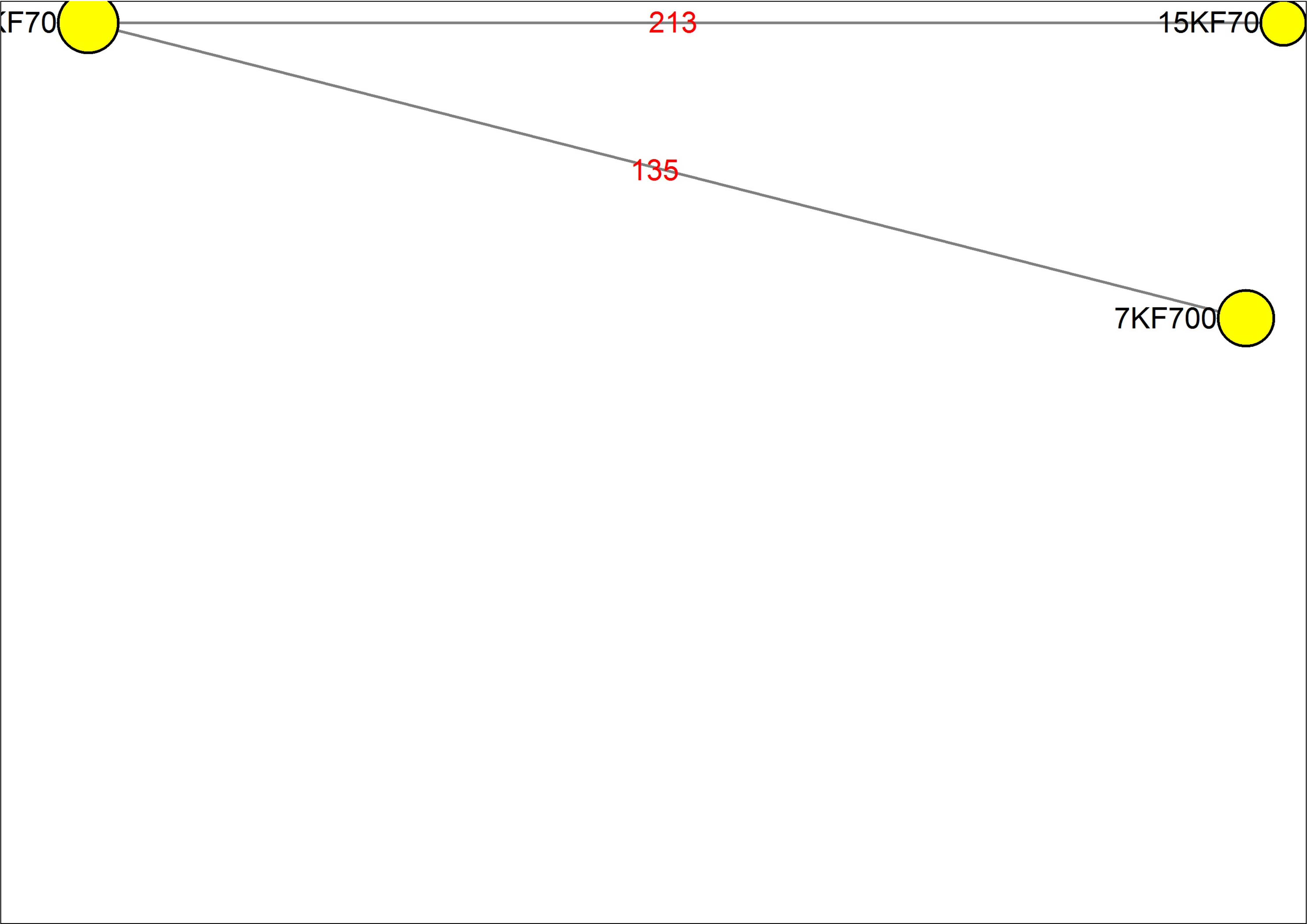

### MJN_case46.pdf

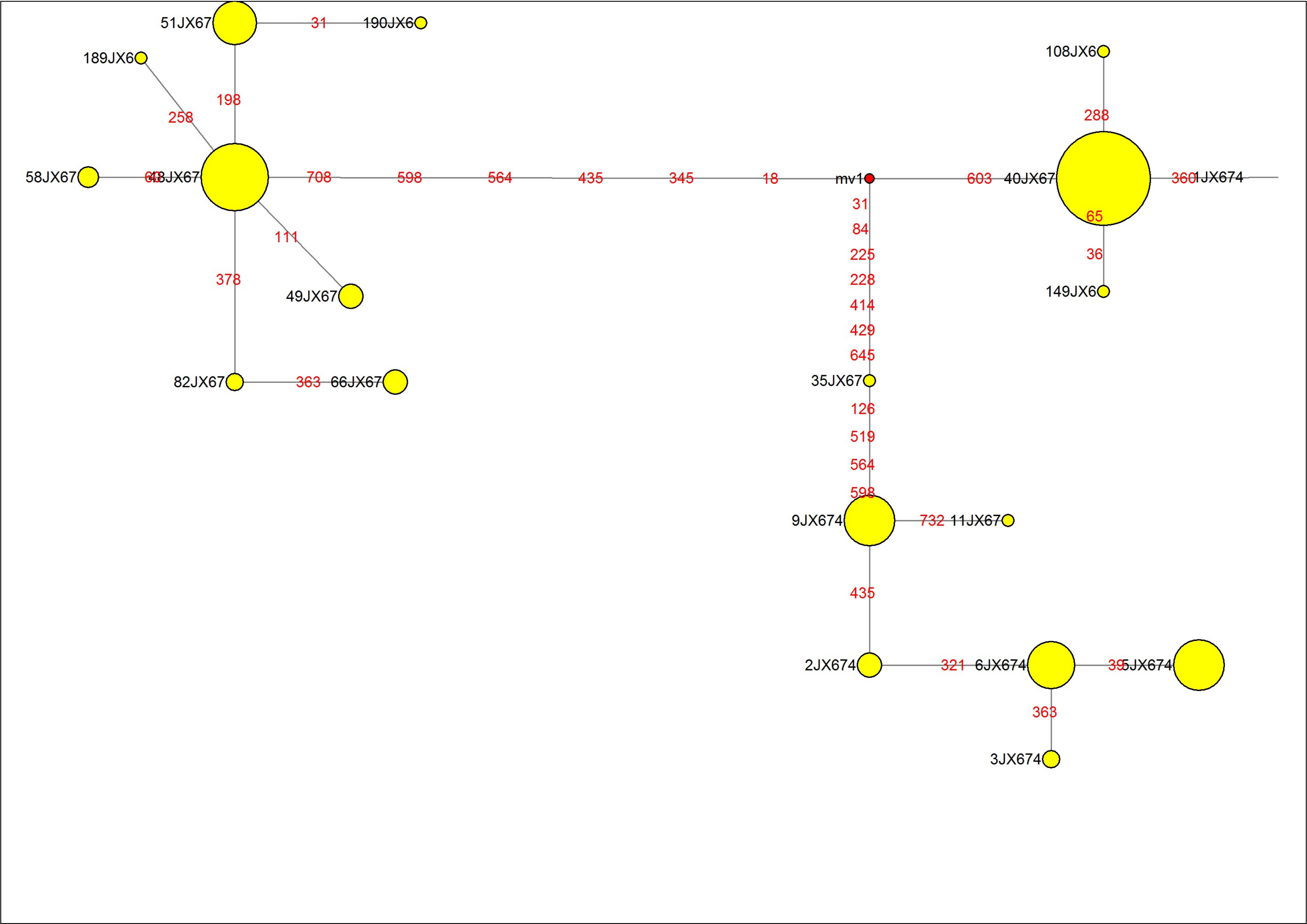

### MJN_case47.pdf

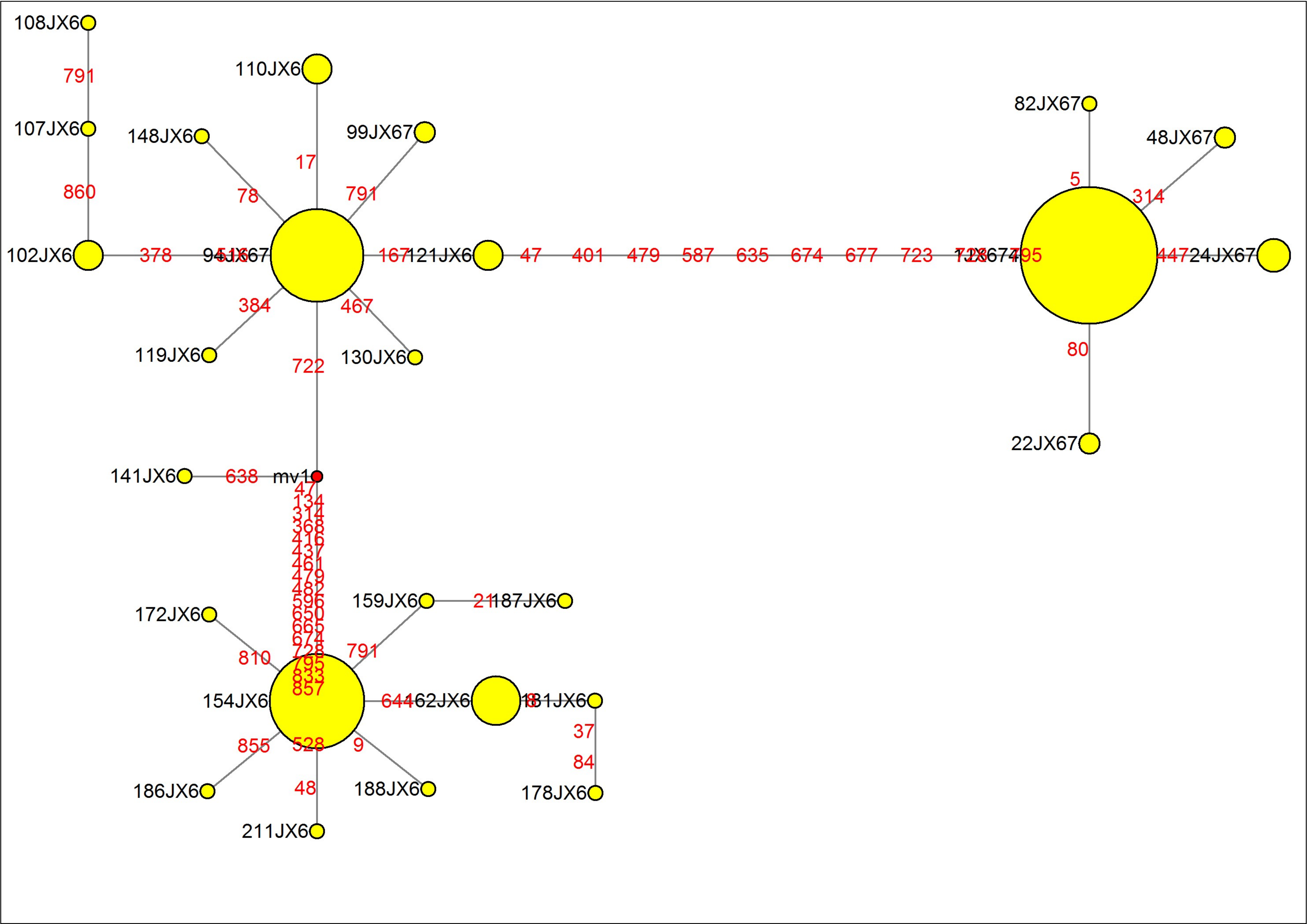

### MJN_case48.pdf

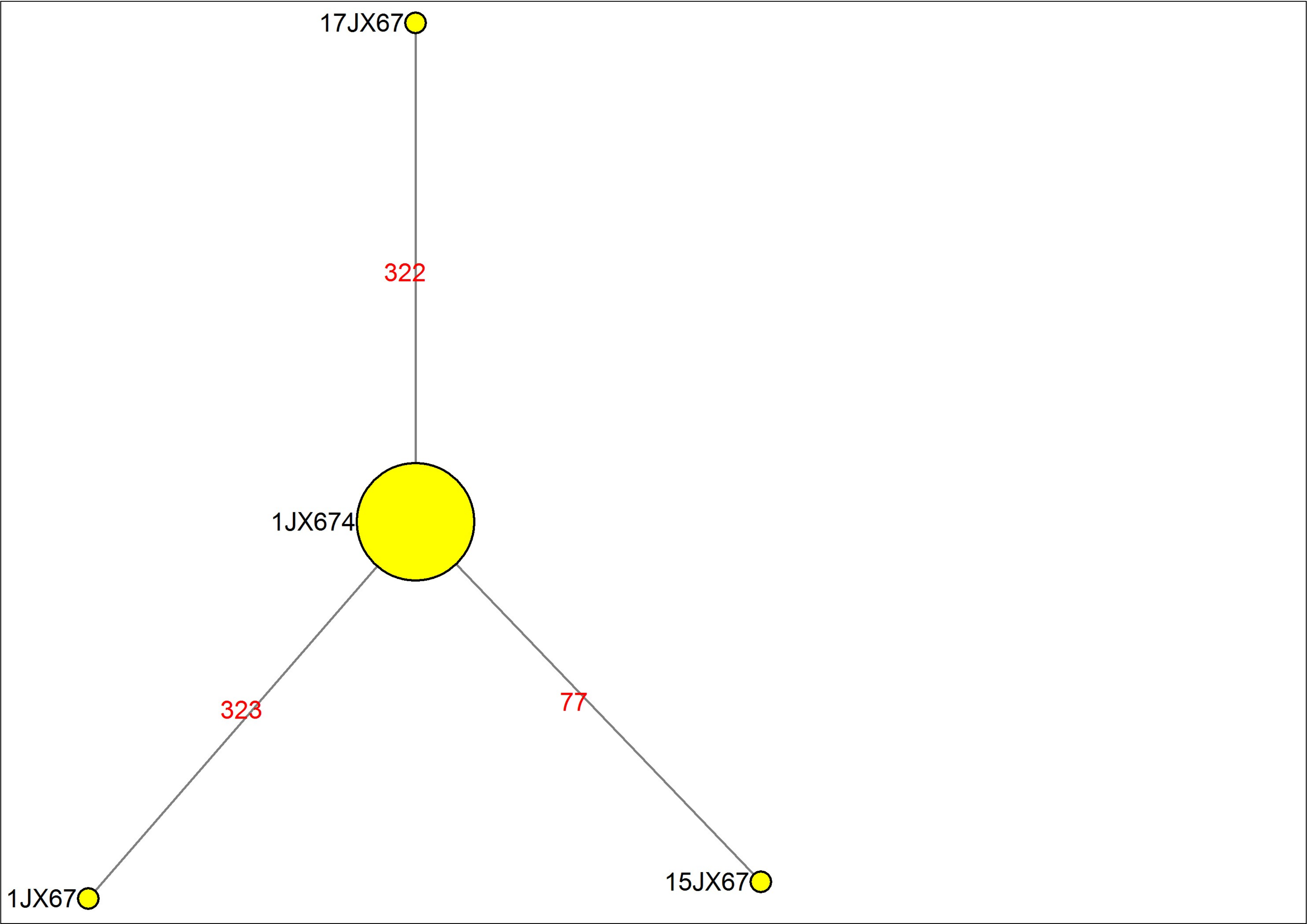

### MJN_case49.pdf

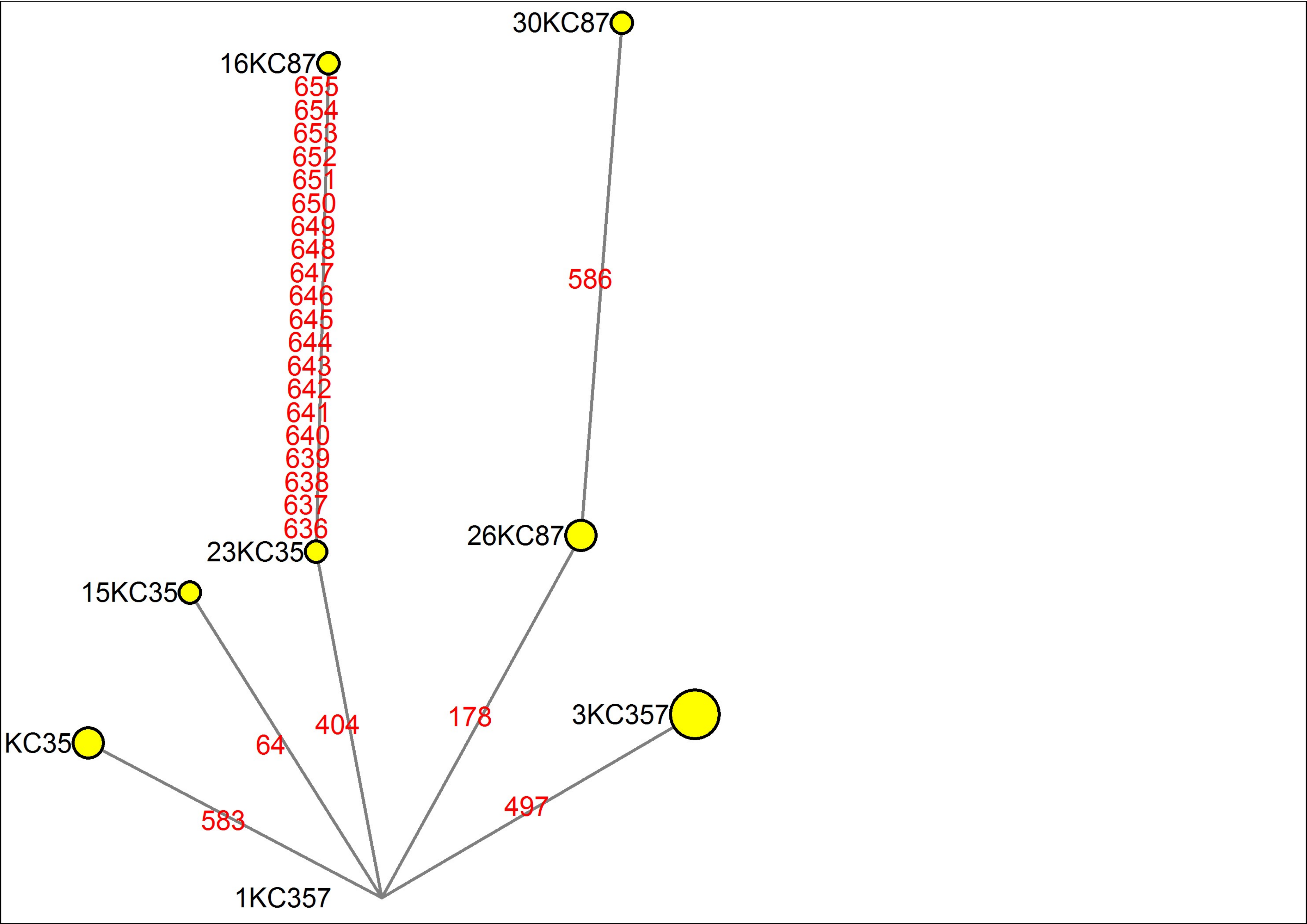

### MJN_case50.pdf

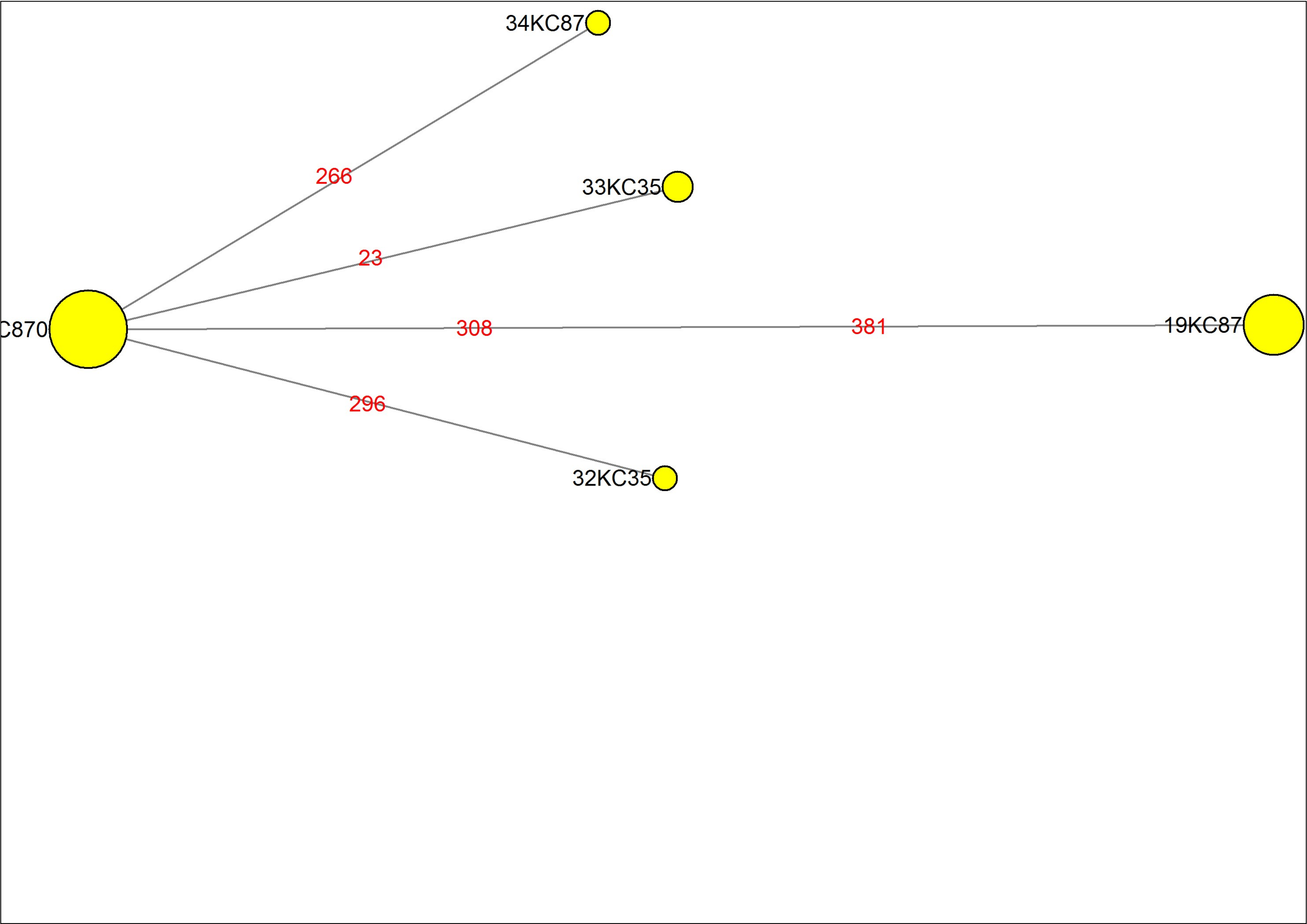

### MJN_case51.pdf

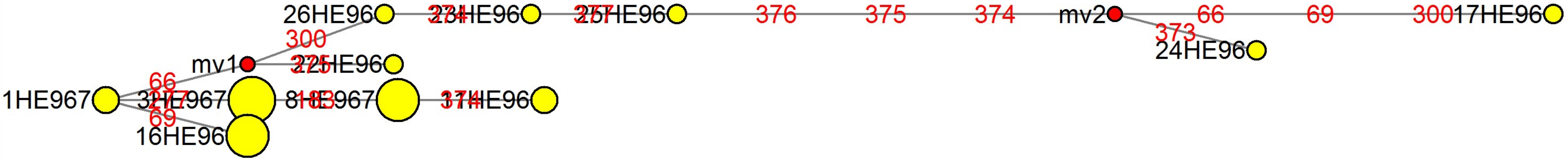

### MJN_case52.pdf

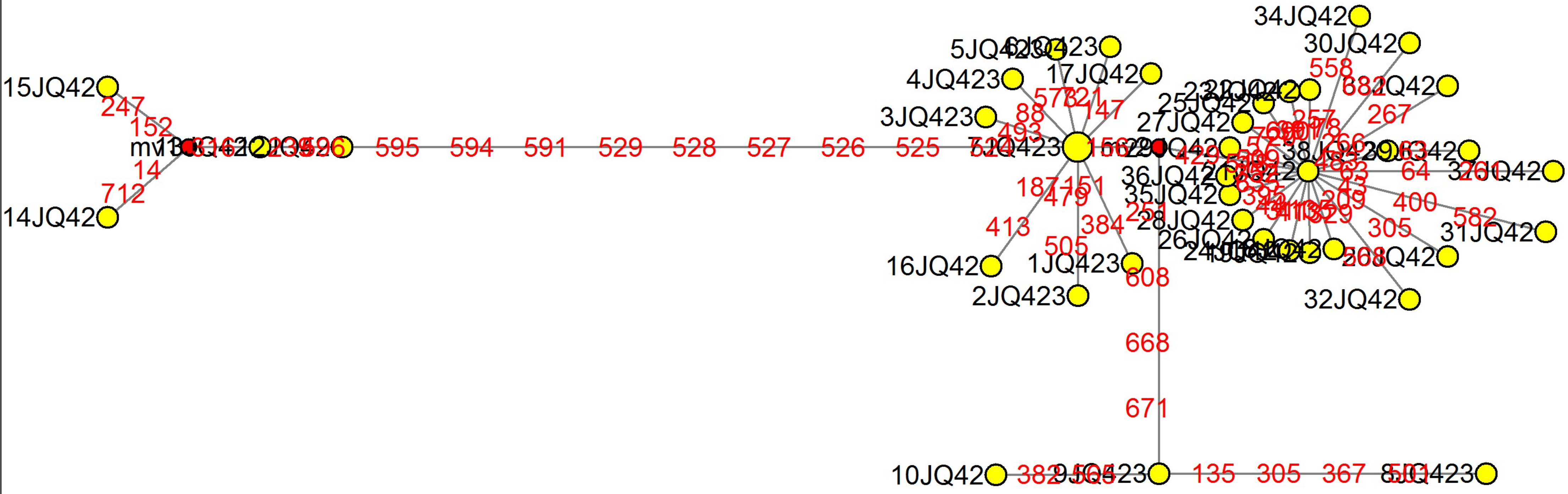

### MJN_case53.pdf

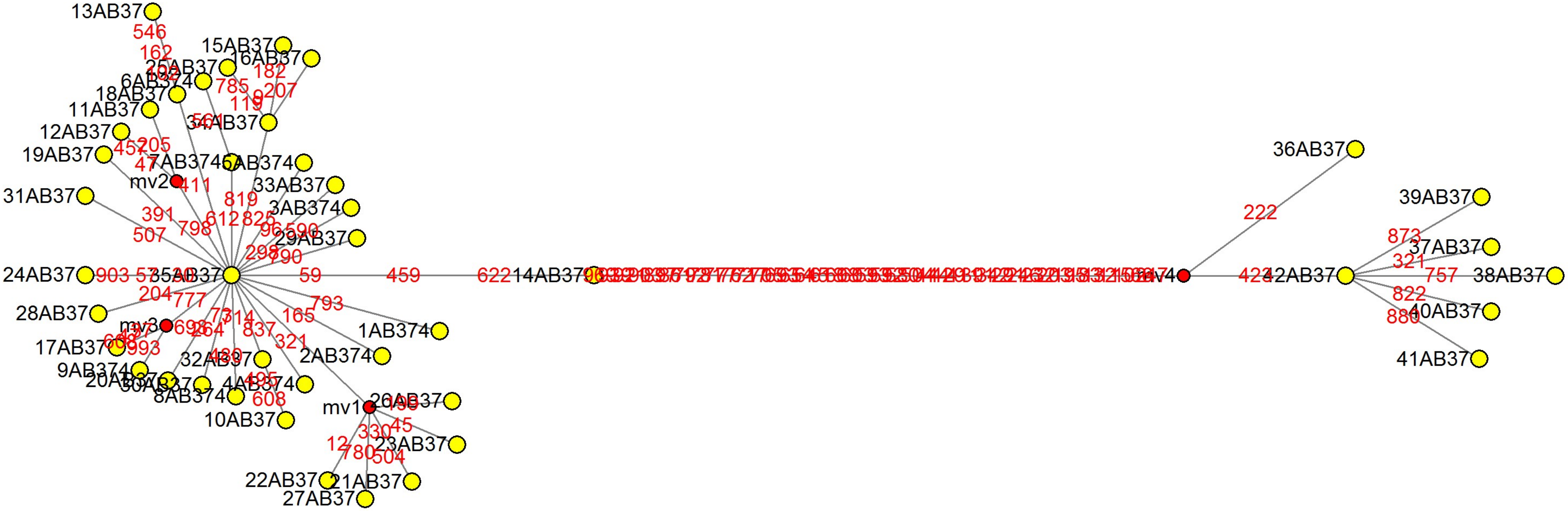

### MJN_case54.pdf

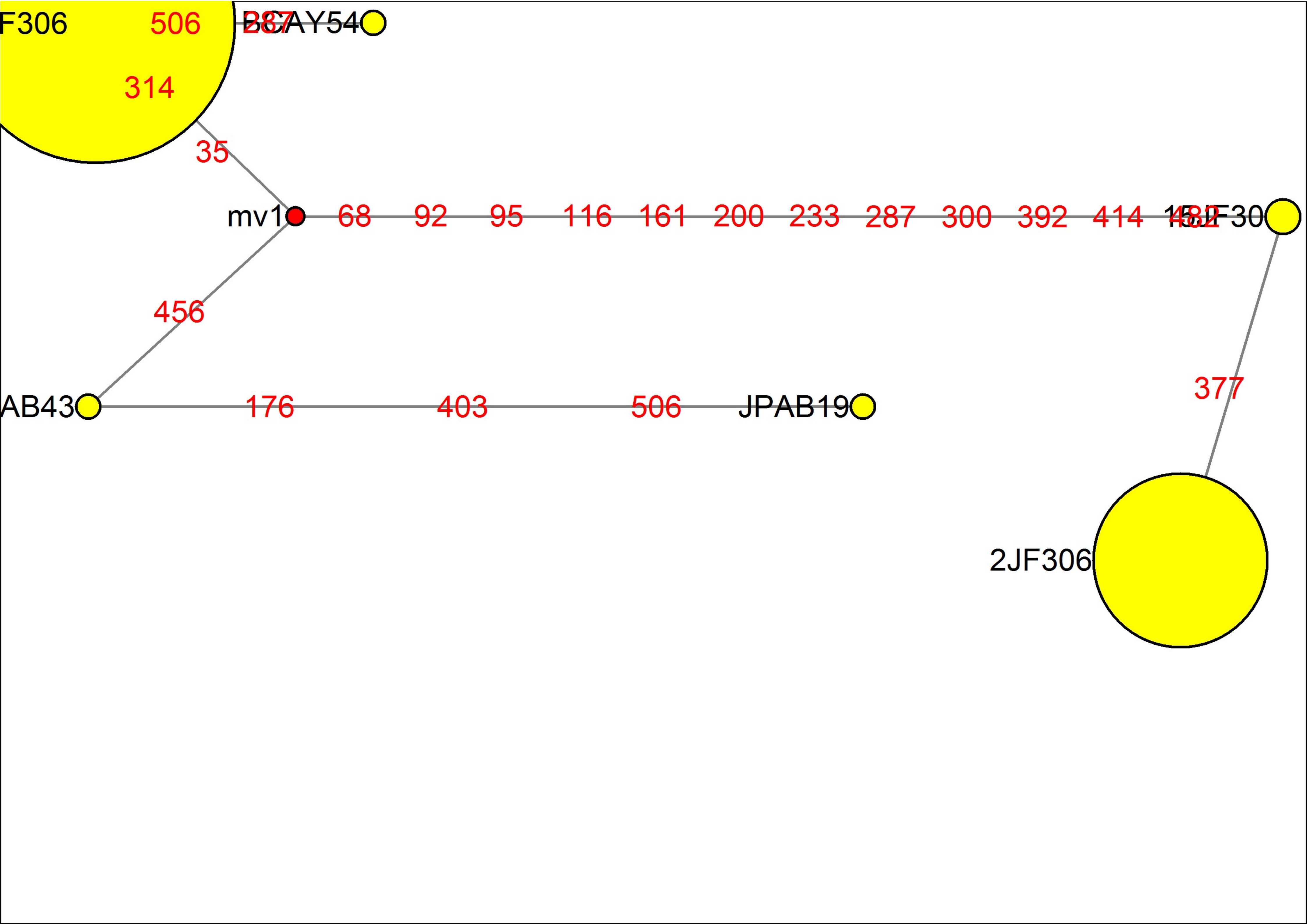

### MJN_case55.pdf

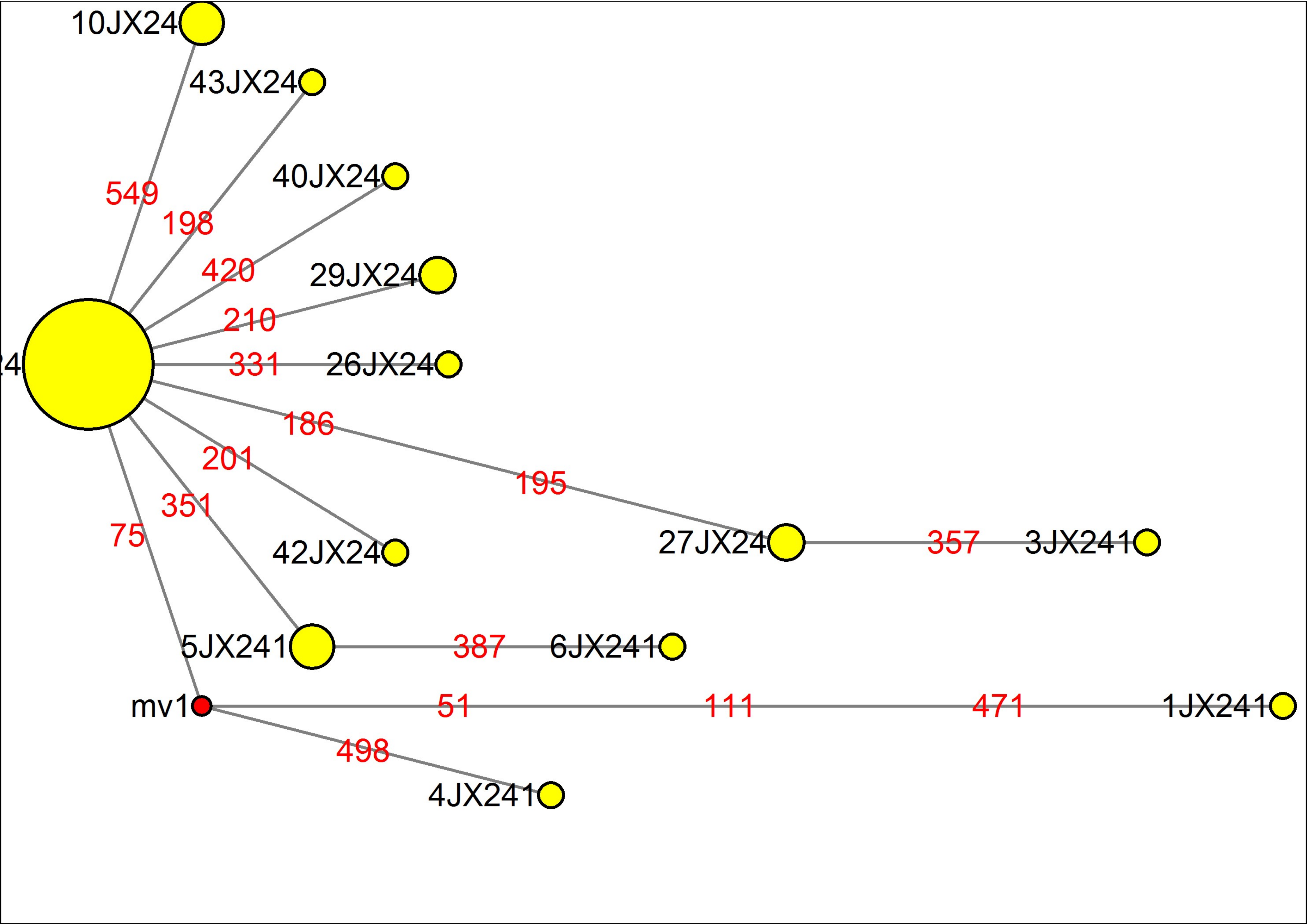

### MJN_case56.pdf

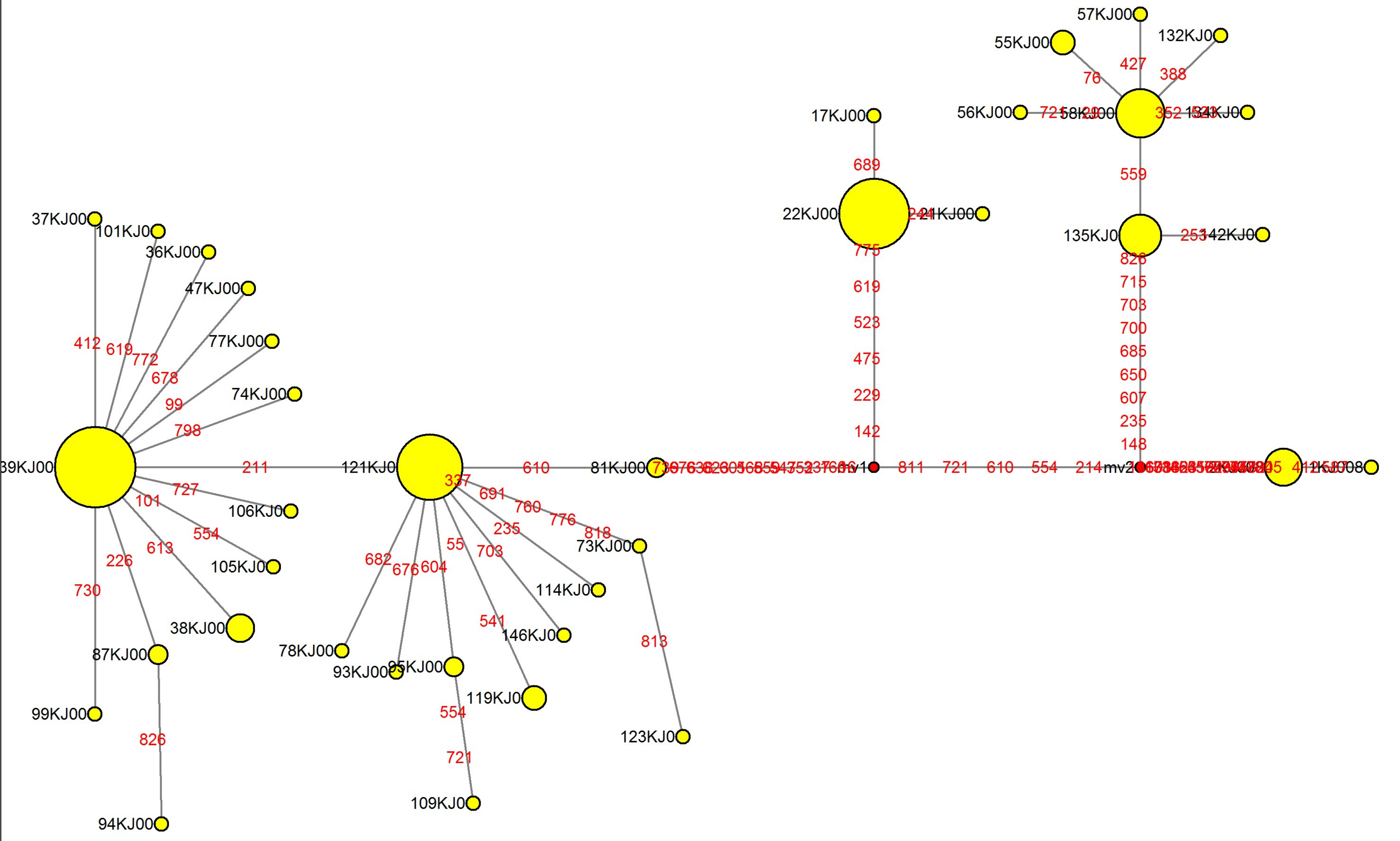

### MJN_case57.pdf

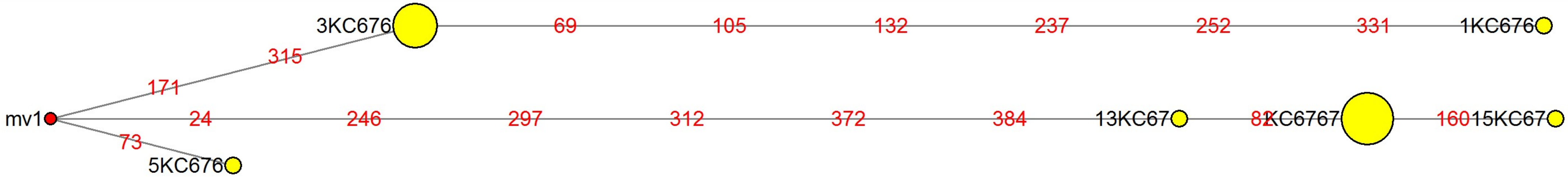

### MJN_case58.pdf

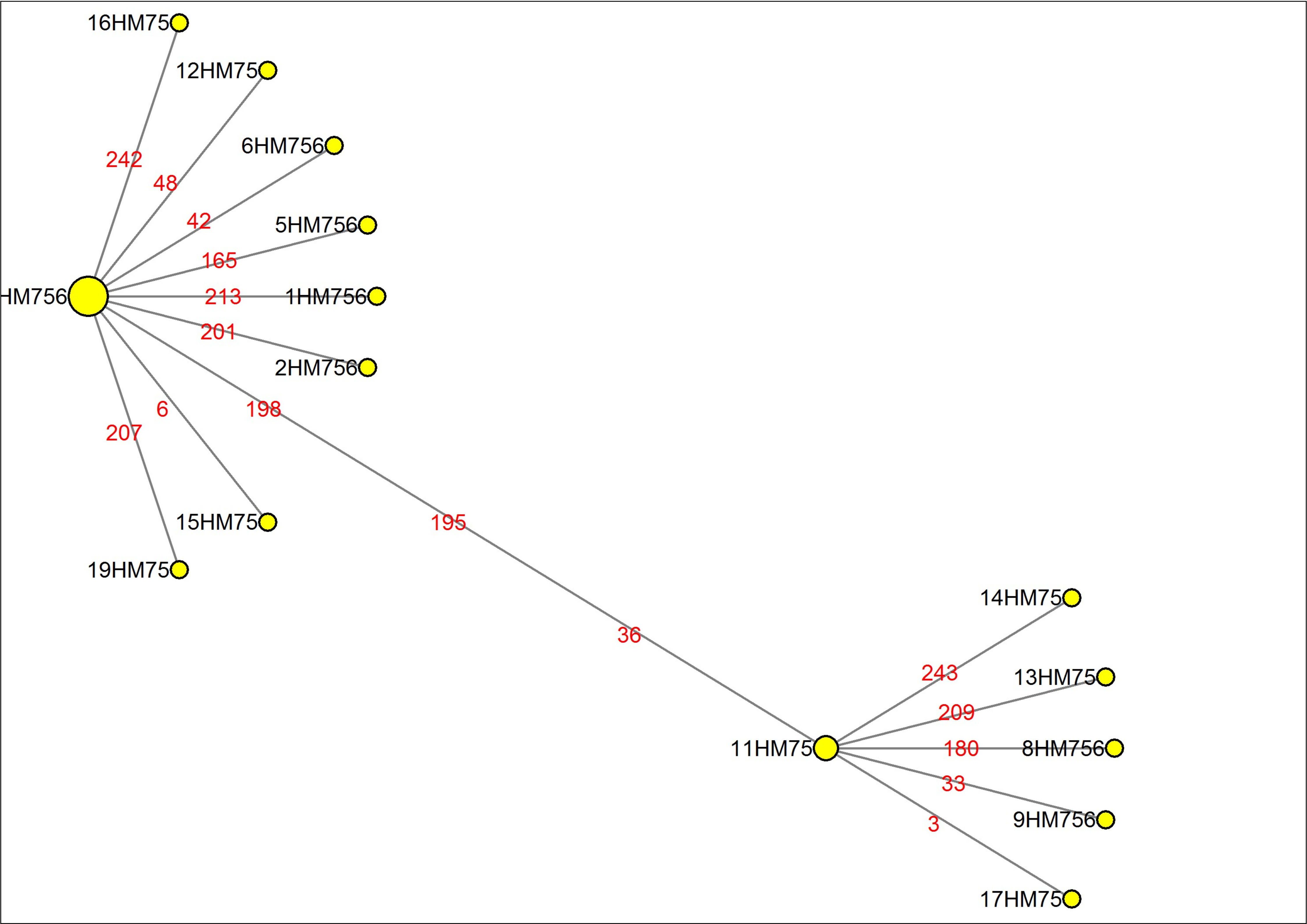

### MJN_case59.pdf

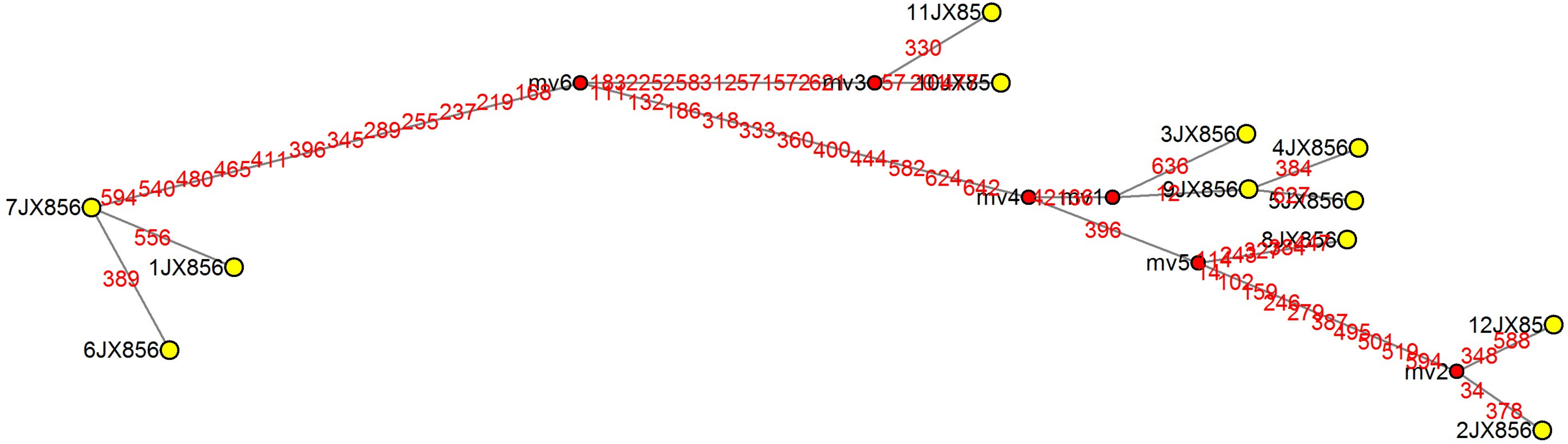

### MJN_case60.pdf

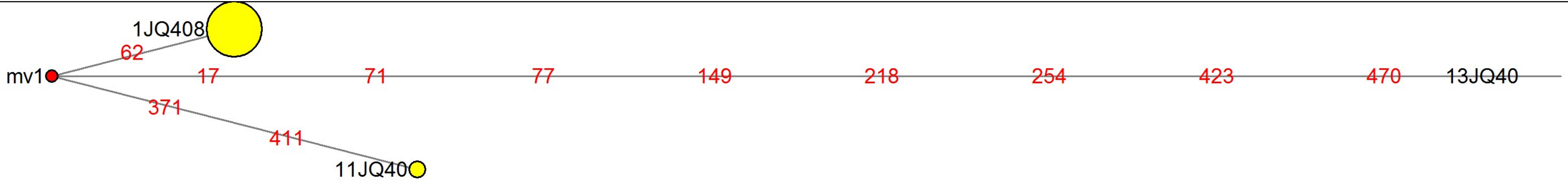

### MJN_case61.pdf

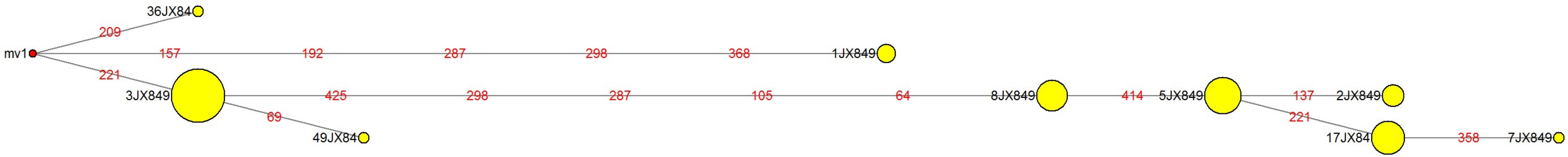

### MJN_case62.pdf

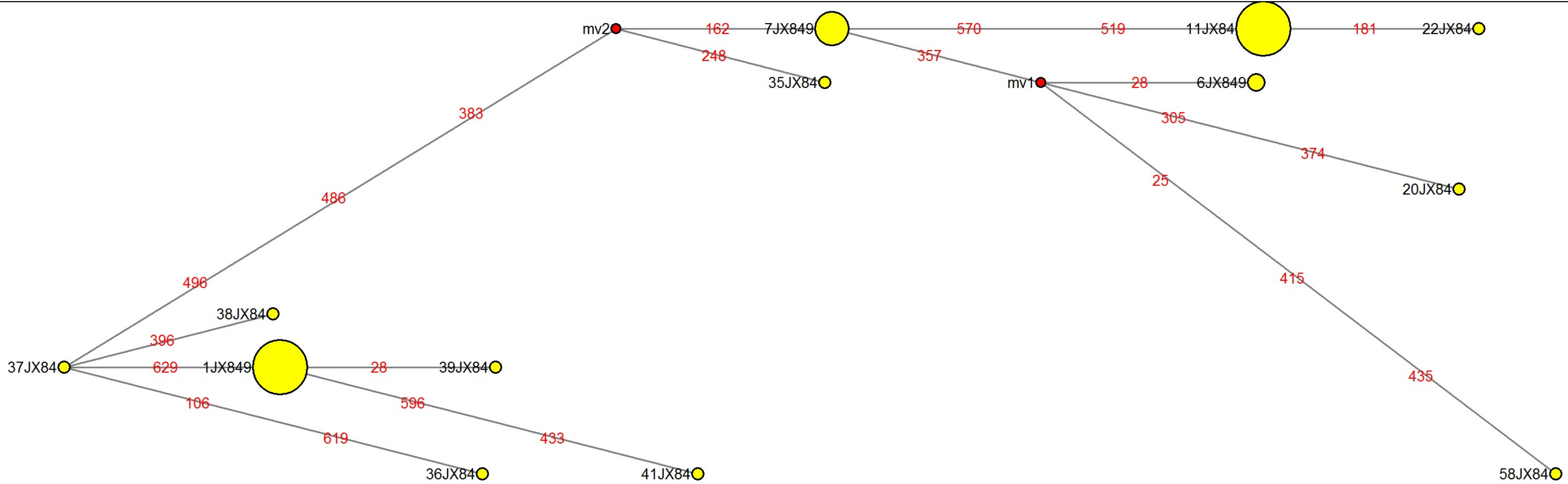

### MJN_case63.pdf

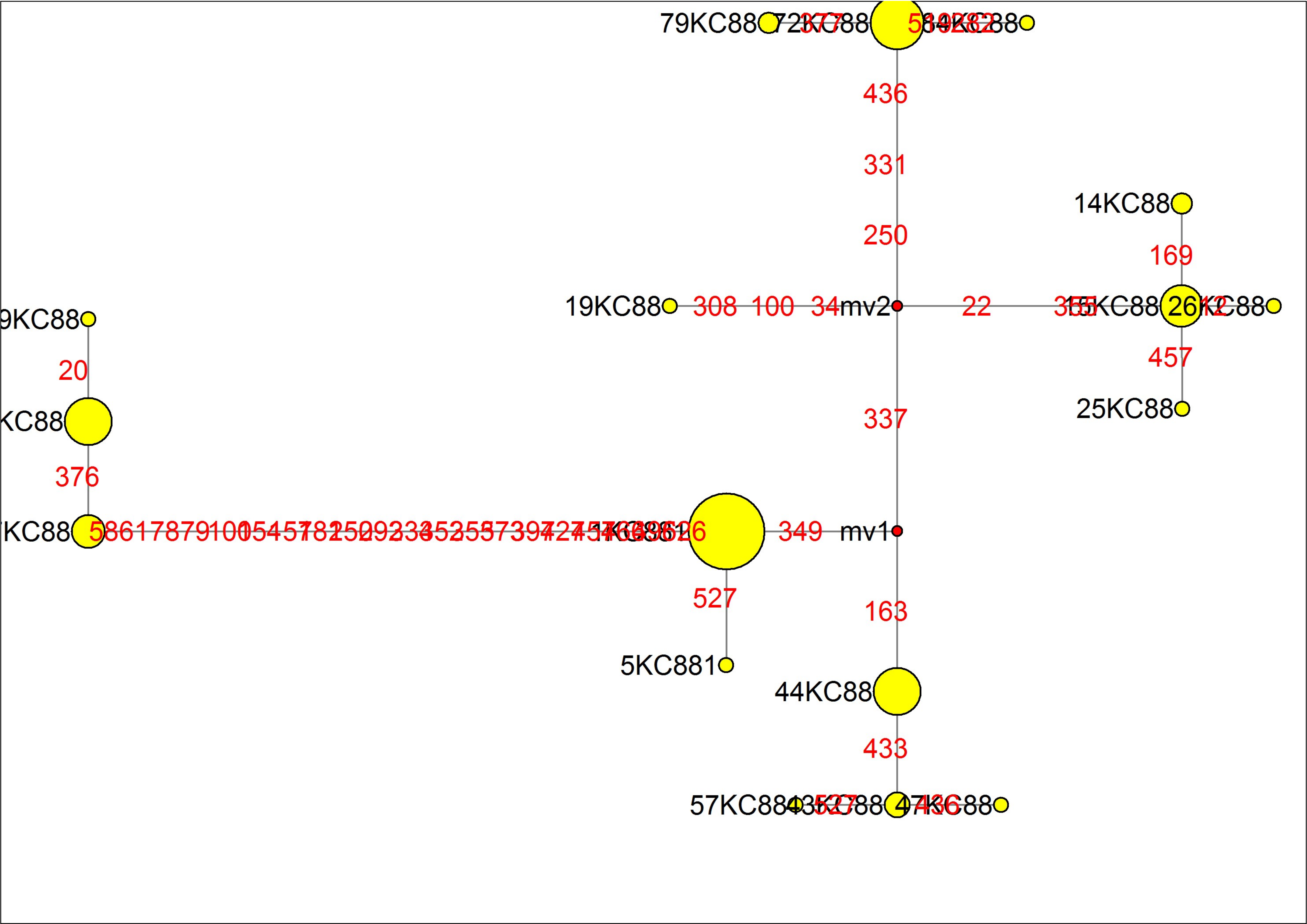

### MJN_case64.pdf

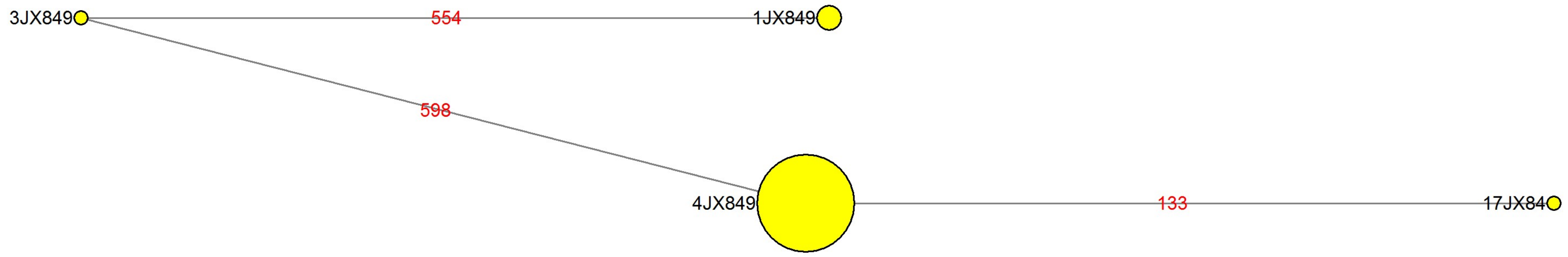

### MJN_case65.pdf

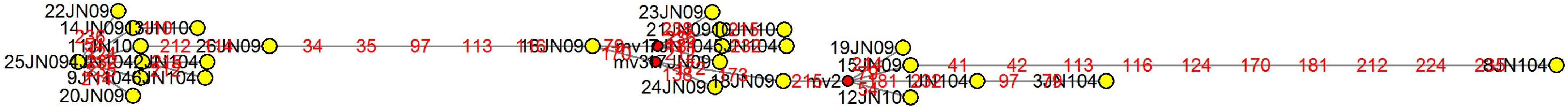

### MJN_case66.pdf

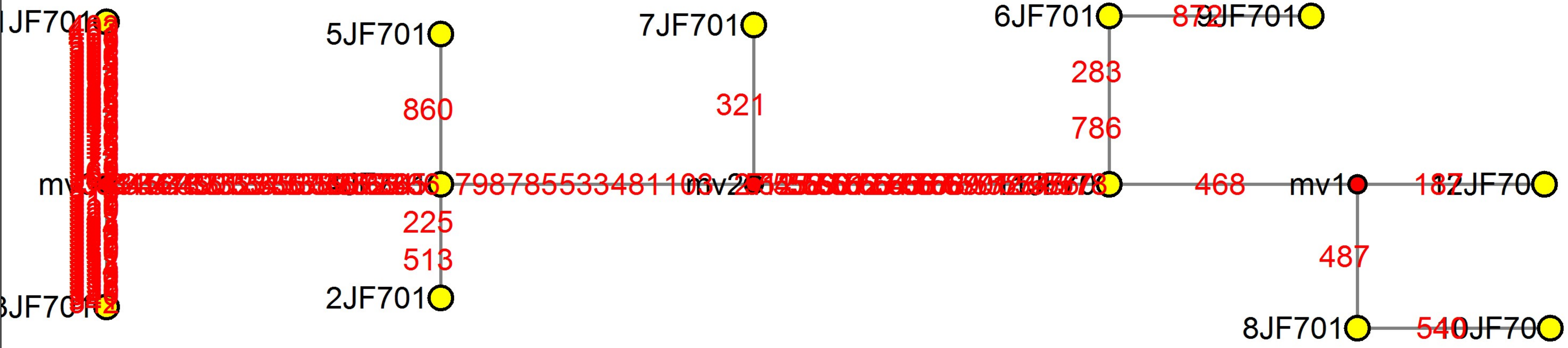

### MJN_case67.pdf

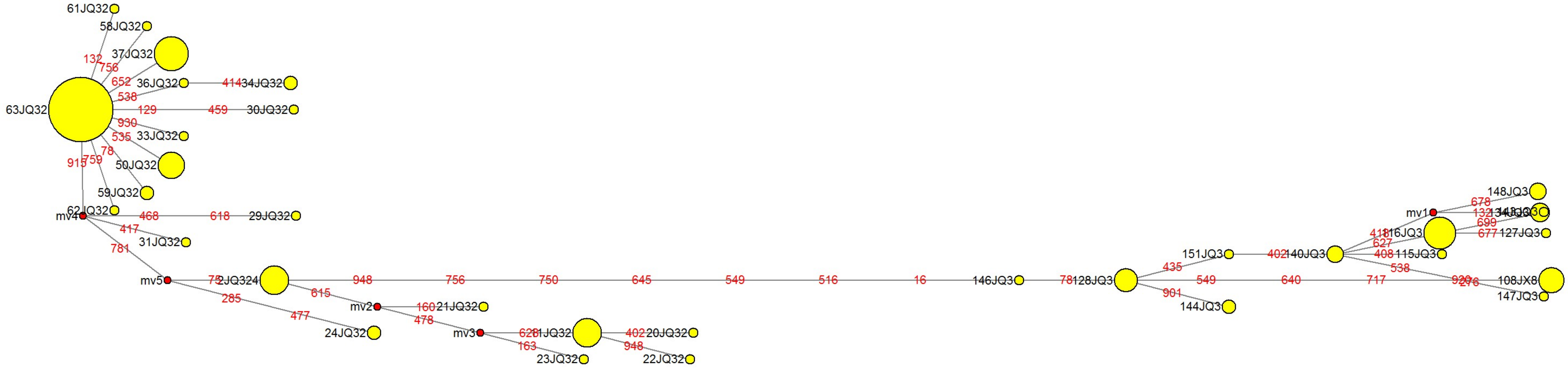

### MJN_case68.pdf

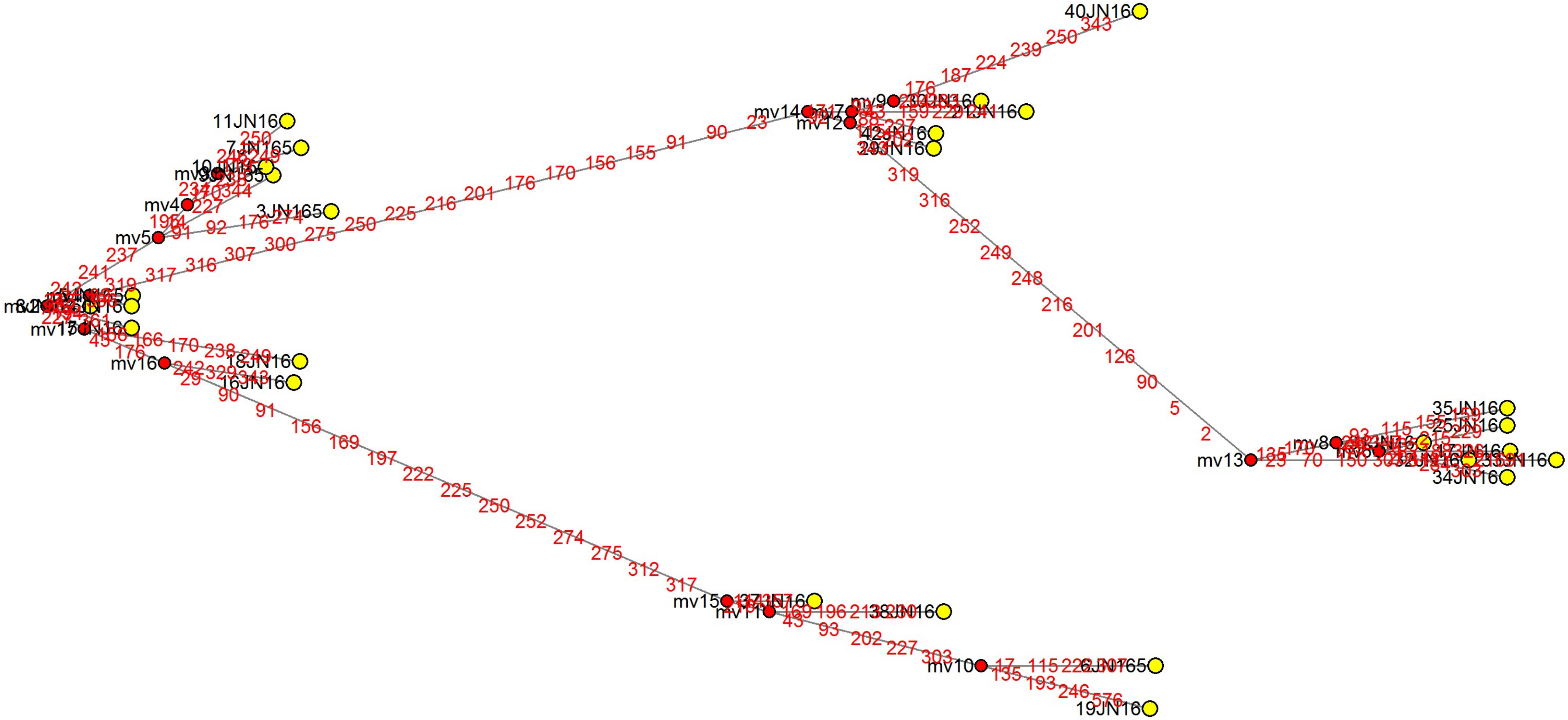

### MJN_case69.pdf

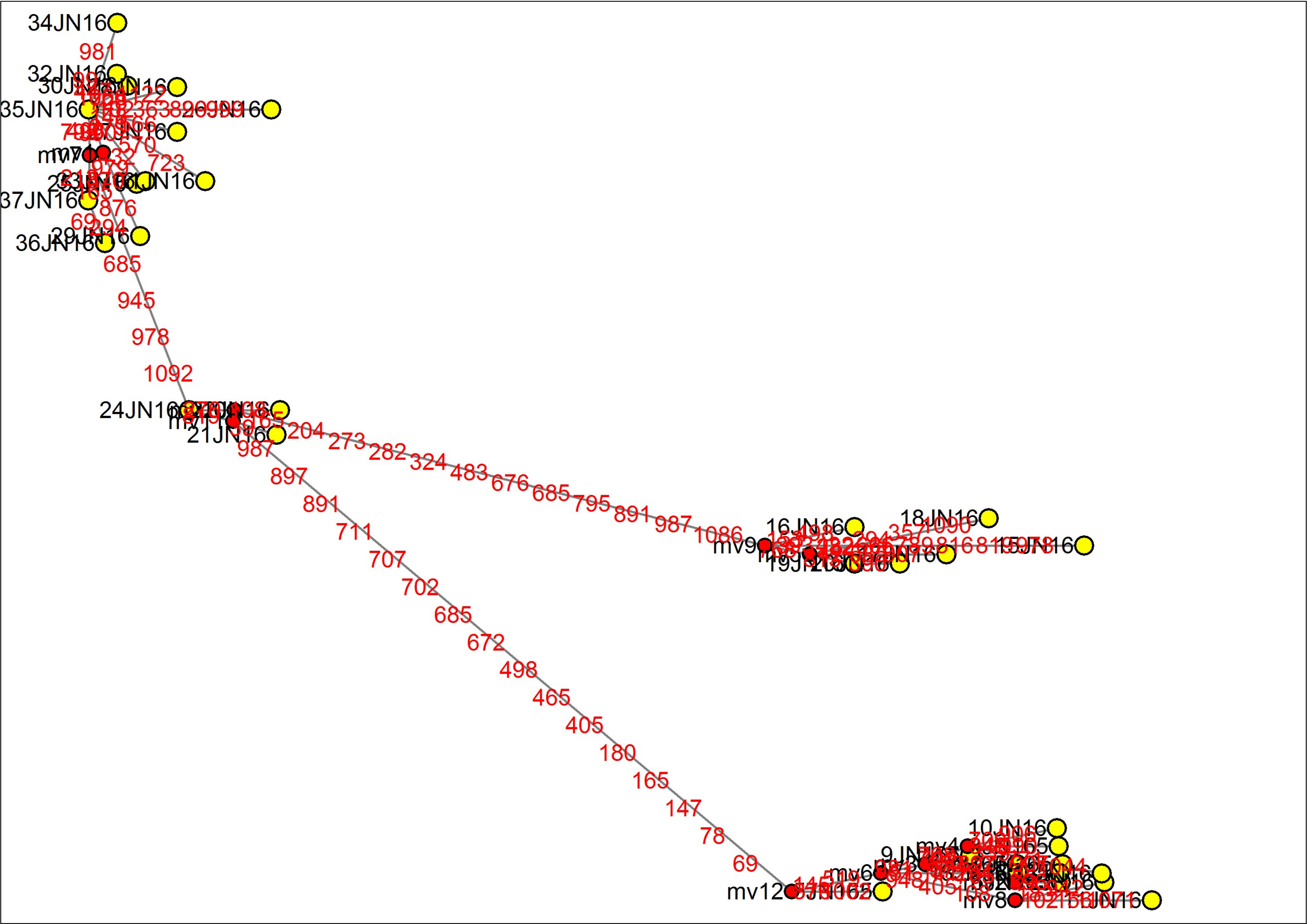

### MJN_case70.pdf

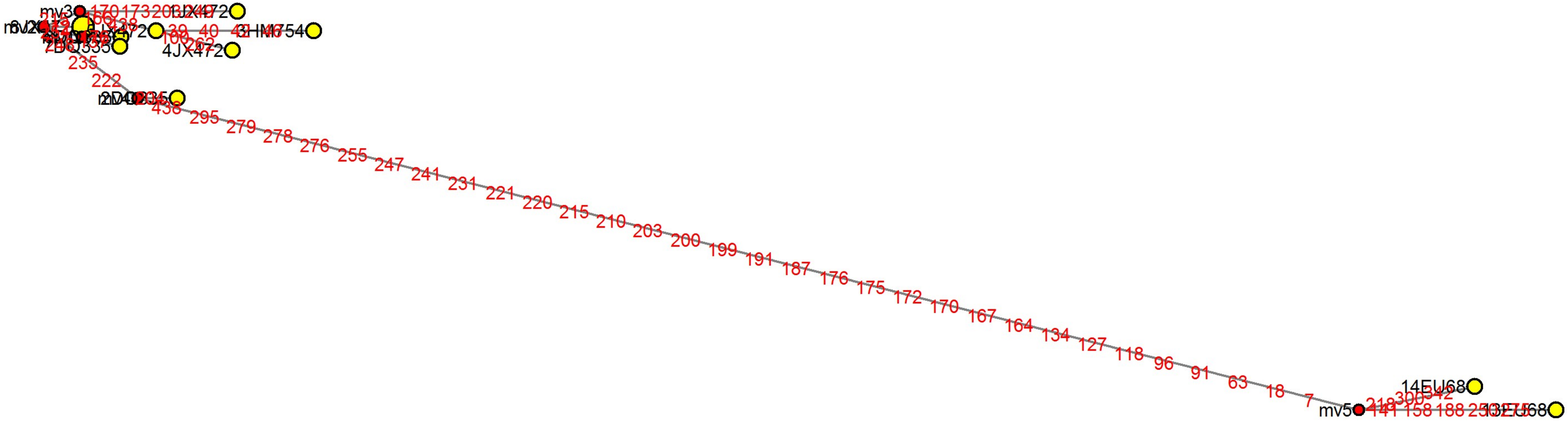

### MJN_case71.pdf

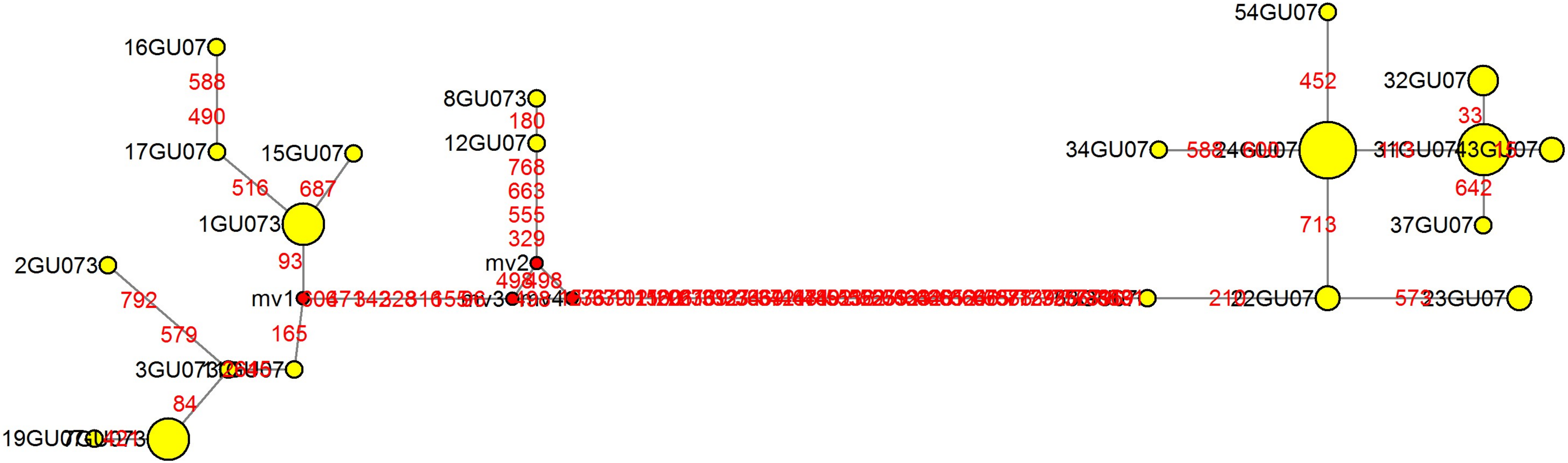

### MJN_case72.pdf

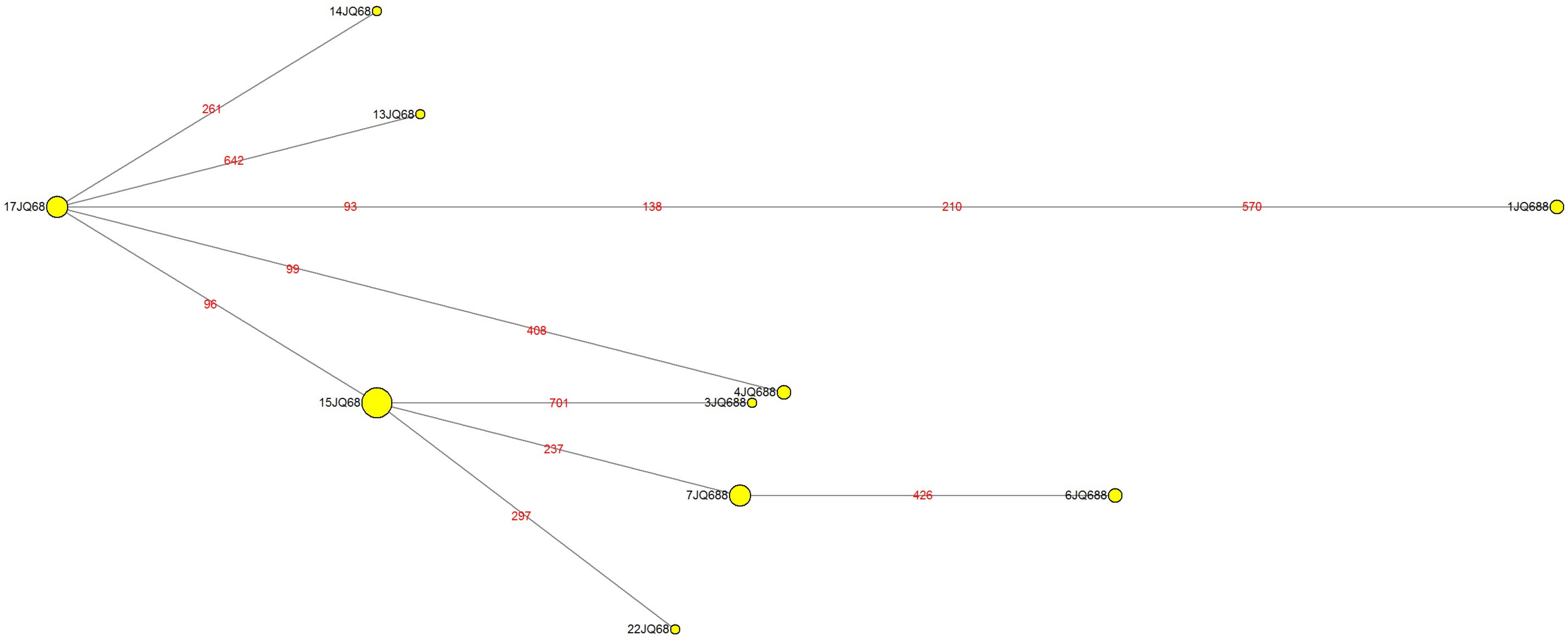

### MJN_case73.pdf

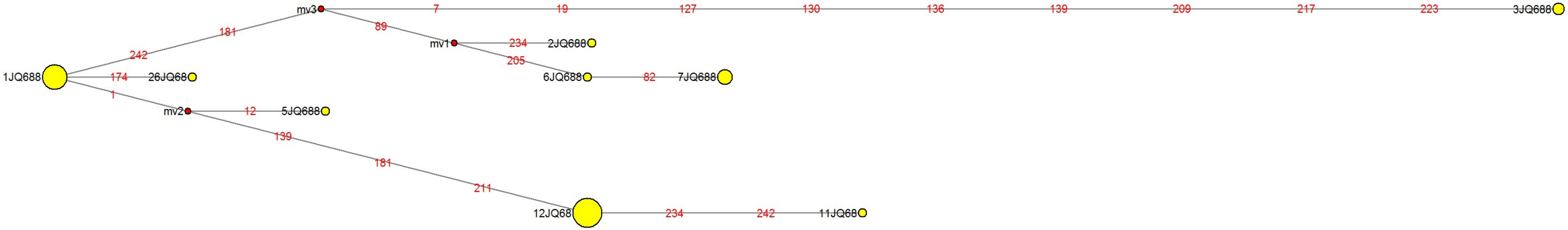

### MJN_case74.pdf

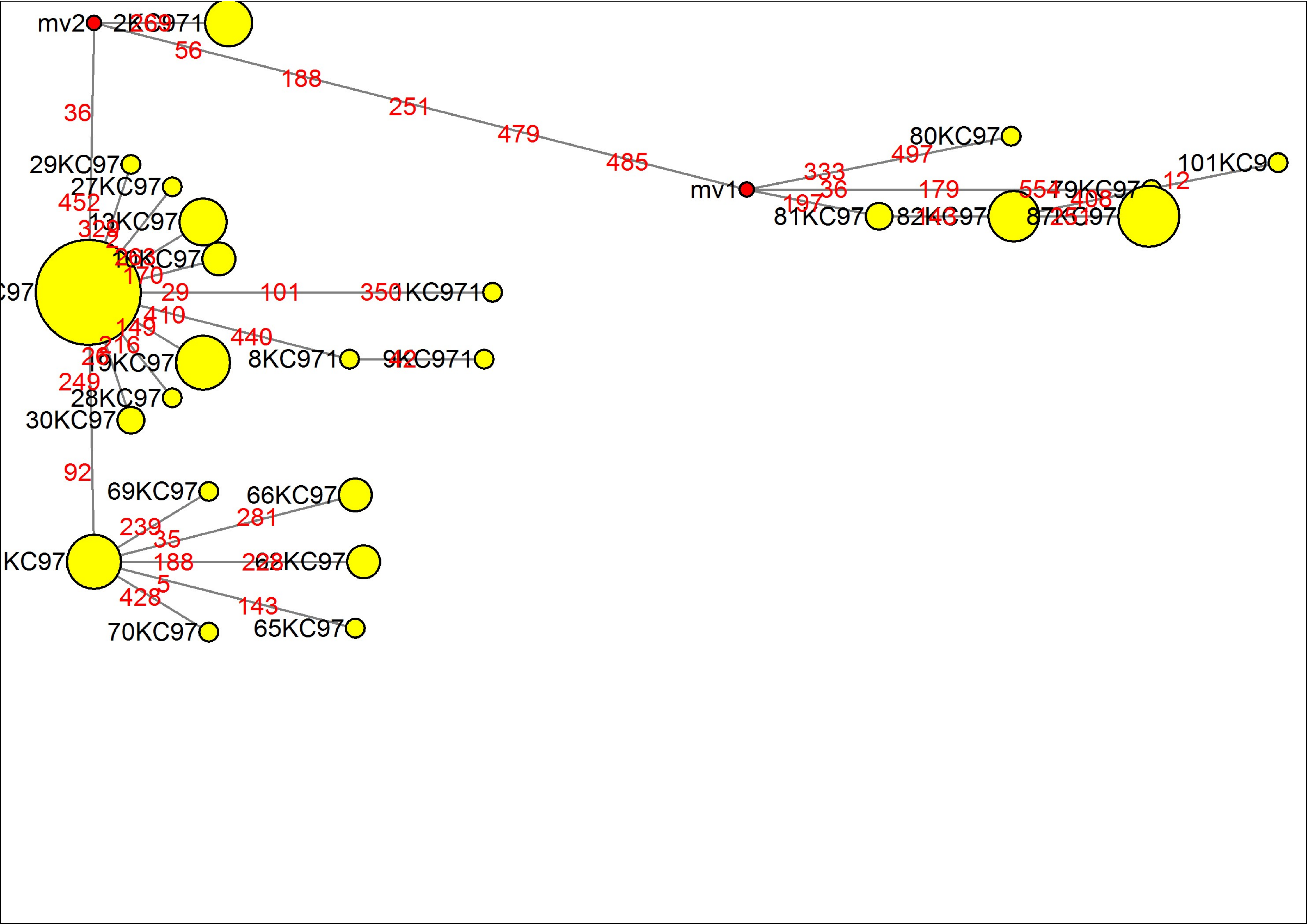

### MJN_case75.pdf

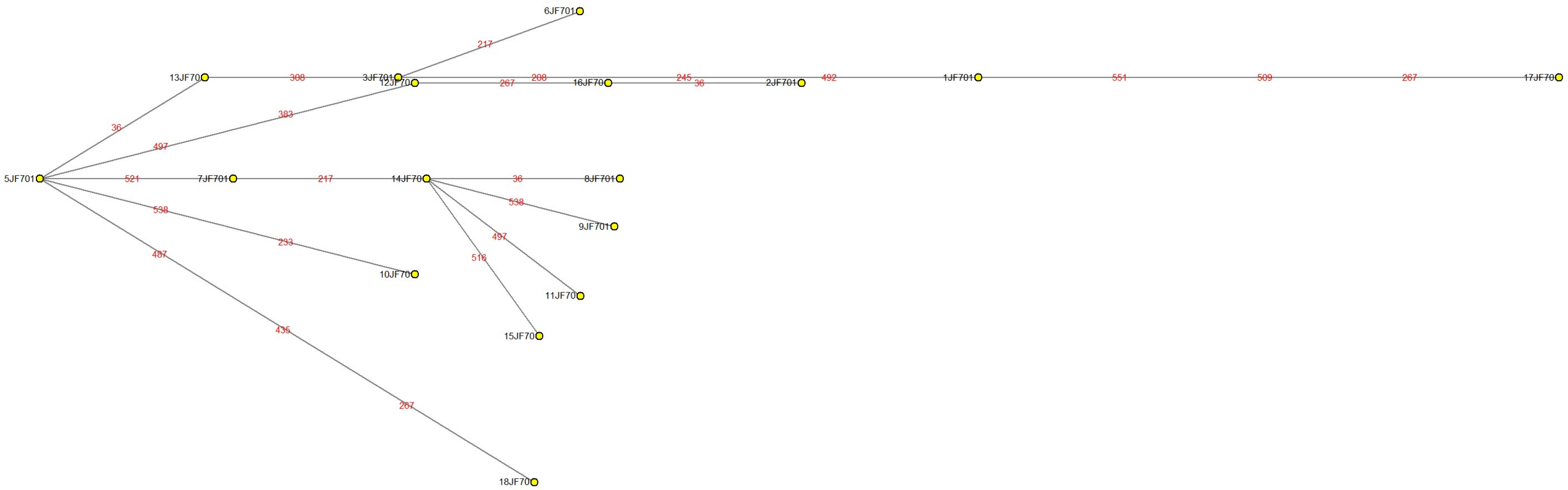

### MJN_case76.pdf

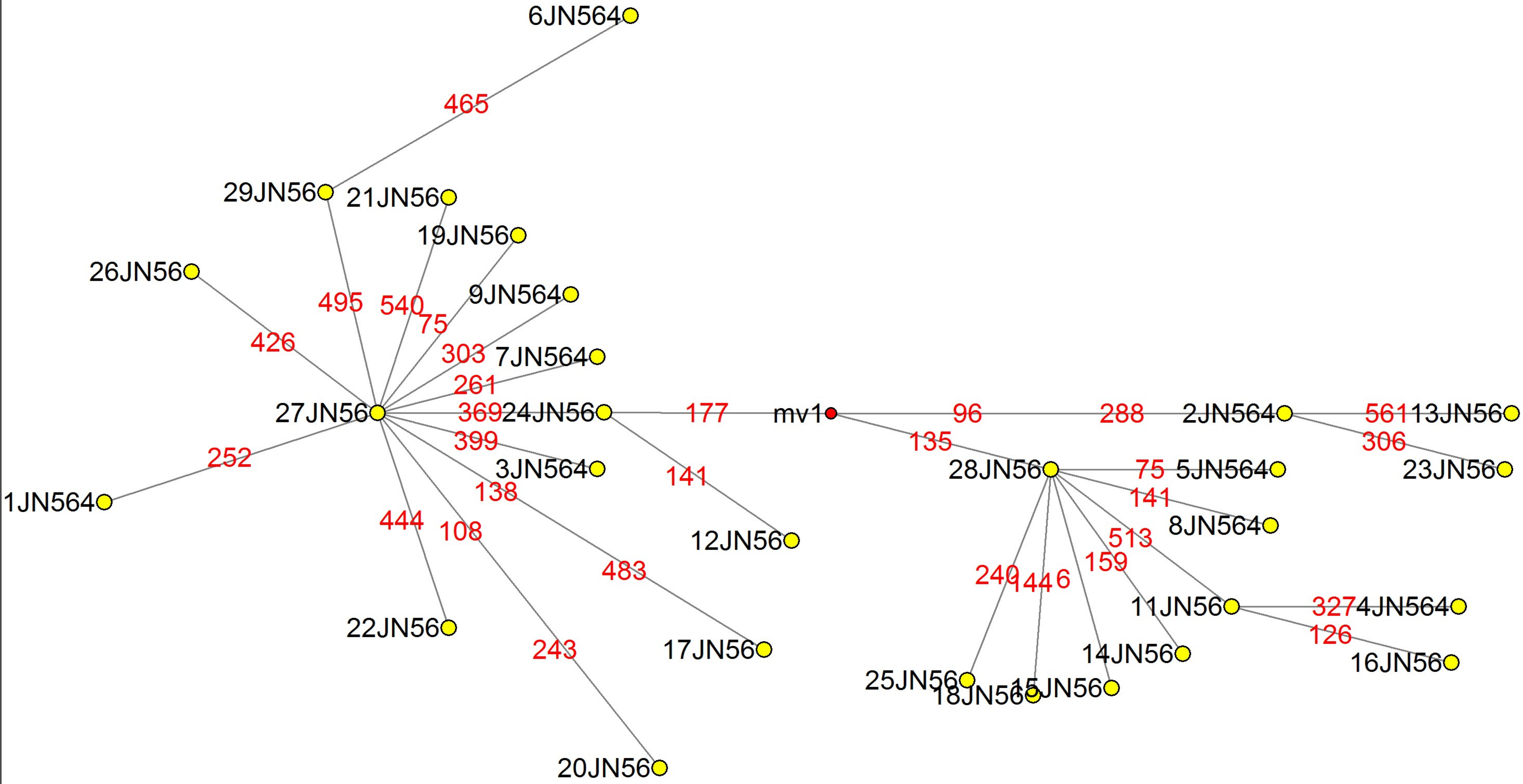

### MJN_case77.pdf

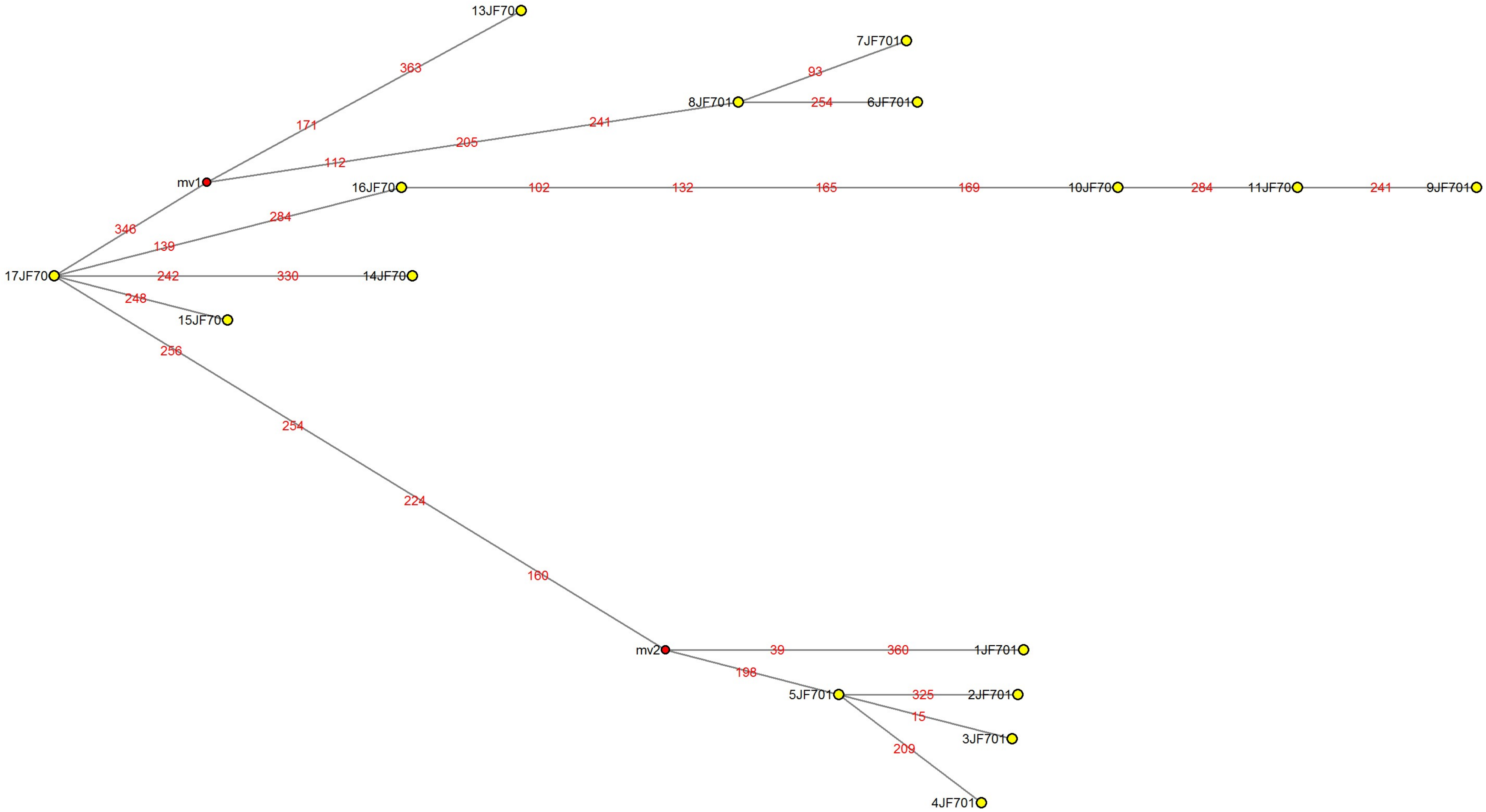

### MJN_case78.pdf

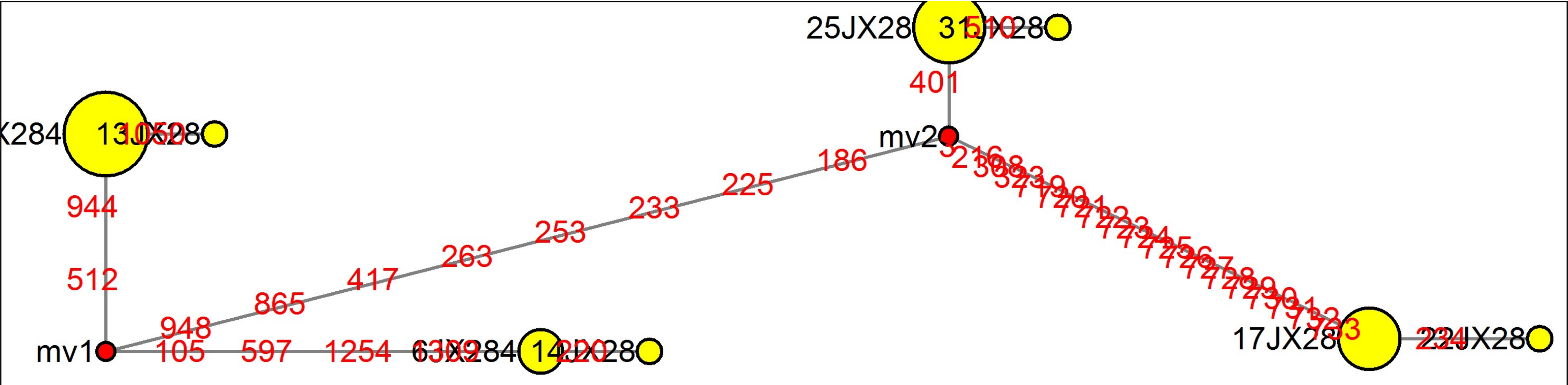

### MJN_case79.pdf

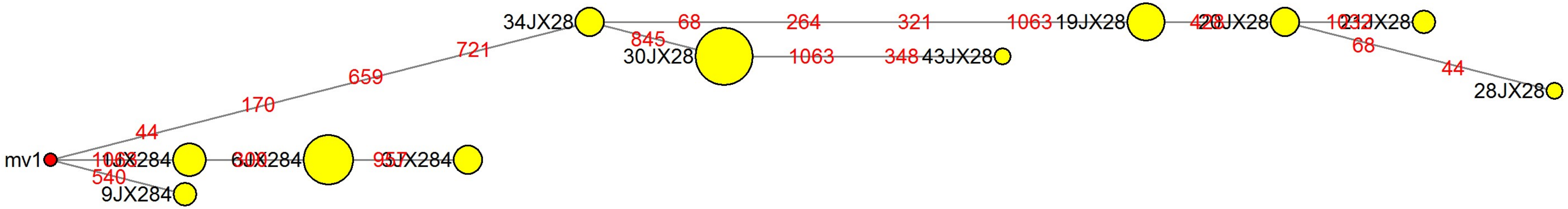

### MJN_case80.pdf

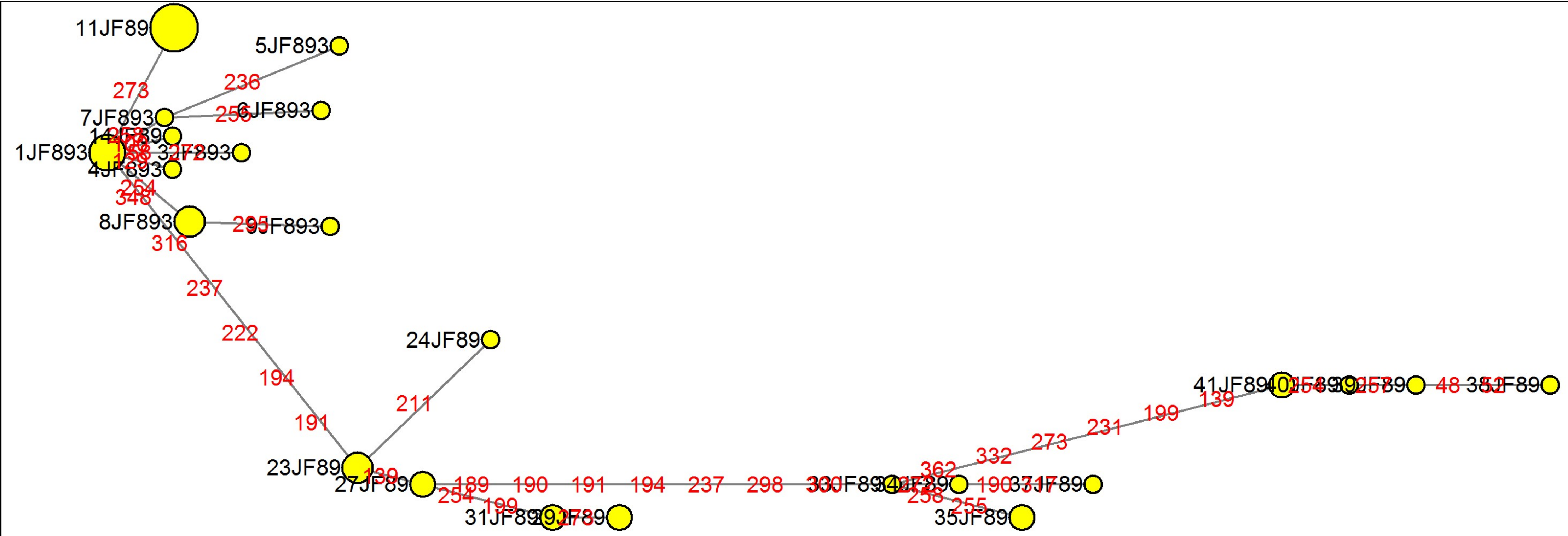

### MJN_case81.pdf

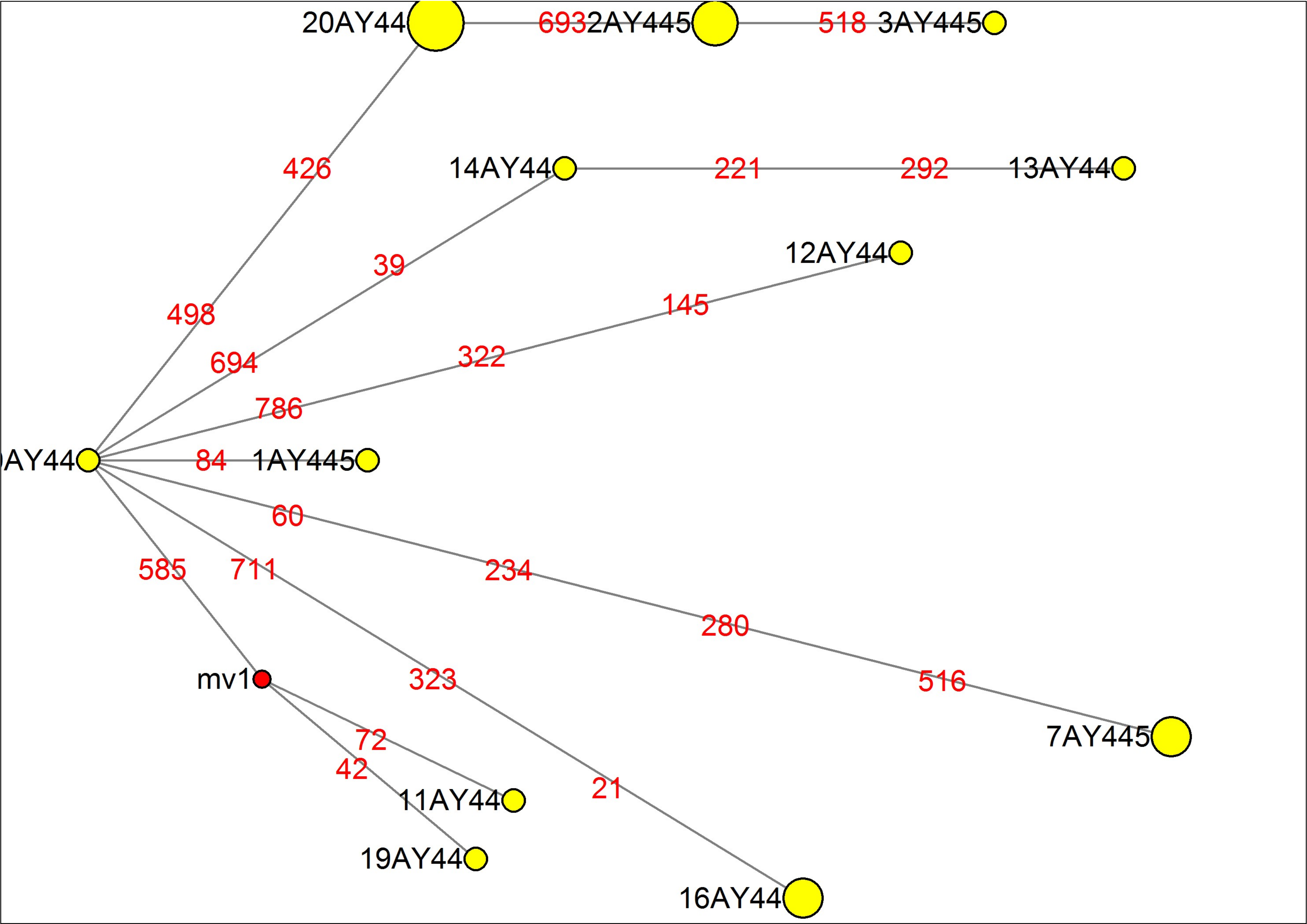

### MJN_case82.pdf

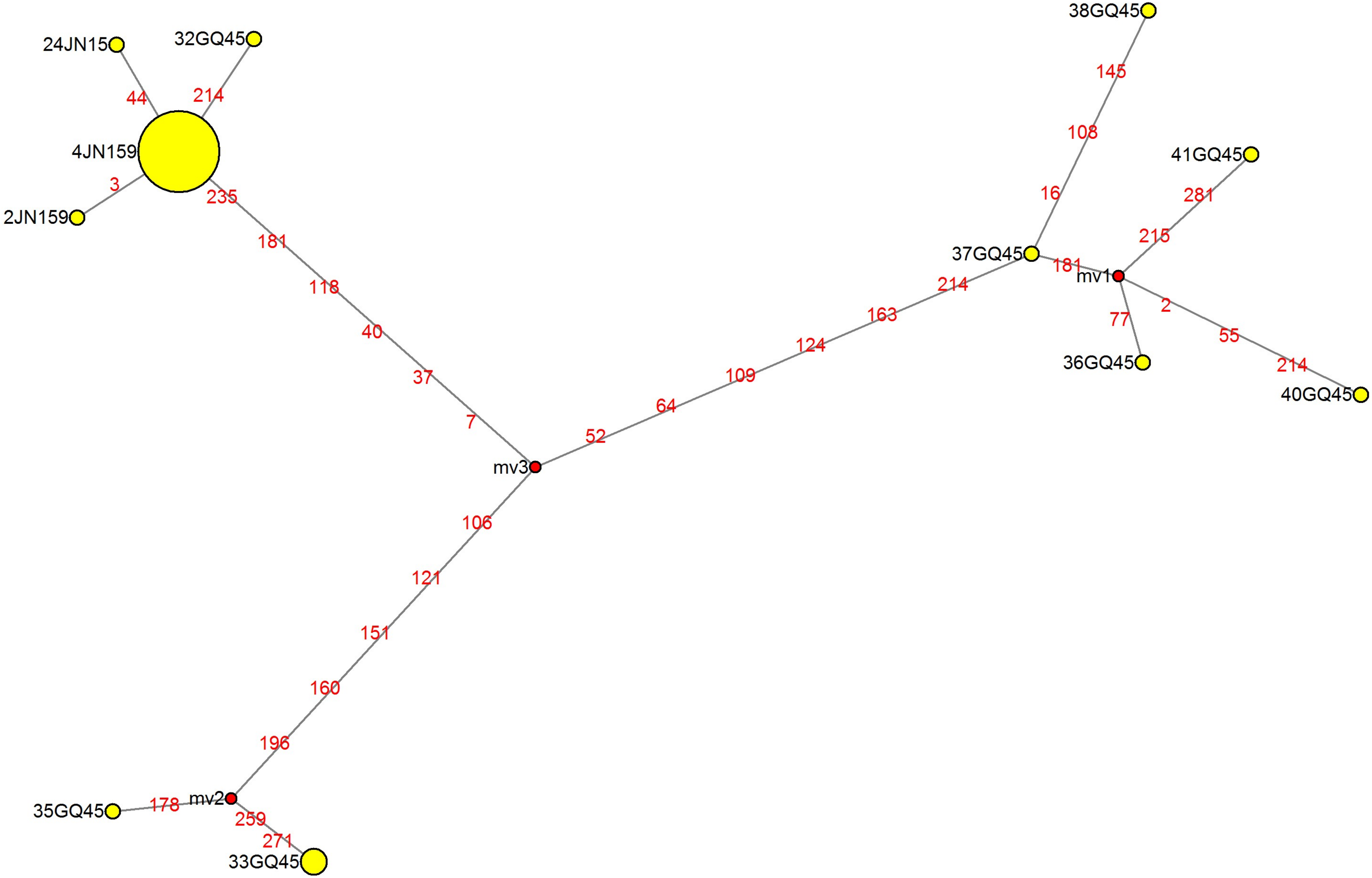

### MJN_case83.pdf

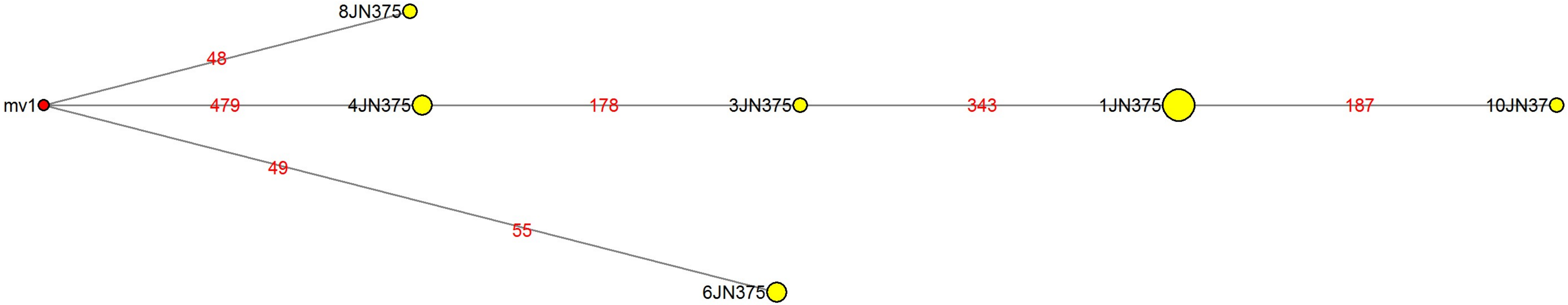

### MJN_case84.pdf

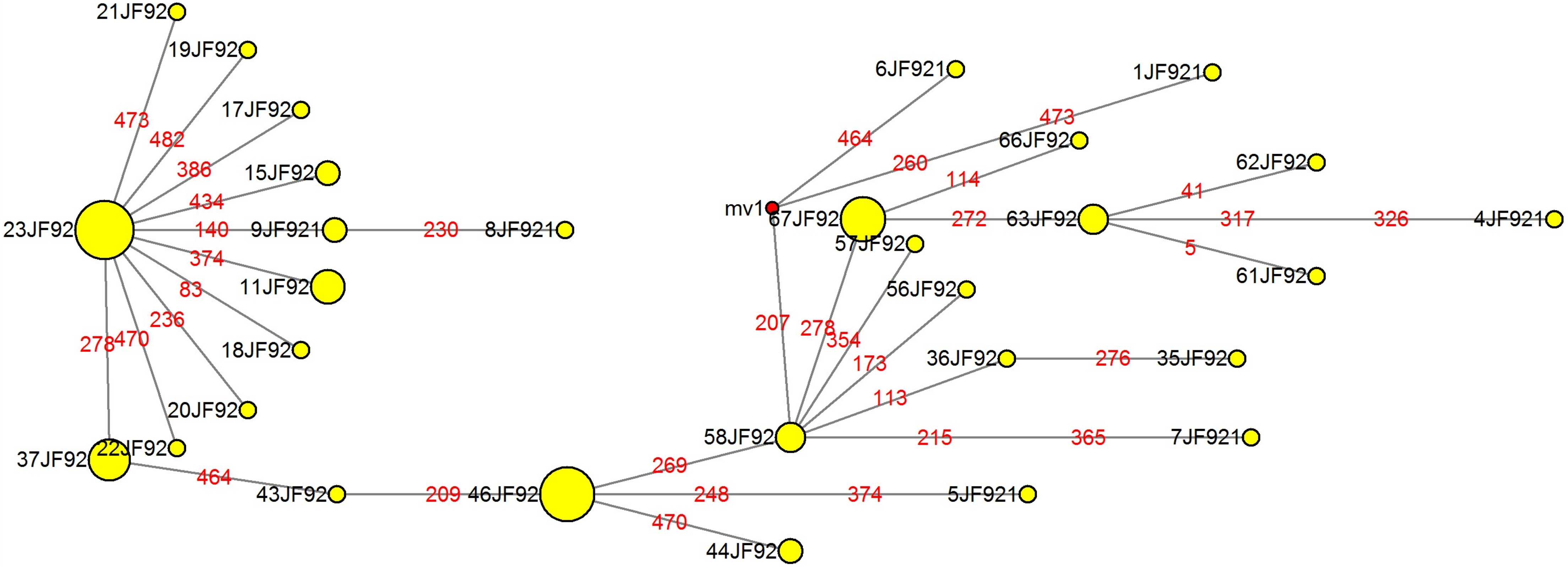

### MJN_case85.pdf

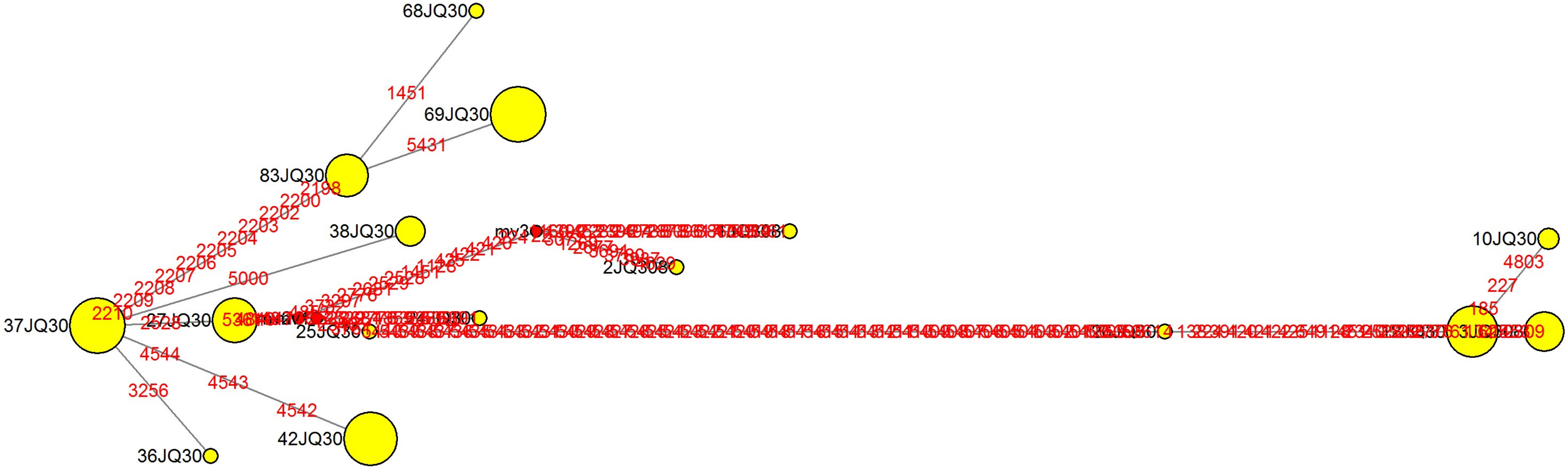
